# Treatment of STEC infection via CRISPR-Cas targeted cleavage of the Shiga toxin gene in animal models

**DOI:** 10.1101/2025.02.28.640725

**Authors:** Matthieu Galtier, Antonina Krawczyk, Fabien J. Fuche, Loïc H Charpenay, Igor Stzepourginski, Simone Pignotti, Marion Arraou, Rémi Terrasse, Andreas K Brödel, Chloé Poquet, Gautier Prevot, Dalila Spadoni, Benjamin Buhot, Kristin Munch, Jan Havránek, Pablo Cárdenas Ramírez, Marie Rouquette, Antoine Decrulle, Olivier Kerbarh, Erica Lieberman, Camila Bramorski, Aurélie Grienenberger, Edith M Hessel, Giuseppina Salzano, Daniel J. Garry, Aymeric Leveau, Xavier Duportet, David Bikard, Jesus Fernandez-Rodriguez

**Author notes:** **Corresponding authors**, Xavier Duportet, David Bikard and Jesus Fernandez-Rodriguez. These authors contributed equally to this work (Matthieu Galtier, Antonina Krawczyk). These authors contributed equally to this work (Fabien J. Fuche, Loïc H Charpenay).

## Abstract

*Escherichia coli* is a ubiquitous gut commensal but also an opportunistic pathogen responsible for severe intestinal and extra-intestinal infections. Shiga toxin-producing *E. coli* (STEC) pose a significant public health threat, particularly in children, where infections can lead to bloody diarrhea and progress to hemolytic uremic syndrome (HUS), a life-threatening condition with long-term complications. Antibiotics are contraindicated in STEC infections due to their potential to induce prophages carrying Shiga toxin (*stx)* genes, triggering toxin production. Here, we present a CRISPR-based antimicrobial strategy that selectively targets and eliminates O157 STEC clinical isolates while preventing toxin release. We designed a Cas12 nuclease to cleave >99% of all *stx* variants found in O157 strains, leading to bacterial killing and suppression of toxin production. To enable targeted delivery, we engineered a bacteriophage-derived capsid to specifically transfer a non-replicative DNA payload to *E. coli* O157, preventing its dissemination. In a mouse STEC colonization model, our therapeutic candidate, EB003, reduced bacterial burden by a factor of 3×10^3^. In an infant rabbit disease model, EB003 mitigated clinical symptoms, abrogated stx-mediated toxicity, and accelerated epithelial repair at therapeutically relevant doses. These findings demonstrate the potential of CRISPR-based antimicrobials for treating STEC infections and support further clinical development of EB003 as a precision therapeutic against antibiotic-refractory bacterial pathogens.

## INTRODUCTION

Shiga toxin (Stx)-producing *E. coli* strains (STEC), also known as enterohemorrhagic *E. coli* (EHEC), are an important cause of foodborne disease responsible for 2,8 million estimated acute illnesses annually worldwide (*1*). STEC affects primarily children with symptoms ranging from mild non-bloody to bloody diarrhea occurring in 70-90% of cases, haemolytic uremic syndrome (HUS) occurring in 10-15% of cases (*2*) to end-stage renal disease and death in 3-5% of the cases (*3*).

STEC strains produce two distinct families of toxins, Shiga toxin 1 (Stx1) and Shiga toxin 2 (Stx2), with Stx2 associated with the vast majority of HUS (*4*). These toxins bind a glycolipid globotriaosylceramide receptor (Gb3) and enter the cytosol where they cleave a specific adenine residue in the 28S ribosomal RNA, leading to protein synthesis inhibition (*5, 6*). The *stx1* and *stx*2 genes are carried by bacteriophages which likely play an important role in the spread of the *stx* genes (*7–9*), with *E. coli* O157:H7 being the most common serotype associated with human disease (*10*).

The *stx* genes are located within the late operons of Stx-encoding phages. The expression and release of Stx2 depends on the induction of the phage lytic cycle (*11, 12*). This likely explains why antibiotic treatment of STEC infection has been associated with increased rates of HUS (*13*). Indeed, many antibiotics are known to induce phages through the SOS response, including antibiotics commonly used in the treatment of diarrhea (*14, 15*). As a result, they are not recommended for the treatment of STEC infections (*16, 17*) and diarrhea patients are typically only treated symptomatically (*18, 19*). There is thus a need for novel strategies able to eradicate STEC strains while preventing toxin release.

Phage therapy has been proposed as an alternative strategy. Phage-based products are already approved by the FDA to decontaminate food products (*20*), and lytic phages have shown some efficacy in the treatment of mice infected with an O157 STEC strain (*21, 22*). However, phage therapy does not directly inhibit toxin production. Rather than using natural phages, genetic engineering may offer interesting solutions to abrogate toxin production and release. Hsu and colleagues showed how engineered temperate phages could propagate through a bacterial community and block the expression of *stx* genes (*23*).

In recent years, phage-based delivery of CRISPR-Cas systems has been proposed to enhance their lytic potential or achieve sequence-specific killing (*24–28*). Some studies have explored the use of CRISPR-Cas to target antibiotic resistance genes or the *eae* virulence factor in EHEC. Cas9 was delivered using an M13 capsid (*26*), and a type I CRISPR-Cas system was added to the genome of phage λ (*28*). However, these vectors are unsuitable for clinically relevant strains. M13 requires a conjugative pilus absent in many strains (*29*), while λ recognizes outer membrane porins frequently hidden behind the O-antigen or capsules (*30*). Engineering phage receptor-binding proteins is hence essential to target clinically relevant strains. For instance, the P4 capsid has been engineered to recognize an O157 strain (*31*). When armed with CRISPR-Cas9, these capsids were shown to kill an O157 STEC strain *in vitro*, but their broad therapeutic potential and targeting of diverse clinical isolates remains untested *in vivo*.

Here, we take a fundamentally different approach by engineering a delivery vector that recognizes the vast majority of O157 clinical isolates while ensuring a direct shutdown of toxin production. Previously, we successfully used a λ-derived particle with engineered receptor-binding proteins to deliver a base editor to *E. coli* colonizing the mouse gut (*32*). Building on this λ-based platform, we developed EB003, a rationally engineered antimicrobial that delivers a non-replicative CRISPR-Cas12 system to *E. coli* O157 clinical isolates. The CRISPR array is designed to target over 99% of *stx* gene variants, ensuring efficacy in clinical settings. Cas-mediated cleavage of the bacterial genome leads to rapid DNA degradation and cell death (*33–35*), while *stx* targeting immediately halts toxin expression and release.Thus, our system efficiently eliminates a broad range of O157 STEC isolates while preventing toxin production, addressing a key unmet medical need. In mouse and rabbit STEC models, EB003 significantly reduced bacterial load and alleviated symptoms, including suppression of toxin production. These results provide strong support for the further development of EB003 as a targeted therapeutic for EHEC infections.

## RESULTS

### An engineered λ bacteriophage capsid can efficiently deliver DNA to O157:H7 strains

Bacteriophage λ is a *Siphoviridae* that was isolated from the *E. coli* K12 strain. Its ability to deliver DNA to *E. coli* K12 is determined by two main components of the capsid. The side tail fiber (stf) gene encodes long appendages that promote adsorption of the phage to target bacteria through interaction with the OmpC outer membrane porin (*36, 37*), and the tail tip protein gpJ which recognises the LamB outer membrane porin (*38, 39*).

We previously demonstrated how the λ capsid could be engineered to carry chimeric forms of the gpJ protein that recognise OmpC from the model O157 strain EDL933 as a receptor rather than LamB (gpJ-1A2 or gpJ-A8) (*32*). This ensures efficient delivery of DNA to *E. coli* in the mouse gut environment, where it expresses OmpC reliably as opposed to LamB (*40, 41*). Our bioinformatic analyses showed that all 121 sequenced O157 strains in our collection isolated between 2015 and 2018 in the UK and France encode the same OmpC variant (Supplementary Methods).

In our previous study the Stf protein of λ was also engineered to recognise the LPS of different *E. coli* strains. O157 strains express a group 4 capsule, which masks surface structures that may serve as bacteriophage binding points (*42*).

To identify STF variants that would enable efficient DNA delivery to O157 strains, we constructed and screened 126 unique chimeras of the Stf protein carrying the N-terminal part of the λ stf protein and receptor binding domains of other phages. Receptor-binding domains of previously described phages infecting *E. coli* as well as from prophages present in our in-house *E. coli* strain collection were identified and cloned as fusions with the λ Stf protein (Supplementary Table 1). The chimeras were cloned under the control of an inducible promoter and transformed into a thermosensitive, λ production strain. In this strain, the λ cos site was deleted to ensure that phage DNA is not packaged (*43*) (s2072, Supplementary Table 2), the λ gpJ was replaced by the 1A2 gpJ variant and the *stf* gene deleted.

To enable the screening of a large number of chimeras against a panel of 22 O157 *E. coli* strains, we established a strategy that relies on the delivery of a reporter DNA payload. A cosmid carrying the λ *cos* packaging signal as well as a GFP reporter and chloramphenicol resistance gene was constructed (p513). We also removed DNA motifs known to be recognised by restriction-modification systems of O157 strains (*44*). (Supplementary Table 3). Initial screening of DNA delivery was performed through transduction assays and spot plating on LB agar with chloramphenicol (Figure 1A). The activity of each λ stf chimera was evaluated by the density of the transductant spot, with denser spots suggesting increased delivery efficiency (Figure 1B and Supplementary Figure 1). This enabled the identification of three candidates (STF-V10, STF74 and STF86) which were further investigated. For these STF chimeras, we measured the fluorescence produced by the DNA payload after delivery to an O157Δ*stx* strain at increasing multiplicity of infection using flow cytometry (Figure 1C). While STF74 and STF86 allowed for a delivery efficiency of about 20% at an MOI around 60, STF-V10 enabled transduction of close to 100% of the target population with an MOI of 20 and was selected for further work.

**Figure 1.**
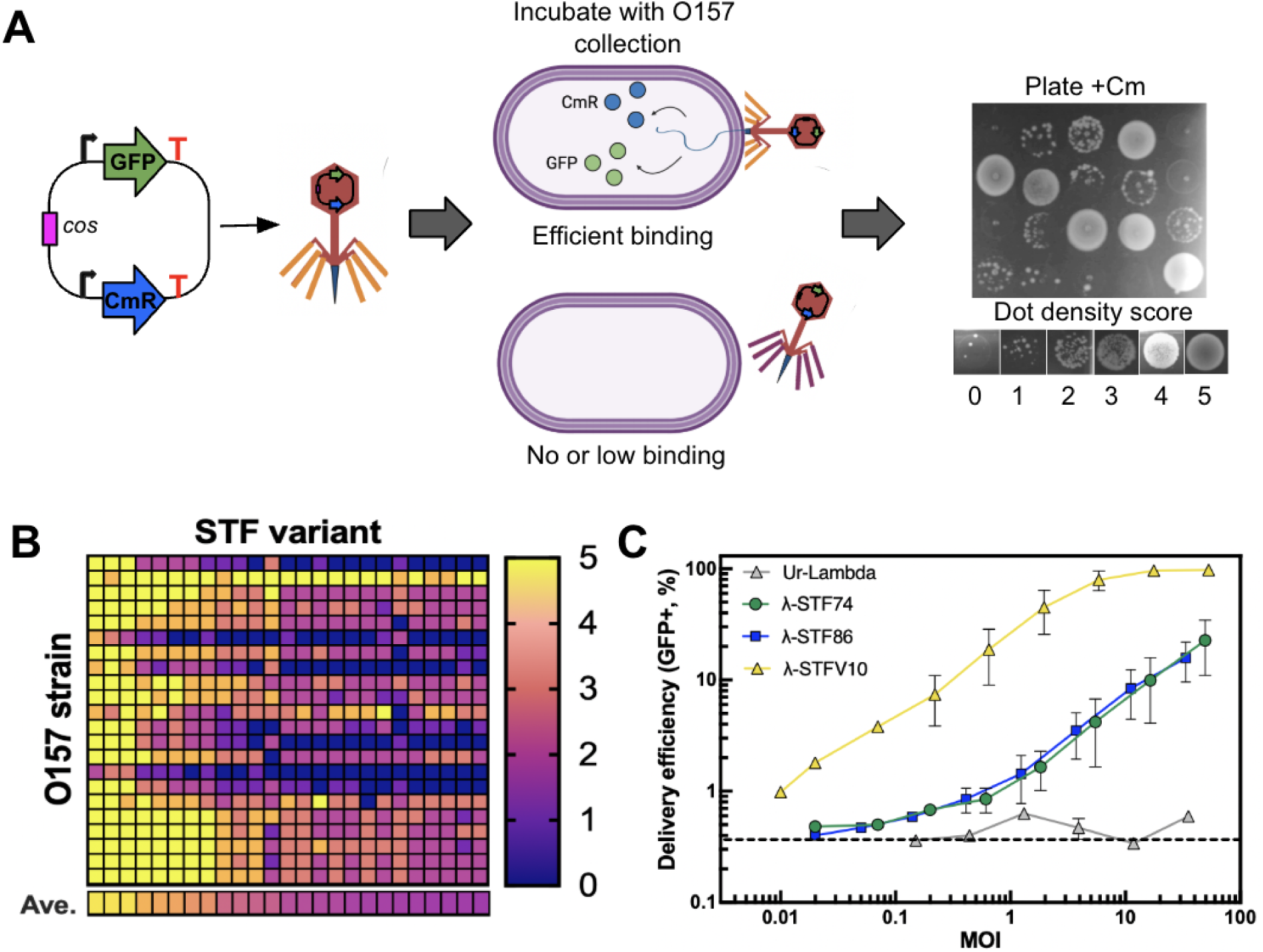
Engineering delivery particles to target a wide range of STEC O157 strains. **A)** Schematic description of the screening assay. Left, architecture of screening plasmid p513 used for plating and flow cytometry assays. GFP, green fluorescent protein; CmR, chloramphenicol acetyltransferase gene; *cos*, λ cos site. Plasmid p513 was packaged into particles carrying the 1A2 gpJ variant and different λ-STF chimeras. Plating assays: dot density was semi-quantitatively scored from 0 (no or <5 colonies per droplet, suggesting poor delivery) to 5 (very dense spot suggesting good delivery). **B)** Semi-quantitative delivery efficiency by agar spot density of cosmids carrying the 1A2 gpJ variant and the best 25 λ-STF chimeras into 22 O157 strains. Data for the complete STF chimera collection can be found in Supplementary Figure 1. Ave., averaged qualitative delivery efficiency for all strains with each STF chimera. **C)** Delivery efficiency as a function of MOI of the top three chimeric STF variants identified in (B) compared to Ur-λ containing the wild-type λ STF. Fluorescence levels were measured by flow cytometry (excitation: 488 nm, emission: 530/30 BP) after transduction of plasmid p513 into O157*Δstx*. Mean and SD of an experiment performed in triplicate. Dotted line shows background fluorescence levels of strain O157*Δstx*.

### Chimeric λ particles can be engineered to ensure resistance to gut proteases

We then assessed the ability of the λ capsid carrying gpJ-1A2 and STF-V10 to survive passage through the mouse gastrointestinal tract. Packaged particles were gavaged to mice and their abundance was quantified in feces samples through transduction assays using as recipient cells either an O157Δ*stx* strain or a K12 strain expressing the EDL933 OmpC ( K12). The number of transducing units measured before gavage were consistent regardless of the strain used. However, titers measured from feces samples on the O157Δ*stx* strain were about 200 fold lower than those measured on strain K12 (Figure 2A). These results suggest that host range determinants required for the transduction of O157 strains are selectively degraded during their transit time in the mouse GI tract. Since we show that the λ STF-V10 chimera is strictly necessary to transduce O157 strains, but not K12 (Figure 2B), we hypothesized that the λ STF-V10 chimera was degraded during passage through the mouse gastrointestinal tract where it is exposed to intestinal proteases.

**Figure 2.**
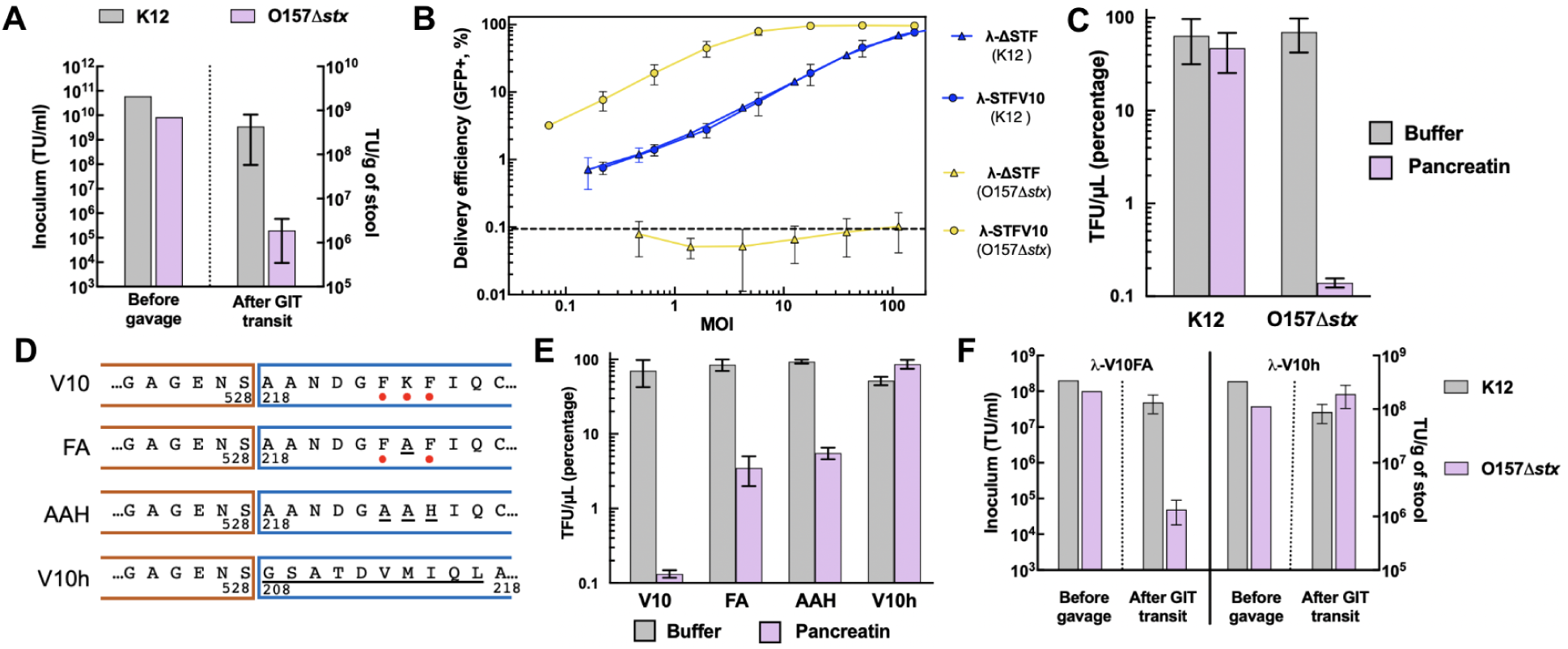
Engineering delivery particles to target STEC O157 in the gut with high efficiency. **A)** *In vivo* stability of engineered particles carrying the 1A2 gpJ and the λ-STFV10 chimera. Cosmid particles from the initial inoculum or recovered from stool samples were titered on two different indicator strains: K12 (to evaluate the functionality of the 1A2 gpJ) and O157*Δstx* (to evaluate the activity of both 1A2 gpJ and λ-STFV10). Mean and SD. **B)** Delivery efficiency of cosmid particles carrying the 1A2 gpJ with and without the λ-STFV10 chimera into K12 and O157*Δstx*. Fluorescence levels were measured by flow cytometry (excitation: 488 nm, emission: 530/30 BP) after transduction of plasmid p513. The dotted line shows the background fluorescence level of the strains. Mean and SD. **C)** Percentage of functional λ-STFV10 chimera cosmids in the absence (grey bars) or presence (purple bars) of pancreatin, measured in a strain for which the STF is dispensable (K12) or required (O157*Δstx*). Mean and SD. **D)** Amino acid sequences of λ STF-V10 tail fiber fusions. Orange boxes, λ-STF moiety; blue boxes, V10 moiety. Numbers represent the amino acid position at which the fusion was done according to the Uniprot accession numbers for λ STF (P03764) and phiV10 tailspike (Q286Z3). Red dots show predicted trypsin and chymotrypsin cleavage sites; underlined letters represent mutations or insertions tested. **E)** Titer of different λ-STFV10 chimeras in the absence (grey bars) or presence (purple bars) of pancreatin, measured in a strain for which the STF is indispensable for injection (O157*Δstx*). Mean and SD. **F)** *In vivo* stability of engineered cosmids carrying the 1A2 gpJ chimera and the pancreatin-resistant (V10h) or a pancreatin-sensitive λ-STF chimera (V10FA). Mean and SD.

To test this hypothesis, we incubated λ capsids carrying gpJ-1A2 and λ STF-V10 in the presence of pancreatin *in vitro*. This resulted in an almost complete DNA delivery inhibition to the O157Δ*stx* strain while delivery to strain K12 was not affected (Figure 2C). An analysis of the λ STF-V10 chimera sequence revealed the presence of an FKF motif at the fusion point that could be targeted by intestinal proteases (Figure 2D). Both F residues are predicted to be cleaved by chymotrypsin using Expasy Peptide Cutter models (*45*), and the K residue is predicted to be cleaved by trypsin. STF-V10 variants in which the FKF motif was replaced by either FAF or AAH partially rescued the sensitivity to pancreatin (Figure 2E). The incomplete rescue suggested that the fusion point may not be folding properly, exposing residues that would otherwise be protected. An analysis of V10 structural homologs (RCSB accession 5W6P) revealed the presence of a short helical peptide upstream of the fusion point we used (position 246-255 of RCSB accession 5W6P). This decapeptide forms a triple helix and connects the C-terminal receptor binding domain to the N-terminal tailspike domain (*46*). Hence, we inserted the putative helical motif-containing decapeptide in our λ STF-V10 design to create λ STF-V10h. STF-V10h was able to efficiently deliver DNA to the O157 strain both before and after pancreatin treatment, showing that it was indeed resistant to proteases.

We finally tested whether STF-V10FA or STF-V10h improved the viability of our capsids during passage through the mouse gastrointestinal tract. Capsids carrying either tail fiber were gavaged to mice and their abundance in feces was measured through transduction assays (Figure 2F). The results corroborated our *in vitro* assays, showing an impaired ability of the λ STF-V10FA capsids to deliver to the O157Δ*stx*, while the λ STF-V10h retained full activity.

### Engineering of a CRISPR-Cas system to target *stx* genes

We then sought to design a CRISPR-Cas12a system able to efficiently kill O157 STEC strains without toxin release by targeting the *stx* genes. To ensure that our guide RNAs would target all possible variants of the *stx* genes, we compiled a list of 1292 genomes corresponding to STEC O157 strains. 13 different *stx* subtypes were found, *stx2c* being the most frequent (Supplementary Table 4). To implement guide RNA redundancy, a final set of 4 guide RNAs targeting the gene of the *stxA* subunit was chosen: two guides targeting the *stx1* locus and two guides targeting the *stx2* locus (Supplementary Table 5). The *stx1* guide 1 targets all *stx1* gene variants while *stx1* guide 2 covers over 99% of the *stx1* gene variants. Both *stx2* guides target all *stx2* variants. Our guide design therefore provides an excellent coverage of the *stx* sequence diversity while providing redundancy.

A CRISPR array containing all 4 guide RNAs was cloned in a cosmid carrying *cas12a* under the control of a pSrpR promoter (*47*). We further designed our cosmid to carry a conditional origin of replication (*32*), ensuring that it can replicate in a production strain for *in vivo* packaging, but not in recipient bacteria. This design ensures that our DNA payload cannot be maintained in bacterial strains that receive the DNA payload without being killed by it. Both the *cas12a* gene and the conditional origin of replication were recoded to remove O157 restriction sites.

We also replaced the antibiotic resistance marker by a recoded *thyA* gene containing no O157 restriction motifs while this gene was deleted in our production strain. This ensured the maintenance of our plasmid in the production cells while preventing the dissemination of an antibiotic resistance gene. The engineered λ capsid carrying STF-V10h and gpJ-1A2 was then used to package this DNA payload (p1392) yielding EB003 (Figure 3A).

**Figure 3.**
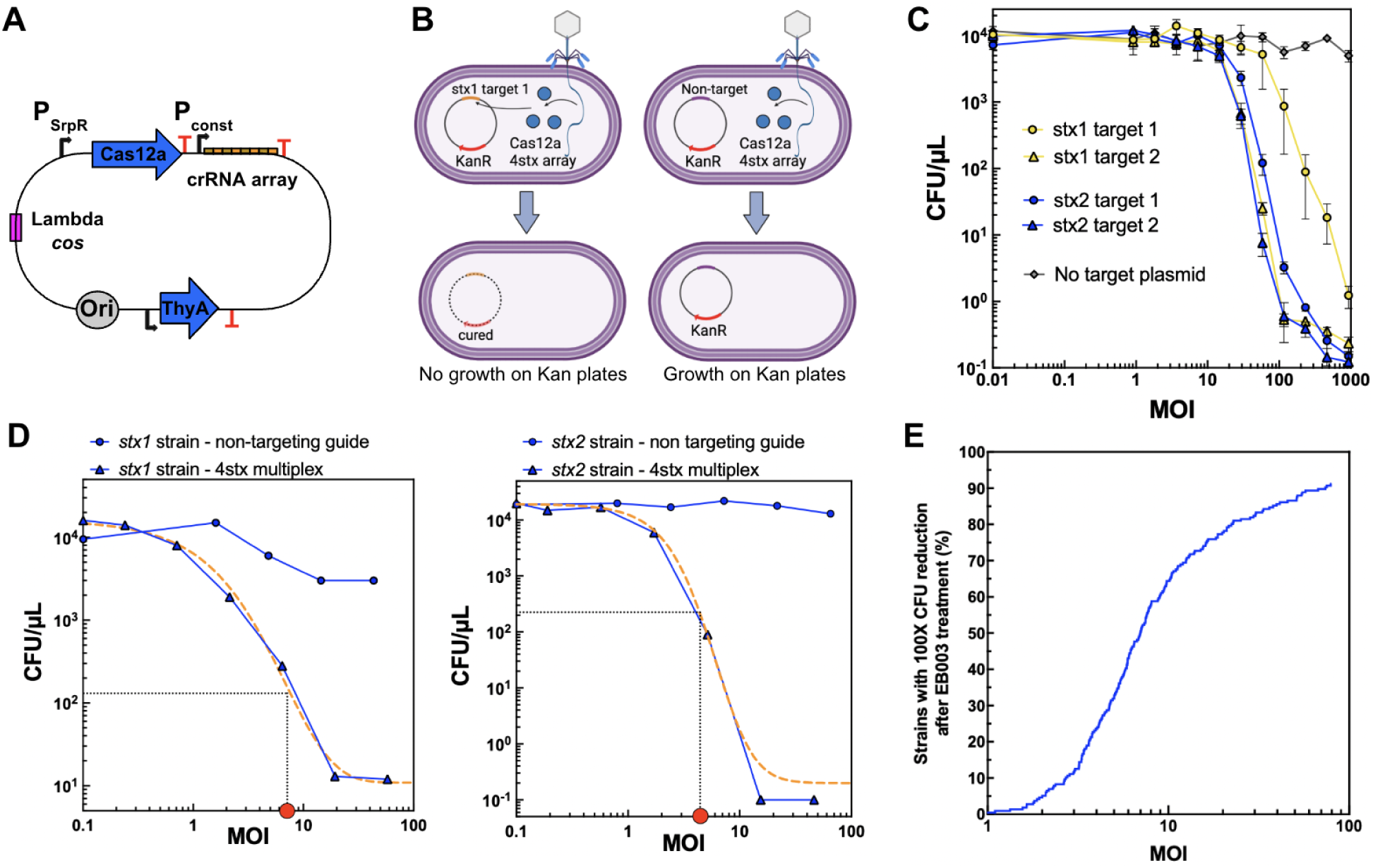
Engineering a DNA payload encoding a Cas12a system that targets all *stx* gene variants. **A)** Schematic representation of the packaged payload p1392. *Ori*, conditional origin of replication; P_srpR_, repressible *cas12a* promoter; P_const_, 4stx crRNA array constitutive promoter; *thyA*, thymidine auxotrophy marker. T signs, transcriptional terminators. Mean and standard deviation of an experiment performed in triplicate. **B)** Schematic representation of the plasmid curing assay to assess the activity of each guide in the multiplexed array. **C)** Plasmid curing activity measured as CFU counts on LB agar supplemented with kanamycin at different MOIs in O157Δ*stx* carrying a plasmid with each Cas12a target or no target control. Mean and SD. **D)** Killing of O157 strains carrying either the *stx1* gene only (left panel) or the *stx2* gene only (right panel) at different MOIs with payload p1392. Orange dashed line, a generalized logistic function was fitted to the data (Methods). The curve fit is used to calculate the MOI for which a 100X reduction in CFUs is observed (red dot), as shown in panel E. **E)** Percentage of 216 O157 STEC strains showing a 100X reduction (99%) in CFUs after treatment with EB003 as a function of MOI *in vitro*.

To assess the activity of the 4 *stx* guide RNAs, we cloned the targets of each guide on a plasmid and performed plasmid curing assays in a O157Δ*stx* strain. Upon treatment with EB003, Cas12a is expected to cleave the plasmid, restoring sensitivity to kanamycin (Figure 3B). Three of the guide RNAs encoded in EB003 allowed for a 10^4^ fold CFU reduction of kanamycin plasmid-containing strains at MOIs below 100, while one of them allowed for about 10 fold reduction (Figure 3C). One of the challenges of CRISPR system design is that not all guide RNAs may cleave their target with the same efficiency (*48, 49*). However, our design contains two guides per *stx* target to allow for redundancy, limiting the impact of a guide RNA with sub-par efficiency.

### EB003 efficiently kills a wide range of STEC O157:H7 clinical isolates *in vitro*

We next assessed the ability of EB003 to eliminate wild type STEC O157 strains carrying either *stx1* or *stx2* genes *in vitro*. Cultures of both *stx1* (s1592) and *stx2* (s1594)-carrying O157 strains were treated with increasing concentrations of either EB003 or a control cosmid encoding a non-targeting guide RNA. While treatment with the control cosmid did not impact the number of colony forming units, treatment with EB003 reduced them by a factor of 1.3×10^3^ for strain s1592 and 2×10^5^ for strain s1594 (Figure 3D), representing more than 99.9% of the cells eliminated at MOIs below 100.

To assess the potential impact of EB003 in the clinic, its antimicrobial activity was tested *in vitro* on a large collection of recent clinical isolates of STEC O157 strains. These strains were isolated in the UK (92 strains) and France (124 strains) between 2015 and 2018, and carry the most prevalent *stx* subtypes with *stx2a, stx2c* and *stx1a* present in 58%, 57% and 33% of the isolates, respectively (Supplementary Table 6).

We assessed the cumulative fraction of the collection that could be eliminated with 99% efficiency (100-fold reduction of colony counts) after treatment with increasing concentrations of EB003. We found that 91% of the STEC O157 strains could be eliminated with MOIs below 100 (Figure 3E and Supplementary Figure 9).

### EB003 kills STEC O157 without toxin release

Nine STEC O157 isolates carrying *stx* subtypes *stx2a, stx2c, stx1a + stx2c,* and *stx1a* + *stx2a* were selected based on the clinical relevance of the *stx* subtype they carry (Supplementary Tables 2 and 6). We evaluated the antimicrobial activity and toxin release after *in vitro* treatment with EB003 or the antibiotic ciprofloxacin (Figure 4A).

**Figure 4.**
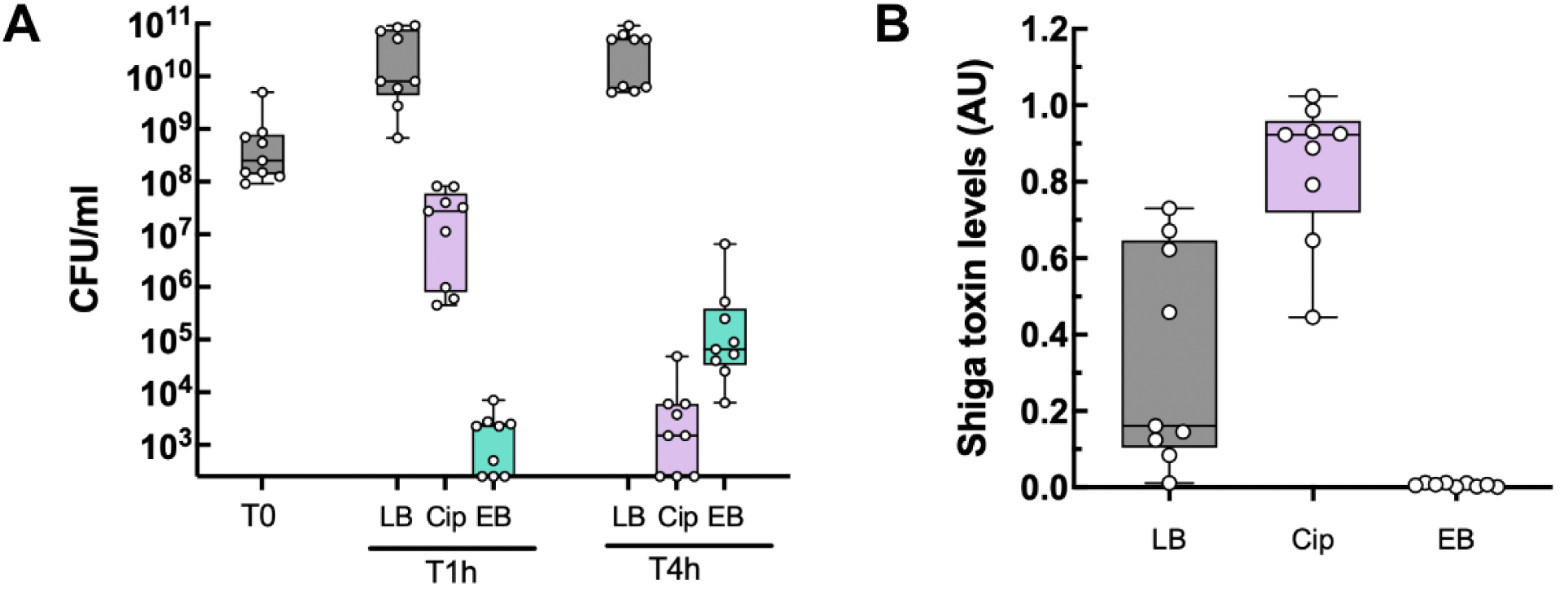
Engineered O157-targeting delivery particles with a DNA payload encoding a type Cas12a system eliminate clinical isolates of STEC O157 without toxin release *in vitro*. A) Time course assay of nine STEC O157 strains treated with EB003 (EB), ciprofloxacin (Cip) or media (LB). Surviving bacteria were measured as CFU mL^-1^ on agar plates after one or four hours after the different treatments. Each circle represents a different strain (Materials and Methods). **B)** Levels of Stx toxin in culture supernatants of nine different STEC O157 strains after four hours of treatment with lethal doses of ciprofloxacin (Cip) or EB003 (EB) as measured by ELISA. LB shows toxin levels of strains treated with media only. Each circle represents a different strain. Box: 25th–75th percentiles; whiskers: min–max; horizontal line: median.

The maximum CFU reduction was observed after 1 hour of incubation of O157 cultures with EB003 and after 4 hours with ciprofloxacin. Shiga toxin release measured by ELISA after 4 hours of treatment showed high toxin levels after ciprofloxacin treatment as previously described (*50*). In contrast, EB003 treatment did not produce any detectable toxin release (Figure 4B).

### EB003 effectively targets O157 STEC in the mouse gut

To assess the efficacy of EB003 *in vivo,* a mouse model of intestinal colonization with STEC O157 was developed. To promote STEC O157 colonization we isolated a streptomycin-resistant mutant of O157 strain s1594 (s17769), which was inoculated orally to streptomycin-treated BALB/c mice. This led to a stable intestinal colonization for a period of 3 days at levels of around 10^8^ CFU g^-1^ of stool (Supplementary Figure 4). Five days following O157 strain inoculation, mice were orally gavaged with EB003. *E. coli* colonization was monitored by plating dilutions of stool samples collected from each mouse on Drigalski agar (Figure 5A). After administration of a single dose of 10^12^ EB003 particles (n=20 mice), O157 levels decreased by a median of 3×10^3^ CFUs 8 hours after treatment, while no impact was observed on the colonization levels of an isogenic strain in which the Shiga toxin gene was deleted (s2185, Figure 5B). After 24 hours, the level of O157 bounced back to a median level of 6×10^6^ CFU g^-1^ of stool, indicating that surviving clones were able to repopulate their niche, although they represented on average less than 0.01% of the original population.

**Figure 5.**
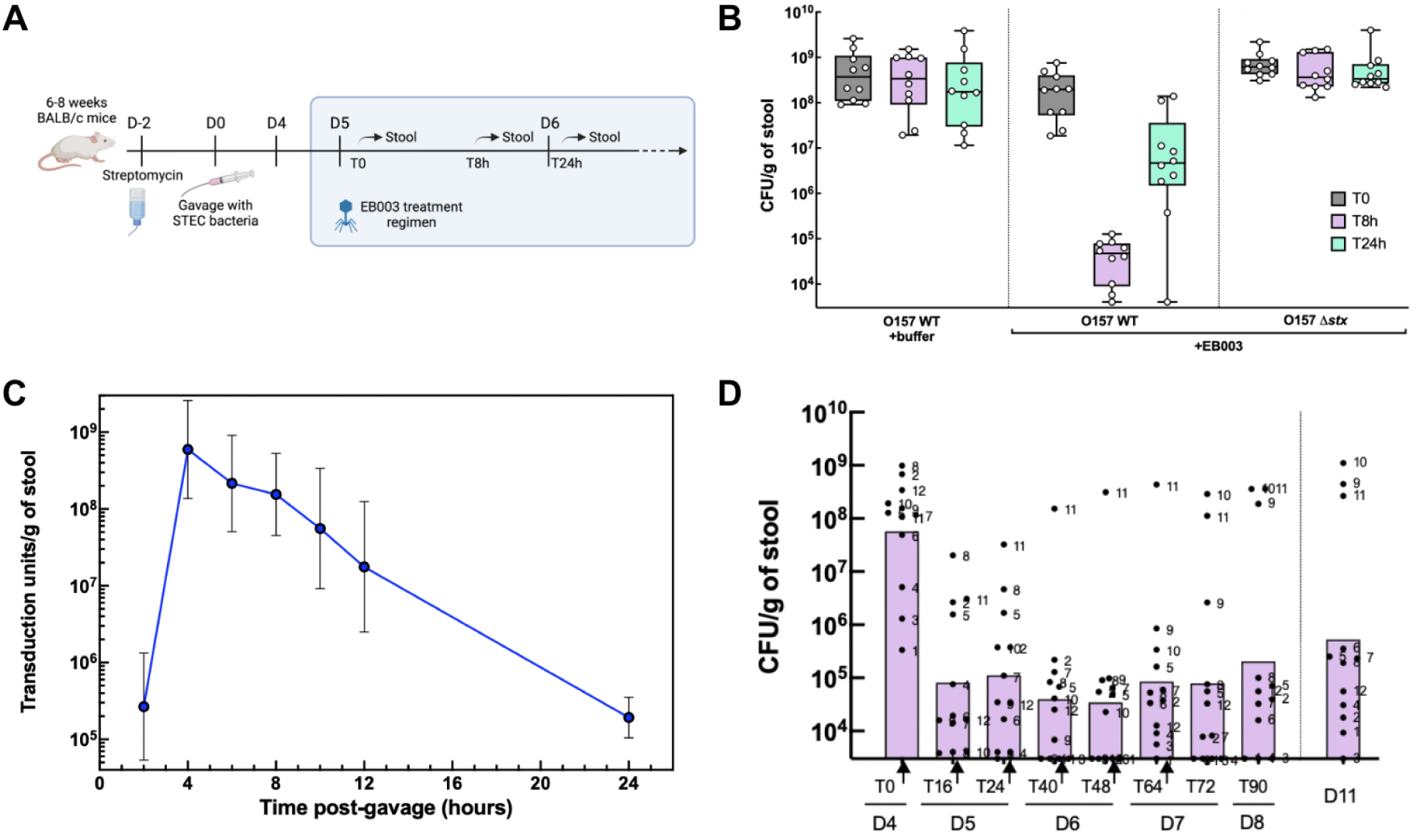
Oral administration of EB003 can eliminate STEC O157 in a mouse model of intestinal infection. **A)** Mice were conditioned with streptomycin to clear up a niche for the engraftment of orally-gavaged STEC O157 bacteria. Treatment with EB003 was started at day 5, following the regimens described in the text. **B)** Effect of a single 10^12^ TFU dose of EB003 on the colonization levels of a STEC O157 strain (O157 WT) or its isogenic mutant strain deleted for the *stx* gene (O157Δ*stx*). Box: 25th–75th percentiles; whiskers: min–max; horizontal line: median. **C)** Kinetics of fecal shedding of EB003 after a single oral dose in mice. EB003 particles were measured by transduction on fresh fecal samples for individual mice. Mean and SD. **D)** Effect of a bi-daily treatment regimen for 4 days on STEC O157 colonization levels. Animals were dosed at T0, T16, T24, T40, T48 and T64 hours (black arrows). Individual mice are indicated by their unique number throughout the experiment. Bar heights represent the median.

We then assessed whether treatment with repeated doses of EB003 would prevent the rebound of the O157 strain colonization described above and maintain the reduction of O157 carriage for at least a week. This corresponds to the treatment window for STEC-infected patients as HUS onsets after a median period of 7 days post infection (*51*). We first determined the pharmacokinetics of λ particles harboring 1A2 gpJ and λ-STFV10h after passage through the gastrointestinal tract of non-colonized mice. As can be seen in Figure 5C, the shedding peak of the particles occurs 4 hours after gavage and decreases steadily to reach about 5% of the maximum peak after 12 hours. This time point was chosen in order to maintain a high level of particles in the mouse gut in a multi-dose experiment. Hence, O157-colonized mice were administered with two daily 10^12^ TFU doses every 12 hours of EB003 over 3 days. This dose regimen allowed to reach and maintain a ∼99.9% reduction of STEC O157 colonization levels for the whole duration of the treatment for all animals except one that remained refractory to the treatment (Figure 5D). Three days after the last dose, the O157 load remained lower than 10^6^ CFU g^-1^ of feces for 8/11 mice.

### EB003 allows for symptom mitigation in an infant rabbit STEC model

In our experimental setup the direct inoculation of STEC bacteria to mice did not recapitulate symptoms, consistently with previous reports (*52, 53*). An infant rabbit model of STEC O157-mediated diarrheal disease was used to assess the efficacy of EB003 in mitigating disease symptoms, since infant rabbits infected with STEC exhibit intestinal and renal symptoms comparable to humans (*54*). Two-day old New Zealand White rabbits were orally infected with 10^7^ CFU of O157 s17773 (Day 0). Three days after infection, when all animals presented diarrhea, treatment was initiated: two groups of rabbits received orally 250 μL of either sucrose/bicarbonate buffer (13 animals) or EB003 active substance (3×10^12^ particles, 12 animals) twice a day every 8 hours for 4 days (Figure 6A). After the complete treatment regimen, the severity of diarrhea was significantly reduced in animals that received EB003 (mean score of 0.8) compared to control animals that received only buffer (mean score of 1.9, p<0.001, Figure 6B). The median bacterial burden recovered from EB003-treated animals was also significantly lower than in buffer-treated animals (Figure 6C), with a 20 fold reduction (95%) in the small intestine and colon and a 2500 fold reduction (99.96%) in the caecum.

**Figure 6.**
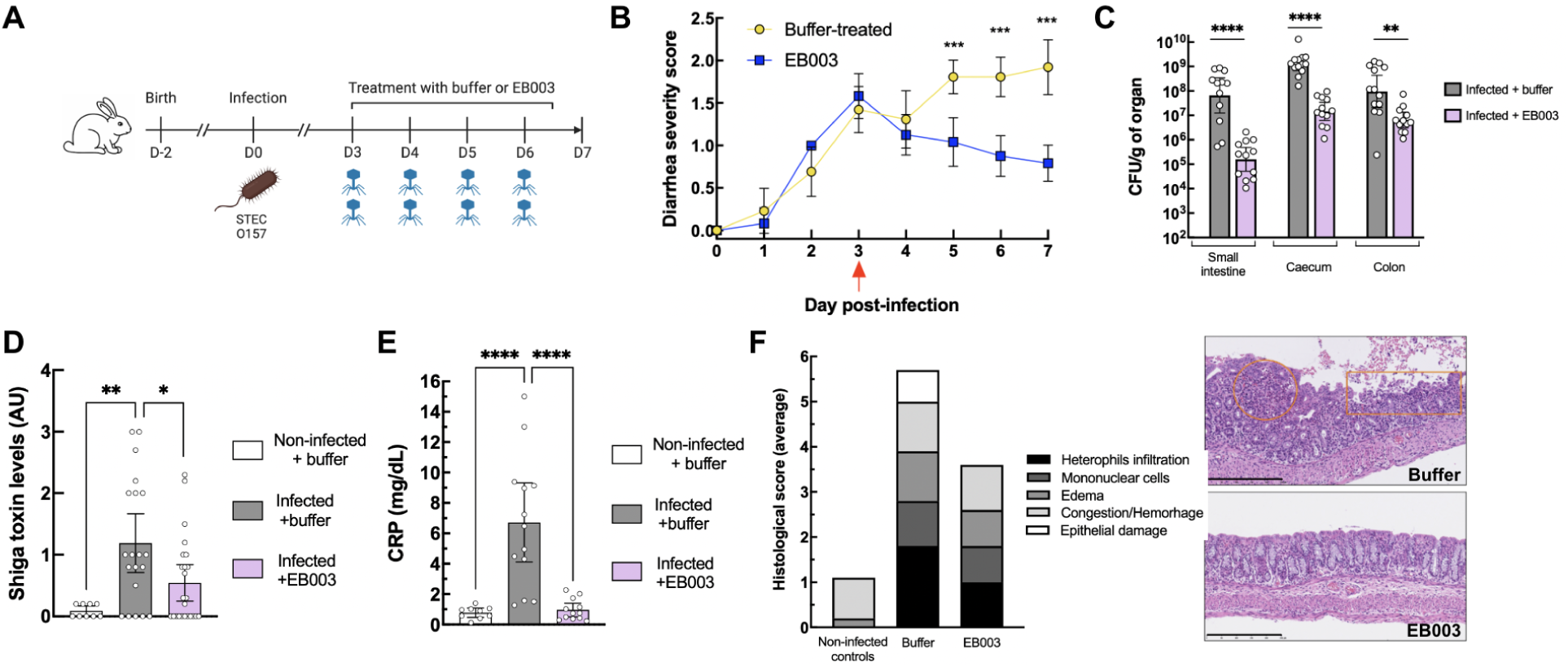
Oral treatment with EB003 rescues symptoms of STEC O157 infection in an infant rabbit model. **A)** Two-day old infant rabbits were given orally STEC O157, and treated 3 days later twice daily with either buffer or EB003. **B)** Evolution of diarrhea severity over the course of the experiment. Plotted: mean with 95% CI. ***:p<0.001 by unpaired t-test for each time point. **C)** STEC CFU in the small intestine, caecum and colon of infected rabbits at Day 7. Plotted: geometric mean with 95% CI. Mann-Whitney test. ****: p<0.0001; **: p<0.01. Each dot represents a different animal. **D)** Stx toxin levels in the colon, as measured by ELISA on total organ homogenates. Plotted: mean with 95% CI. Mann-Whitney test. **: p<0.01; *: p<0.05. Each dot represents a different animal. **E)** Levels of C-reactive Protein (CRP) in the blood of animals at Day 7. Plotted: mean with 95% CI. Mann-Whitney test. ****: p<0.0001. Each dot represents a different animal. **F)** Histopathological assessment of colon biopsies at Day 7 (left graph). The scoring was performed by a third-party blinded to the group attribution of each animal, and the average of all the animals within a group is shown. Representative pictures are also provided (right). Buffer: infected, buffer-treated animals with superficial acute erosive colitis with mild heterophilic inflammation (circle) and focal erosion (rectangle); EB003: infected, EB003-treated animals, normal colon. Black bars represent 250 µm.

Measurements of Stx toxin levels were done using a Shiga Toxin Quick Chek assay, which allows for the semi-quantitative determination of toxin presence. Animals treated with buffer showed a significantly higher Stx signal than EB003-treated animals (Supplementary Figure 7). To allow for a better quantification of Stx, another experiment was performed to measure Stx by ELISA from colon samples of infected mice. This showed a significant reduction after treatment with EB003 compared to buffer (Figure 6D).

The levels of C-reactive protein were also quantified, showing that treatment with EB003 allowed for a marked reduction in inflammation. Buffer-treated, infected animals showed CRP levels of 6.72 g dL^-1^ on average, while EB003-treated animals presented CRP levels indistinguishable from those of non-infected animals (0.96 vs 0.78 g dL^-1^, respectively, Figure 6E). Finally, a histological analysis of colon and ileum samples was performed to compare the non-infected controls to the O157*stx*-infected buffer- or EB003 treated animals (Figure 6F and Supplementary Figure 5). Non-infected controls showed normal colon and ileum morphologies while infected rabbits treated with buffer presented damaged colon and ileum samples. Animals treated with EB003 presented minimal damage to the colon and ileum tissues. Furthermore, animals receiving only buffer exhibited a 163% weight gain by the end of the study, whereas EB003-treated animals exhibited a 182% weight gain, comparable to that of non-infected controls (Supplementary Figure 6).

In conclusion, post-symptom administration of EB003 effectively reduced diarrhea severity and intestinal burden of STEC O157, decreased stx production, accelerated tissue repair and restored normal weight gain in an infant rabbit model.

## Discussion

Here we demonstrate the potential of EB003, a non-replicative CRISPR-Cas antimicrobial, for the treatment of STEC infections. We show efficient targeting of the vast majority of clinically-relevant O157 *E. coli* strains *in vitro*, efficient reduction of the bacterial load in a mouse gut colonization model and symptom mitigation in an infant rabbit disease model, including reduced toxin production.

The development of EB003 involved the engineering of a delivery vector based on phage λ and a DNA payload carrying a CRISPR-Cas12a system targeting the Shiga toxin genes. The lambda tail was engineered to carry a modified gpJ protein, ensuring the recognition of a primary receptor, OmpC, that is consistently expressed by *E. coli* in the gut environment. In addition, the side tail fibers (STF) were modified to recognize the O157 antigen, ensuring efficient delivery into the vast majority of clinical isolates of O157 *E. coli* strains. The specificity of phage-based therapeutics is a key factor in their clinical translation, as they must effectively target epidemiologically relevant strains.

While O157 *E. coli* remains a leading cause of STEC infections (>70% of outbreaks in the USA between 2010 and 2017 (*55*)), other serogroups, such as O26 and O103, are becoming increasingly prevalent (*56*). Previously, we demonstrated that the gpJ chimera used in this study efficiently recognizes most OmpC variants from *E. coli*.

However, our engineered STF-V10h is unlikely to effectively target O-antigens beyond O157. Expanding or screening our existing λ-STF and gpJ chimeric libraries could identify optimized delivery vectors for these and other emerging STEC serotypes.

The DNA payload of EB003 was designed to express Cas12a in target bacteria without the need for plasmid replication. The CRISPR array specifically targets four conserved sites in *stx1* and *stx2*, which corresponds to >99% of all *stx* variants. Importantly, we confirmed that EB003 efficiently kills O157 strains without triggering toxin release. Its killing efficiency was validated against a collection of over 200 clinical isolates, demonstrating the robustness of EB003 against the natural genetic diversity of O157 STEC.

Unlike natural phages, EB003 does not replicate within target bacteria, requiring administration at a high MOI to achieve a significant reduction in O157 bacterial load. While an MOI of 10 to 100 was effective for most strains *in vitro*, it was unclear whether this would be feasible in an animal model, where bacterial accessibility and drug stability pose challenges. Indeed, our initial EB003 designs were susceptible to proteolytic inactivation, necessitating further engineering of the phage STF. The optimized particles successfully delivered the CRISPR-Cas12a system to the majority of target O157 STEC in both mouse and rabbit models following administration of 10¹² particles.

In the mouse model, we observed a drastic reduction in bacterial load within 8 hours of treatment, though partial recolonization occurred at 24 hours. This rebound is likely influenced by the continuous administration of streptomycin, which strongly favors the recolonization of the streptomycin-resistant O157 strain used. Notably, this dynamic does not reflect clinical infections, as STEC O157 does not establish stable colonization in patients and is naturally shed after a median of 13 days (*57*). In this context, accelerating bacterial clearance while preventing toxin release is expected to provide clinical benefit.

Crucially, repeated administration of EB003 over three days maintained bacterial suppression despite streptomycin-driven selection pressure. Treatment failure in one animal may have resulted from the emergence of a resistant mutant, highlighting the importance of investigating resistance mechanisms to enhance treatment efficacy and ensure that any surviving bacteria do not produce toxins. Importantly, in the infant rabbit disease model, EB003 mitigated symptoms in all treated animals, suggesting that resistance emergence does not pose an immediate threat to therapeutic efficacy.

Since their initial proof-of-concept demonstration in 2014 (*26, 27*), few studies have explored the feasibility of CRISPR-Cas antimicrobials for treating specific infections in realistic preclinical models. In this work, we provide evidence that a CRISPR-Cas system can effectively mitigate infection symptoms by selectively targeting a bacterial population within a complex microbiome.

While others have investigated lytic phages carrying CRISPR-Cas systems (*58*), these engineered phages can eliminate target bacteria independently of the CRISPR-Cas action, making their therapeutic properties largely similar to classical phage therapy. In contrast, EB003 relies entirely on CRISPR-Cas for bacterial killing. EB003 does not produce progeny and its DNA payload is engineered to be non-replicative to avoid the maintenance of transgenes in bacteria that it does not kill. This additional step should facilitate regulatory approval in comparison to replicative, genetically modified phages. While challenges such as dosing, resistance emergence, and delivery optimization remain, our findings provide a compelling foundation for the further development of CRISPR-based therapeutics. Future work will focus on broadening target coverage and assessing efficacy in clinical settings. Ultimately, EB003 represents a promising step toward the next generation of precision antimicrobials for combating STEC infections and potentially other bacterial pathogens.

## Materials and Methods

### Study design

The goal of this study was to evaluate the *in vitro* and *in vivo* efficacy of our non-replicative, CRISPR-Cas-based antimicrobial drug candidate EB003 on *E. coli* O157 strains and to develop translational methods for use in the clinic. EB003 antimicrobial activity in a diverse O157 strain collection and toxin abrogation with different *stx* gene variants was first validated *in vitro*. Subsequently, we engineered EB003 to be able to withstand passage through the mouse gut and performed pharmacodynamics studies to obtain a dose regimen for treatment. We then confirmed the antimicrobial activity of our drug candidate in a mouse model. To verify symptom mitigation of our drug candidate, we used infant rabbit models. In both mouse and rabbit experiments, sample sizes were established by logistical and ethical reasons. We selected groups of at least 10 animals which should enable to measure a difference in bacterial CFUs between treated and non-treated animals (*59*). Mouse models were performed by Eligo Bioscience. Rabbit models were performed by Vivexia on behalf of Eligo Bioscience. Animals were divided into two groups, one receiving the EB003 drug candidate and the other one sucrose-bicarbonate buffer. One of the rabbits in the EB003 group had to be euthanized at day 4 post-infection since it displayed a major weight loss and a strong dehydration. For the data shown in Figure 4D, one of the animals belonging to the EB003 group had to be euthanized at day 2 post-infection since it displayed a major weight loss and a strong dehydration. Histopathology data of infant rabbit models were analyzed in a blinded fashion.

### Strains and media

For experiments in Figures 1 and 2, O157Δ*stx* strains s17465 and s2185 were used. Strain K12 is strain s14269. The O157 strains s17769 and s17773 were used in mouse and rabbit *in vivo* experiments, respectively. For mouse models with an isogenic O157Δ*stx* variant, strain s2185 was used. Strain genotypes are listed in Supplementary Table 2. The O157-strain collections were isolated in the UK (92 strains, Public Health England collection) and France (124 strains, Institut Pasteur collection) between 2015 and 2018 (See Supplementary Methods).

### Cloning and plasmid construction

To generate λ STF chimeras with receptor-binding proteins from previously described *E. coli*-targeting phages, their crystal structure was analyzed from PDB entries, if available, to identify a fusion point. If no crystal structure was available, a BLAST search in the PDB database was performed to find the closest structural homolog. Since these RBPs showed no homology to λ STF, they were all cloned at position 528 of the λ STF (Uniprot P03764). To generate λ STF chimeras with receptor binding proteins from *E. coli* prophages, genomic sequences of an in-house *E. coli* collection were analyzed with PHASTER (*60*) and their putative RBPs identified. Then, alignments with the λ STF protein were performed to identify RBPs with and without homology to the λ STF. For RBPs with homology, the first homology stretch to the λ STF was chosen as an insertion point. For RBPs without homology to λ STF, a structural analysis with the PDB database or an HHMER analysis were performed to identify an insertion point. Sequences for the λ STF-V10 chimera were obtained by synthesizing and fusing the tail fiber protein from *Escherichia coli* phage phiV10 (amino acid position 218 onwards, Uniprot Q286Z3) to phage λ STF (amino acid positions 1-393, Uniprot accession P03764). For the λ STF-V10h variant, the sequence of Uniprot Q286Z3 from amino acid 208 onwards was fused to λ STF. Plasmids are listed in Supplementary Table 7. Production of packaged cosmids and delivery efficiency assay were done as in (*32*) (Supplementary methods).

### *In vitro* killing and plasmid curing assay

For killing assays, strains O157 s1594 and O157 s1592 were used. For plasmid curing assays, strain O157Δ*stx* s2185 carrying plasmids p766, p767, p768, p840 or p764 maintained with 100 µg mL^-1^ kanamycin was used. For each O157 strain, an overnight culture was diluted 1:100 and grown in liquid LB + 5 mM CaCl_2_ at 37°C until an OD600 of 0.5-0.7 (with or without kanamycin, as needed). Then, the samples were diluted to an OD of 0.025 in fresh LB + 5 mM CaCl_2_. 90 µL of the cell suspensions were treated with serial dilutions of 10 µL packaged phagemids carrying the p1392 cosmid for 30min at 37°C, then plated on LB agar (killing assays) or LB agar supplemented with 100 µg mL^-1^ kanamycin (plasmid curing) and incubated for 18 hours at 37°C. CFUs were counted to assess the antimicrobial efficacy of EB003 (killing assays) or plasmid curing activity as a function of MOI. For curve fits and killing data shown in Figure 3D, Figure 3E and Supplementary Figures 3 and 9, the raw colony counts and MOI values were used to fit a generalized logistic function [latex code <math<Y(t) = A + { K-A \over (C + Q e^{-B t}) ^ {1 / \nu} }</math>] using the minimize function from the lmfit Python package(*61*) with the Nelder method. For Figure 4A and Supplementary Figure 9, EB003 was manufactured and purified by Jafral (Slovenia). For supplementary Figure 3, an in-house small scale production was performed.

### *In vitro* toxin release and killing assay

Overnight cultures of strains s1594, s13699, s13729, s13791, s13715, s13765, s13734, s13732 and s13760 (Supplementary Table 2) were diluted 1:100 in fresh LB media supplemented with 5 mM CaCl_2_ at 37°C for 2 hours with 180 rpm shaking. OD_600_ was then adjusted to 0.6. Each strain was treated with 100 ng mL^-1^ ciprofloxacin, EB003 (2×10^12^ particles) or LB and incubated at 37°C with 180 rpm shaking. At time 0, time 1 hour and time 4 hours, samples were serially diluted and spotted on LB agar plates to perform CFU count after 18 hour incubation at 37°C. To analyze toxin release with ELISA (adapted from (*62*)), each incubation was centrifuged and the supernatant filtered through a 0.45 µm pore size membrane and stored at -20°C until measurement. ELISA plates were coated with globotriaosylceramide from porcine erythrocytes (Nacalai). After blocking, 100 µL of the O157 *stx* strain supernatant was added as serial dilutions in triplicate to the ELISA plate. The plate was incubated for 3 hours at room temperature. Primary mouse antibody against *stx* toxin 2 (sc-52727, Santa Cruz Biotechnology) was added and incubated at 37°C for one hour. After washing, the secondary antibody goat anti-mouse HRP (62-6520 Thermo Fisher) was added for one hour at 37°C. After washing, 100 µL of One Step TMB substrate (Sigma Aldrich) was added to the wells and incubated for 15 minutes in the dark. The absorbance was then read with a TECAN reader at 450 nm.

### Guide RNA design

An in-silico analysis identified short sequences of 28 base pairs, including the protospacer adjacent motif (PAM) 5’-TTTV-3’ conserved in all STEC O157 strains analyzed. Sequencing data originate from EnteroBase (https://enterobase.warwick.ac.uk/species/index/ecoli); NCBI RefSeq; NCBI SRA BioProjects PRJNA248042 and PRJNA259645; and an in-house collection including isolates from Public Health England (PHE) and the Pasteur Institute. A total of 1292 genomes were included in the analysis (Supplementary Table 4). For the raw read sequencing data of BioProjects PRJNA248042 and PRJNA259645, a local assembly of the *stx* genes was performed by extracting the reads matching a comprehensive database of Stx protein sequences; we used DIAMOND (*63*) to align the translated nucleotide sequences of the short reads to the database and ABYSS 2.0 (*64*) to assemble the filtered read sets. For the in-house collection, we assembled the full genomes using SPADES (*65*). For the other datasets, we used the assemblies already available on the respective databases. *stx* genes were identified and subtyped by aligning the assemblies to the Phylotyper *stx* databases. The best BLAST hit was used to assign the subtype. The subunit A of each *stx* system was extracted and used for downstream analysis. Redundant sequences were removed from the resulting dataset, and each representative sequence was assigned a weight equal to the number of isolates in which it appears. Supplementary Figure 8 shows an extract from the multiple sequence alignment of the representative sequences for *stx2*, each associated with its weight. The non-redundant *stx* subunit A sequences were analyzed with FlashFry (*66*) to discover potential targets for the guide RNAs. The resulting targets were scored based on the number of isolates in which they appear. From the top scoring targets, we selected 2 guides for *stx1* variants and 2 guides for *stx2* variants. Percentages of targeted isolates and gene variants are shown in Supplementary Table 8.

### Mouse model

Specific pathogen-free 5 to 9 week old female BALB/cYJ mice were supplied by Charles River Laboratories and housed in an animal facility in accordance with Institut Pasteur’s guidelines and European recommendations. Animal procedures were approved by the Institut Pasteur (approval ID: 180015) and the French Research Ministry (APAFIS ID: 25949) and animal experiments were performed in compliance with applicable ethical regulations. Water and food were provided *ad libitum*. Animals were acclimated for 5 days before streptomycin sulfate (5 mg ml^-1^, Sigma-Aldrich) was added to the autoclaved drinking water to decrease the number of facultative aerobic/anaerobic resident bacteria (*67*). Drinking water containing streptomycin was prepared fresh weekly. Three days later (D0), mice were orally gavaged with 1×10^7^ cfu of strain s17769 supplemented with 100 µg mL^-1^ streptomycin and resuspended in 200 µl of sterile gavage buffer (20% sucrose, 2.6% sodium bicarbonate, pH 8). Starting at D5, mice were orally administered 200 µl of either gavage buffer or packaged cosmids diluted 1:1 in buffer. For mice experiments, EB003 was manufactured and purified by Jafral (Slovenia). The appropriate dose was calculated by titration. *E. coli* CFUs were calculated by resuspending fecal material in PBS and plating on Drigalski agar.

### Pharmacokinetics of packaged particles

1.5×10^10^ total λ chimeric particles harboring 1A2 gpJ and λ-STFV10h were orally gavaged as explained above to seven female BALB/cYJ mice, which were not colonized with an O157 strain. Mouse stool was collected every 2 hours for the first 12 hours, and a final collection at 24 hours after gavage. Engineered particles present in feces were enumerated as described in the mouse model above.

### Infant rabbit model

Infant rabbit experiments were performed by Vivexia (Dijon, France) on behalf of Eligo Bioscience. Each litter was nursed by the mother once daily throughout the study. All procedures using animals were approved by the Animal Care and Use Committee C2EA (local ethics committee) agreed by French authorities (APAFIS#31111-2021042108428770 v2). Four litters of two-day old rabbits from the BSL2 area (University of Burgundy, Dijon, France, Agreement N° C 21 464 04 EA) were given orally 200µL of strain s117773 or 200µL of bicarbonate/sucrose solution (non-infected control group). At D3 post-infection, infected and non-infected animals were split: one group received 300µL sucrose/bicarbonate buffer; the second group received 300µL of EB003 (1.5×10^12^ particles). Treatments were administered orally twice a day on D3, D4, D5 and D6 post-infection. At D7 post-infection, all animals were sacrificed. The diarrhea was scored as in (*68*). For determination of CFU counts, parts of the small intestine, caecum colon and organs were excised, homogenized in 5 or 10 times their weight in PBS and plated on Drigalski agar plates supplemented with 100 µg mL^-1^ streptomycin (Sigma Aldrich) and CFUs counted after overnight incubation. Shiga toxin detection was performed at D7 with a Shiga Toxin Quik test (Abbott, France). CRP levels were measured in blood serum (Cerbavet, Massy, France) with a (PTX1) Rabbit ELISA Kit (Abcam, ab157726). For histopathological analysis at D7 post-infection, a piece of distal colon or ileum was excised from all animals (10mm slice) and then kept in histological cassette in 10% NBF. All samples were transferred into 70% ethanol 48h later and then sent for paraffin-embedding and further histological analysis (H&E staining) to Cerba Research (Montpellier, France). For the detection of *stx2* by ELISA, a ProSpecT Shiga Toxin *E. coli* (STEC) Microplate Assay (Thermo Fisher) was used.

Colon samples from rabbits treated with EB003-2, composed of the same delivery particle λ-STFV10h 1A2 gpJ packaging a modified version of the *cas12* nuclease with similar activity to the one in EB003 were homogenized in 450 µL of PBS and ELISA was performed according to the manufacturer’s instructions. All infant rabbit experiments were performed with EB003 manufactured and purified by Jafral (Slovenia).

### Statistical Analysis

Figure representations and statistical analyses, when needed, are presented for each figure in the corresponding legend. Mann-Whitney tests or unpaired t-tests were performed for Figure 6. Statistical analyses were performed with GraphPad Prism version 10.3.1.

## Acknowledgements

Figures 1A, 3B, 5A and 6A were created with BioRender.com

## Contributions

Conceptualization: MG, AK, FJF, LHC, IS, SP, RT, AKB, AD, XD, DB, JFR

Methodology: MG, AK, FJF, LHC, IS, SP, MA, RT, AKB, CP, GP, DS, BB, KM, JH, PCR, OK, EL, AD, CB, GS, DJG, AL, XD, JFR

Software: SP, BB, KM, JH

Validation: MG, AK, FJF, LHC, IS, MA, RT, AKB, CP, GP, DS, PCR, CB, GS, AL, DB, JFR

Formal analysis: MG, AK, FJF, LHC, SP, MA, RT, AKB, CP, GP, DS, BB, KM, JH, PCR, JFR

Investigation: MG, AK, FJF, LHC, SP, MA, RT, AKB, CP, GP, DS, PCR, OK, EL, AD, CB, GS, DJG, AL, JFR

Resources: MG, AK, FJF, LHC, IS, SP, MA, RT, AKB, CP, GP, DS, PCR, MR, OK, EL, CB, AG, DB, JFR

Data Curation: MG, AK, FJF, LHC, SP, MA, RT, AKB, CP, GP, DS, PCR, JFR

Writing – original draft: MG, FJF, LHC, IS, SP, AKB, DB, JFR

Writing: - review and editing: MG, AK, FJF, LHC, IS, SP, AKB, XD, DB, JFR

Visualization: MG, FJF, LHC, SP, RT, AKB, JFR

Supervision: MG, IS, MR, AD, AG, EMH, XD, DB, JFR

Project administration: IS, MR, AG, EMH, XD, DB, JFR

Funding acquisition: IS, XD

Author contributions are listed following CRediT guidelines (https://www.elsevier.com/researcher/author/policies-and-guidelines/credit-author-state ment).

## Data and materials availability

All data are available in the main text or the supplementary materials

## List of supplementary materials

- **Supplementary Materials and Methods**
- **Supplementary Figure 1. STF chimera host range in O157 strain collection.**
- **Supplementary Figure 2. Targeting of *stx*-contaning O157 strains with engineered cosmids**
- **Supplementary Figure 3. Cumulative killing of a collection of O157*stx* strains**
- **Supplementary Figure 4. Stability of mouse gut colonization by a O157 STEC strain**
- **Supplementary Figure 5. Histopathological analysis of rabbit colon and ileum samples**
- **Supplementary Figure 6. Evolution of STEC-infected infant rabbit weight after treatment initiation**
- **Supplementary Figure 7. Detection of Shiga toxin in rabbit feces samples**
- **Supplementary Figure 8. Multiple sequence alignment of the unique stx gene variants in the database**
- **Supplementary Figure 9. Killing activity of EB003 in the full O157 *stx* strain collection**
- **Supplementary Table 1. gpJ and STF variants**
- **Supplementary Table 2. Strains used in this study**
- **Supplementary Table 3. O157 restriction-modification motifs**
- **Supplementary Table 4. Isolates and genotypes of O157 strains used to find guide RNAs targeting all *stx* gene variants**
- **Supplementary Table 5. Guide RNAs found in the multiplex array**
- **Supplementary Table 6. Composition of in-house O157 collections**
- **Supplementary Table 7. Plasmids used in this study**
- **Supplementary Table 8. Percentage of *stx* gene variants and clinical isolates targeted by EB003 *in silico*.**
- **Supplementary References**

