## Supplementary material for "Treatment of STEC infection via CRISPR-Cas targeted cleavage of the Shiga toxin gene in animal models"

### Supplementary Materials and Methods

#### Strains and media

The *E. coli* and λ prophage genomes were engineered using a strategy explained in *(1)*. Packaged cosmids were produced using an engineered *E. coli* K-12 strain carrying the thermosensitive cI857 λ prophage with its *cos* site deleted and the SrpR repressor inserted in the genome *(2)*. The chimeric *stf* genes were expressed *in trans* with the *stf* gene deleted from the prophage. For experiments with plasmid p1392, the λ STF-V10h chimera was inserted into the prophage genome. All experiments were performed with cells grown in LB plus 5 mM $CaCl_2$, supplemented with antibiotics when necessary (chloramphenicol (12.5 to 25 μg $ml^{-1}$), kanamycin (50 μg $ml^{-1}$)). 2,4-Diacetylphloroglucinol (DAPG; 100 μM, Santa Cruz Biotechnology) was added in order to induce the pPhlF promoter *(2)*. All titration assays for the STF chimera screening assay were carried out with *E. coli* s14269. Titrations of EB003 with a *thyA* marker were performed in strain s17466 with plasmid p1399. For delivery efficiency assays, superfolder Green Fluorescent Protein (sfGFP; GenBank no. AYN72676) encoded in plasmid p513 was used. For experiments in Figures 1 and 2, O157Δ*stx* strains s2185 or s17465 were used. The K12 strain is strain s14269.

#### Cloning and plasmid construction

DNA cloning was performed with chemically competent DH10B cells (Thermo Scientific) or in strains s2072 or s15816 using Gibson Assembly*(3)*. Cas12a and *thyA* (p1392) were codon-optimized for *E. coli* and synthesized (Twist Bioscience). Restriction site removal for the p513 plasmid or the primase-dependent origin of replication (p1392) was done by inserting single mutations in O157 restriction sites and testing functionality. gpJ variants were generated as in *(4)*. The primase gene from the *E. coli* CFT073 strain (NCBI no. AAN79964, locus AE016759_238) was amplified from the CFT073 genome and cloned into plasmid p1400 (constitutive promoter). The cohesive end site (*cos*) of the λ genome was cloned onto plasmids designed to be packed into λ cosmid particles. All plasmids were purified using a Plasmid DNA Miniprep Kit (Omega Bio-Tek) and sequence-verified by Sanger sequencing (Eurofins Genomics, Microsynth).

#### Packaged cosmid production

For the packaged cosmids in Fig. 1b, the production strains contained the plasmids p513 and different plasmids encoding λ STF chimeras; for the packaged cosmids in Fig. 1c, the production strains carried the plasmids p513 and p521, p522 or p363 or p513 only (Ur-λ). In Figure 2b and 2c, the production strains contained plasmid p513 and the λ STF-V10 inserted into the genome or deleted. For the packaged cosmids in Figure 2e, the production strains contained plasmid p513 and p363, p871 or p3606. Strains in Figure 3c contained plasmids p766, p767, p768, p840 or p764. In Figures 3, 4, 5 and 6 production strains contained packaged cosmid p1392 and plasmid p1400 produced in strain s15816.

**Packaged cosmid delivery efficiency**

Delivery efficiency into the strains s17465 or s14269 was analyzed using cosmid particles packaging p513 and equipped with λ STF-74, λ STF-86 or λ STF-V10 and gpJ 1A2. Cells were grown in LB supplemented with 5 mM of $CaCl_2$ to an $OD_{600}$ of 0.2 to 0.6. Cell density was adjusted to $OD_{600}$ 0.025 in fresh LB supplemented with 5 mM $CaCl_2$, and 90 µl of cell culture was mixed with 10 µl of each cosmid serially diluted in LB plus 5 mM $CaCl_2$ to reach different MOIs. The samples were incubated for 45 min at 37ºC and 8 µl were added to 250 µl ice-cold PBS plus 1 mg $ml^{-1}$ kanamycin prior to analysis by flow cytometry (excitation: 488 nm, emission: 530/30 BP; Attune NxT Thermo Scientific).

**Pancreatin resistance assay**

Packaged cosmid particles were diluted 1:100 in 10 mL simulated intestinal fluid (6.8 g $L^{-1}$ $KH_2PO_4$ at pH 6.8 with 10 mg $mL^{-1}$ pancreatin from porcine pancreas, Sigma Aldrich) or PBS buffer as a control. The mixture was incubated at 37ºC for 3 hours at 180 rpm. After incubation, the samples were titrated on s14269 or s17465 to determine functional particles.

**OmpC analysis of O157:H7 strains**

The OmpC protein sequence was extracted from 121 *E. coli* O157 genome assemblies using TBLASTN v.2.11.0 *(5)* with a curated set of OmpC sequences as query. The sequences were aligned using MAFFT *(6)* v.7.520 with default parameters.

### Supplementary Figures

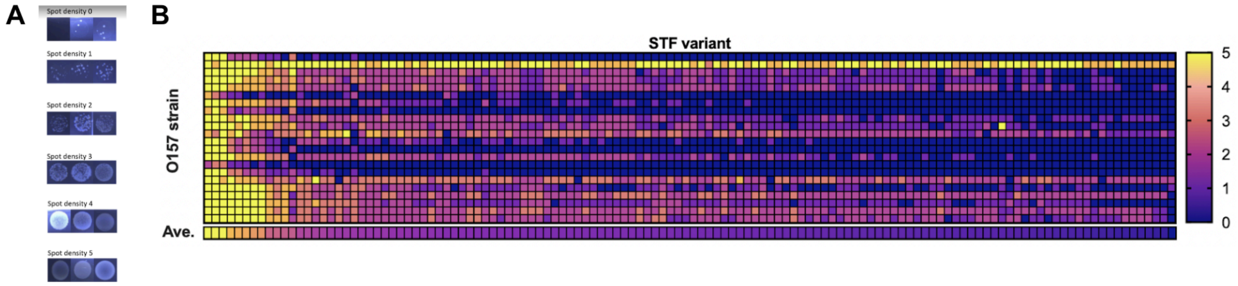

**Supplementary Figure 1. STF chimera host range in O157 strain collection.** A) Qualitative spot density key used for visual determination of delivery efficiency in spotting assays in Fig. 1B. Qualitative delivery efficiency by agar spot density (from 0 - low to 5- high) of engineered particles carrying the 1A2 gpJ variant and different chimeric STFs (all STF chimeras shown) and plasmid p513. Ave., average qualitative delivery efficiency of 22 strains with each STF chimera

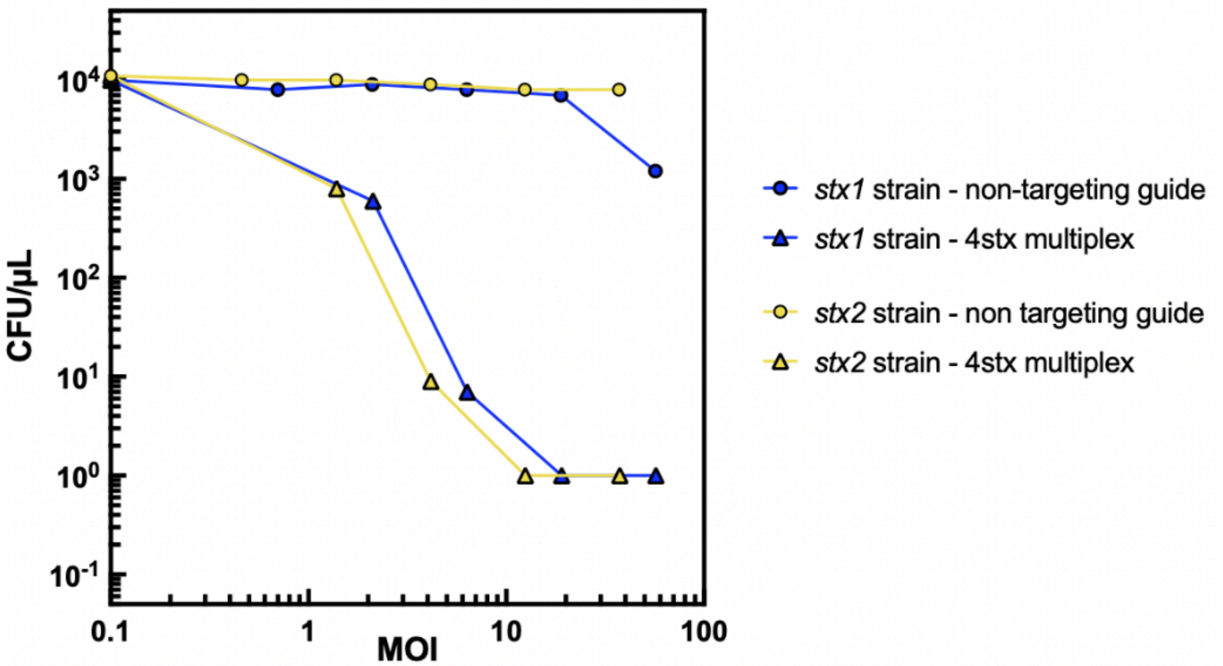

**Supplementary Figure 2. Targeting of *stx*-contaning O157 strains with engineered cosmids**. Engineered cosmid-mediated targeted killing of O157 strains carrying either the *stx1* gene only or the *stx2* gene only at different MOIs in the absence of selection. Blue lines, *stx1* strain; yellow lines, *stx2* strain. Triangles, assay run with the payload shown in Figure 3A, 4stx multiplex; Circles, assay run with a payload containing a non-targeting guide instead of the 4stx multiplex array.

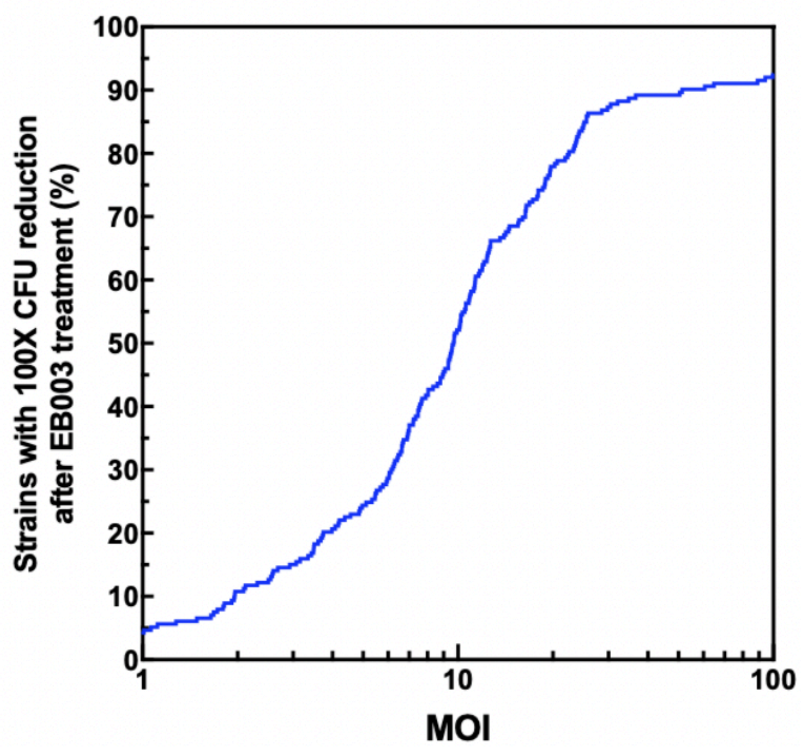

**Supplementary Figure 3. Cumulative killing of a collection of O157*stx* strains.** Percentage of 213 different O157 *stx* strains showing a 100X reduction in CFUs (99%) after treatment with EB003 as a function of MOI.

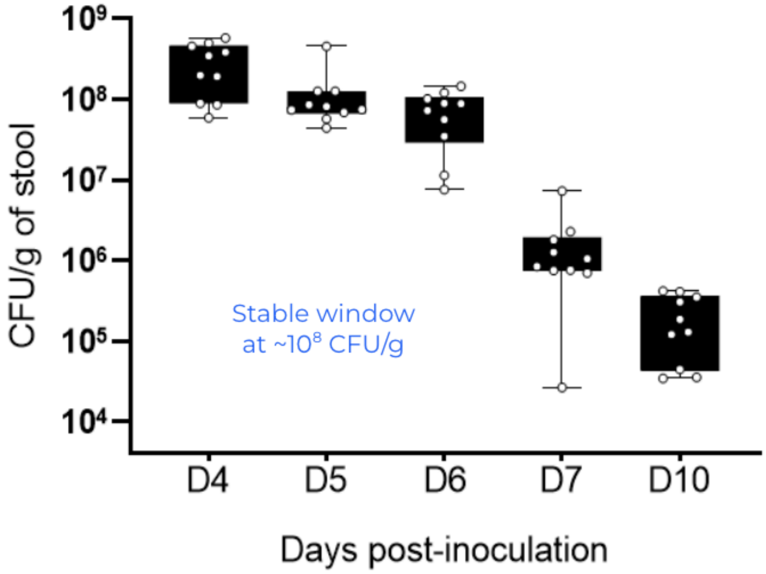

**Supplementary Figure 4. Stability of mouse gut colonization by a O157 STEC strain.** Three days before O157 strain s17769 inoculation, mice were treated with streptomycin 5 g $L^{-1}$ in water. At Day 0, the strain was gavaged and at day 4, streptomycin was removed from the water source. The colonization was monitored by plating resuspended feces on Drigalski media over a period of 10 days. Box extends from the 25th to the 75th percentiles; whiskers show minimum and maximum values.

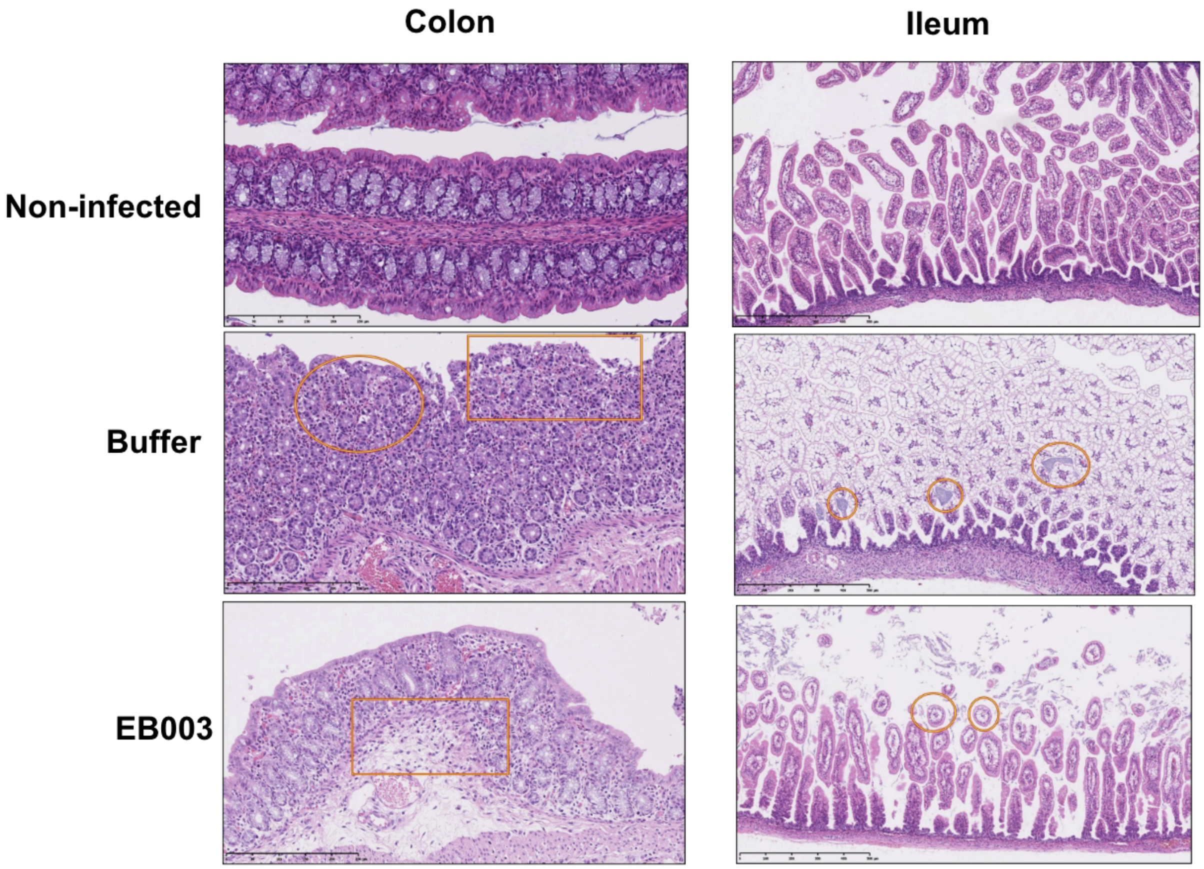

**Supplementary Figure 5. Histopathological analysis of rabbit colon and ileum samples**. Colon and ileum samples from the three groups of animals (non-infected controls treated with buffer, O157*stx* infected and treated with buffer, O157*stx* infected and treated with EB003). Colon samples, top to bottom: normal colon; superficial acute erosive colitis with mild heterophilic inflammation (circle) and focal erosion (rectangle); superficial acute colitis with mild mucosal edema (rectangle) and no epithelial damages. Ileum, from top to bottom: normal ileum; extensive and severe goblet cell hyperplasia with intra-luminal bacterial colonies; minimal goblet cell hyperplasia (circles). Bar represents 250 μm.

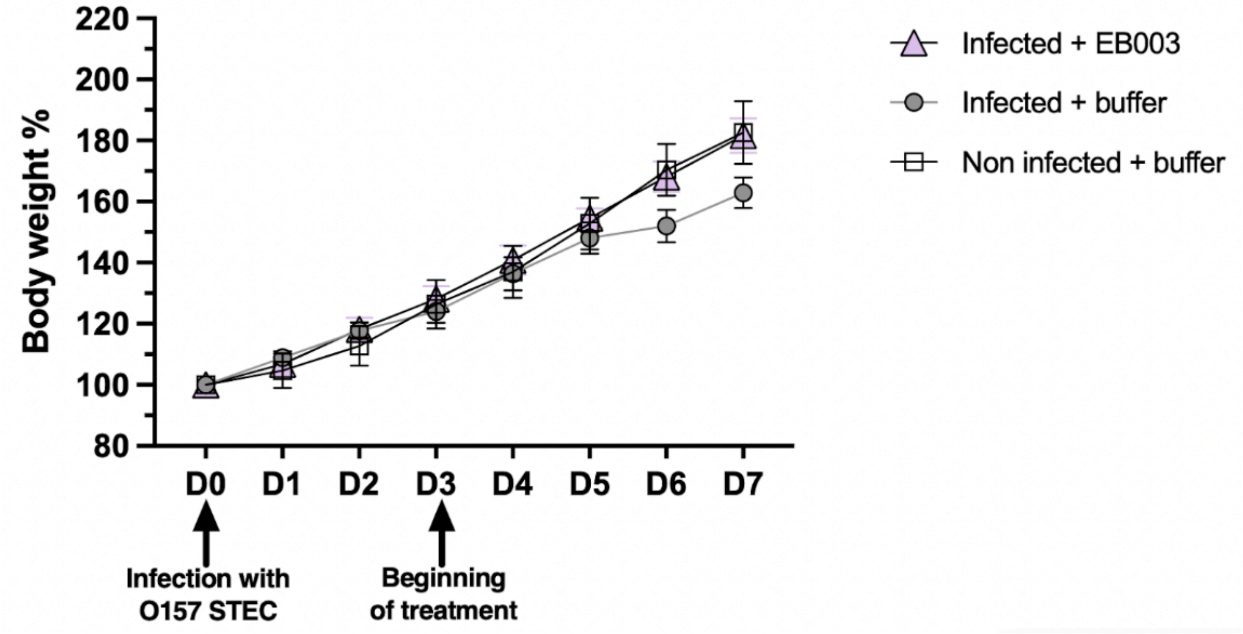

**Supplementary Figure 6. Evolution of STEC-infected infant rabbit weight after treatment initiation** as percentage of weight at Day 0. Plotted: mean and standard error.

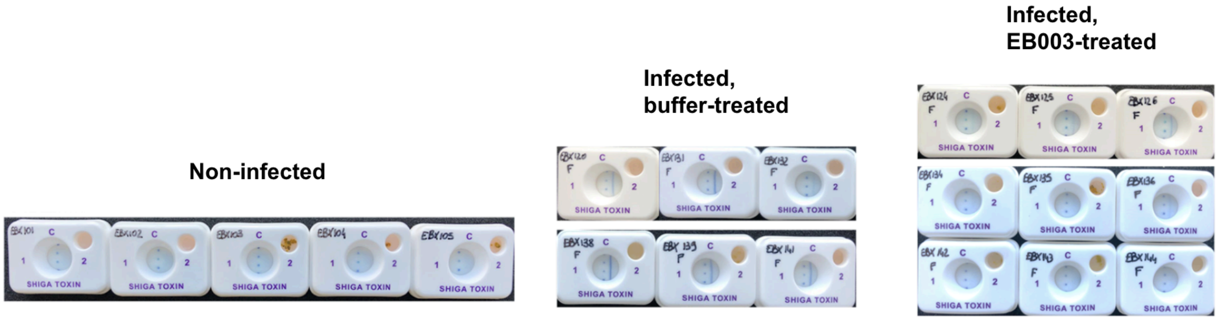

**Supplementary Figure 7. Detection of Shiga toxin in rabbit feces samples**. Fecal samples were resuspended in PBS and applied onto a Shiga Toxin Quick Chek test. The presence of Shiga toxin 2 (right side of the tests, labeled as “2”) can be detected strongly in O157*stx* infected rabbits treated with buffer (middle panel, blue vertical lines), while absent in non-infected rabbits (left panel). Rabbits treated with EB003 show reduced intensity or absence of the shiga toxin 2 band (right panels). The three dots in the center of the tests represent a validation control (i.e. the test was functional). No band is observed for Shiga toxin 1 for any of the samples since the strain used in rabbit models only expresses stx2.

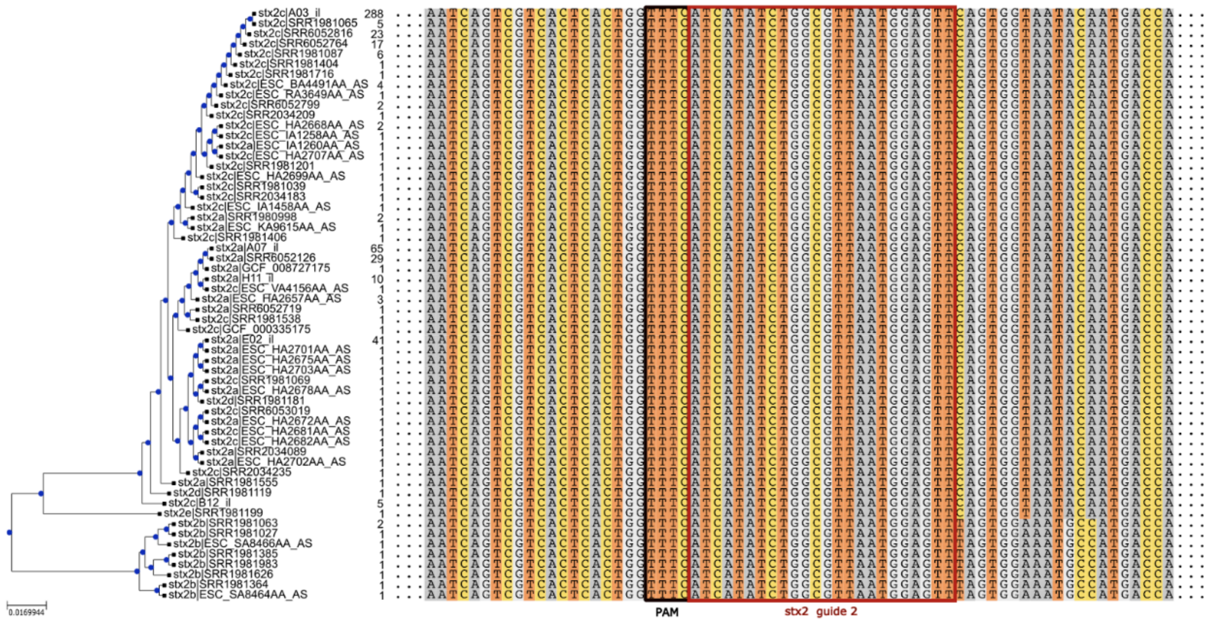

**Supplementary Figure 8. Multiple sequence alignment of the unique stx gene variants in the database**, restricted to the neighborhood of stx2 guide 2. Each representative sequence identifier is followed by the number of isolates it appears in.

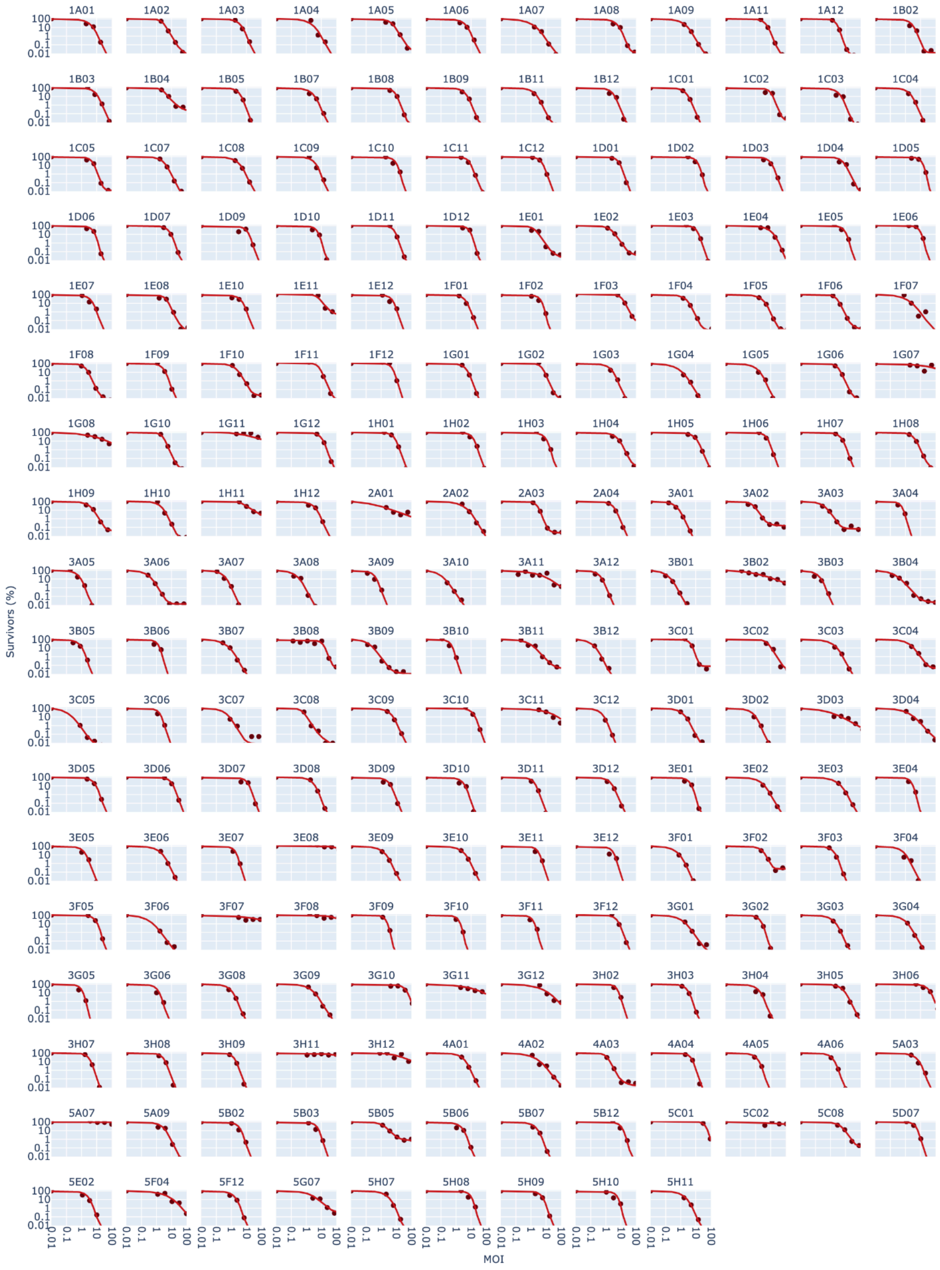

**Supplementary Figure 9. Killing activity of EB003 in the full O157 *stx* strain collection.** Each graph represents the EB003-mediated killing of a single strain (top graph labels) as a percentage of survivors (Y axis) versus MOI (X axis). Liquid bacterial cultures at an $OD_{600}$ of 0.025 were treated with serial 1:3 dilutions of EB003 and incubated at 37ºC for 45 minutes. 10 µL of the incubations were spotted on LB agar plates and incubated for 18 hours at 37ºC. Then, CFUs on agar plates were quantified. Red dots represent CFU counts for a given MOI, red lines are a fitted curve to the killing data (see Methods).

### Supplementary Tables

| STF chimera name | STF chimera plasmid number | STF chimera sequence |
|---|---|---|
| *V10* | p363 | MAVKISGVLKDGTGKPVQNCTIQLKARRNSTTVVVNTVG<br>SENPDEAGRYSMDVEYGQYSVILQVDGFPPSHAGTITV<br>YEDSQPGTLNDFLCAMTEDDARPEVLRRLELMVEEVAR<br>NASVVAQSTADAKKSAGDASASAAQVAALVTDATDSAR<br>AASTSAGQAASSAQEASSGAEAASAKATEAEKSAAAAE<br>SSKNAAATSAGAAKTSETNAAASQQSAATSASTAATKAS<br>EAATSARDAVASKEAAKSSETNASSSAGRAASSATAAEN<br>SARAAKTSETNARSSETAAERSASAAADAKTAAAGSAST<br>ASTKATEAAGSAVSASQSKSAAEAAAIRAKNSAKRAEDI<br>ASAVALEDADTTRKGIVQLSSATNSTSETLAATPKAVKVV<br>MDETNRKAPLDSPALTGTPTAPTALRGTNNTQIANTAFVL<br>AAIADVIDASPDALNTLNELAAALGNDPDFATTMTNALAG<br>KQPKNATLTALAGLSTAKNKLPYFAENDAASLTELTQVGR<br>DILAKNSVADVLEYLGAGENSAANDGFKFIGQCPDILTLR<br>TIEPEKNGQRITLRQHTIGTGLGGGVFRAVLDGTGYTDD<br>DGVVIKTAGGSVWLRVNADKVNPFMFGATGVADDTAAL<br>QKMLECGRAAELGTNVWKASNLELNNKSCSLSGSGLH<br>VSRIEQISGATGALLTITQDCSLIYLSDCGLYGDGITAGTS<br>GVTMETGNPGGAPSYPFNTAPDVRRDLYISNVHITGFDE<br>LGFDYPETNFSVSTHGLFIRNIKKTGAKIGTTDFTWTNLQ<br>IDTCGQECLVLDGAGNCRIIGAKLIWAGSENETPYSGLRI<br>SNSQNVNMTGVELQDCAYDGLYIKNSTVAISGLNTNRNS<br>ASSNLSYHNMVFENSIVTVDGYVCRNYAATSLYDLNSQA<br>GNVRCIGSDSTVLINGIYESEVNSERLMGDNNLIQPYSG<br>DLIINGLKNYYTYTGSVKNNIPTFDGVVTTATYVSAPSILG<br>QGNMLKLTQSNKDKLLFSDKVSRHGCTIGLVLIPSFTGAT<br>TMTAFTLGSGYSPSGNSAVMQFIVNSSGVQTIAILLSGD<br>GITQTLTSDLTTEQALASGGVYHFAMGFAPGRLWWSIIDI<br>NTGRRIRRAYRQPDLHAAFNSIFNSGTSSITAFSGPLAGD<br>IACEGAGSHVYVGGFSSESDYAASRMYGLFTPVDLDKQ<br>YSFRTLNGNINYIVRLA |
| *V10FA* | p871 | MAVKISGVLKDGTGKPVQNCTIQLKARRNSTTVVVNTVG<br>SENPDEAGRYSMDVEYGQYSVILQVDGFPPSHAGTITV<br>YEDSQPGTLNDFLCAMTEDDARPEVLRRLELMVEEVAR<br>NASVVAQSTADAKKSAGDASASAAQVAALVTDATDSAR<br>AASTSAGQAASSAQEASSGAEAASAKATEAEKSAAAAE<br>SSKNAAATSAGAAKTSETNAAASQQSAATSASTAATKAS<br>EAATSARDAVASKEAAKSSETNASSSAGRAASSATAAEN<br>SARAAKTSETNARSSETAAERSASAAADAKTAAAGSAST<br>ASTKATEAAGSAVSASQSKSAAEAAAIRAKNSAKRAEDI<br>ASAVALEDADTTRKGIVQLSSATNSTSETLAATPKAVKVV<br>MDETNRKAPLDSPALTGTPTAPTALRGTNNTQIANTAFVL<br>AAIADVIDASPDALNTLNELAAALGNDPDFATTMTNALAG<br>KQPKNATLTALAGLSTAKNKLPYFAENDAASLTELTQVGR<br>DILAKNSVADVLEYLGAGENSAANDGFAFIGQCPDILTLR<br>TIEPEKNGQRITLRQHTIGTGLGGGVFRAVLDGTGYTDD<br>DGVVIKTAGGSVWLRVNADKVNPFMFGATGVADDTAAL<br>QKMLECGRAAELGTNVWKASNLELNNKSCSLSGSGLH<br>VSRIEQISGATGALLTITQDCSLIYLSDCGLYGDGITAGTS |

| | | |
|---|---|---|
| | | GVTMETGNPGGAPSYPFNTAPDVRRDLYISNVHITGFDE<br>LGFDYPETNFSVSTHGLFIRNIKKTGAKIGTTDFTWTNLQ<br>IDTCGQECLVLDGAGNCRIIGAKLIWAGSENETPYSGLRI<br>SNSQNVNMTGVELQDCAYDGLYIKNSTVAISGLNTNRNS<br>ASSNLSYHNMVFENSIVTVDGYVCRNYAATSLYDLNSQA<br>GNVRCIGSDSTVLINGIYESEVNSERLMGDNNLIQPYSG<br>DLIINGLKNYYTYTGSVKNNIPTFDGVVTTATYVSAPSILG<br>QGNMLKLTQSNKDKLLFSDKVSRHGCTIGLVLIPSFTGAT<br>TMTAFTLGSGYSPSGNSAVMQFIVNSSGVQTIAILLSGD<br>GITQTLTSDLTTEQALASGGVYHFAMGFAPGRLWWSIIDI<br>NTGRRIRRAYRQPDLHAAFNSIFNSGTSSITAFSGPLAGD<br>IACEGAGSHVYVGGFSSESDYAASRMYGLFTPVDLDKQ<br>YSFRTLNGNINYIVRLA |
| *V10AAH* | p3606 | MAVKISGVLKDGTGKPVQNCTIQLKARRNSTTVVVNTVG<br>SENPDEAGRYSMDVEYGQYSVILQVDGFPPSHAGTITV<br>YEDSQPGTLNDFLCAMTEDDARPEVLRRLELMVEEVAR<br>NASVVAQSTADAKKSAGDASASAAQVAALVTDATDSAR<br>AASTSAGQAASSAQEASSGAEAASAKATEAEKSAAAAE<br>SSKNAAATSAGAAKTSETNAAASQQSAATSASTAATKAS<br>EAATSARDAVASKEAAKSSETNASSSAGRAASSATAAEN<br>SARAAKTSETNARSSETAAERSASAAADAKTAAAGSAST<br>ASTKATEAAGSAVSASQSKSAAEAAAIRAKNSAKRAEDI<br>ASAVALEDADTTRKGIVQLSSATNSTSETLAATPKAVKVV<br>MDETNRKAPLDSPALTGTPTAPTALRGTNNTQIANTAFVL<br>AAIADVIDASPDALNTLNELAAALGNDPDFATTMTNALAG<br>KQPKNATLTALAGLSTAKNKLPYFAENDAASLTELTQVGR<br>DILAKNSVADVLEYLGAGENSAANDGAAHIGQCPDILTLR<br>TIEPEKNGQRITLRQHTIGTGLGGGVFRAVLDGTGYTDD<br>DGVVIKTAGGSVWLRVNADKVNPFMFGATGVADDTAAL<br>QKMLECGRAAELGTNVWKASNLELNNKSCSLSGSGLH<br>VSRIEQISGATGALLTITQDCSLIYLSDCGLYGDGITAGTS<br>GVTMETGNPGGAPSYPFNTAPDVRRDLYISNVHITGFDE<br>LGFDYPETNFSVSTHGLFIRNIKKTGAKIGTTDFTWTNLQ<br>IDTCGQECLVLDGAGNCRIIGAKLIWAGSENETPYSGLRI<br>SNSQNVNMTGVELQDCAYDGLYIKNSTVAISGLNTNRNS<br>ASSNLSYHNMVFENSIVTVDGYVCRNYAATSLYDLNSQA<br>GNVRCIGSDSTVLINGIYESEVNSERLMGDNNLIQPYSG<br>DLIINGLKNYYTYTGSVKNNIPTFDGVVTTATYVSAPSILG<br>QGNMLKLTQSNKDKLLFSDKVSRHGCTIGLVLIPSFTGAT<br>TMTAFTLGSGYSPSGNSAVMQFIVNSSGVQTIAILLSGD<br>GITQTLTSDLTTEQALASGGVYHFAMGFAPGRLWWSIIDI<br>NTGRRIRRAYRQPDLHAAFNSIFNSGTSSITAFSGPLAGD<br>IACEGAGSHVYVGGFSSESDYAASRMYGLFTPVDLDKQ<br>YSFRTLNGNINYIVRLA |
| *V10h* | p872 | MAVKISGVLKDGTGKPVQNCTIQLKARRNSTTVVVNTVG<br>SENPDEAGRYSMDVEYGQYSVILQVDGFPPSHAGTITV<br>YEDSQPGTLNDFLCAMTEDDARPEVLRRLELMVEEVAR<br>NASVVAQSTADAKKSAGDASASAAQVAALVTDATDSAR<br>AASTSAGQAASSAQEASSGAEAASAKATEAEKSAAAAE<br>SSKNAAATSAGAAKTSETNAAASQQSAATSASTAATKAS<br>EAATSARDAVASKEAAKSSETNASSSAGRAASSATAAEN<br>SARAAKTSETNARSSETAAERSASAAADAKTAAAGSAST<br>ASTKATEAAGSAVSASQSKSAAEAAAIRAKNSAKRAEDI |

| | | |
|---|---|---|
| | | ASAVALEDADTTRKGIVQLSSATNSTSETLAATPKAVKVV<br>MDETNRKAPLDSPALTGTPTAPTALRGTNNTQIANTAFVL<br>AAIADVIDASPDALNTLNELAAALGNDPDFATTMTNALAG<br>KQPKNATLTALAGLSTAKNKLPYFAENDAASLTELTQVGR<br>DILAKNSVADVLEYLGAGENSGSATDVMIQLAANDGFKFI<br>GQCPDILTLRTIEPEKNGQRITLRQHTIGTGLGGGVFRAV<br>LDGTGYTDDDGVVIKTAGGSVWLRVNADKVNPFMFGAT<br>GVADDTAALQKMLECGRAAELGTNVWKASNLELNNKSC<br>SLSGSGLHVSRIEQISGATGALLTITQDCSLIYLSDCGLYG<br>DGITAGTSGVTMETGNPGGAPSYPFNTAPDVRRDLYISN<br>VHITGFDELGFDYPETNFSVSTHGLFIRNIKKTGAKIGTT<br>DFTWTNLQIDTCGQECLVLDGAGNCRIIGAKLIWAGSEN<br>ETPYSGLRISNSQNVNMTGVELQDCAYDGLYIKNSTVAIS<br>GLNTNRNSASSNLSYHNMVFENSIVTVDGYVCRNYAAT<br>SLYDLNSQAGNVRCIGSDSTVLINGIYESEVNSERLMGD<br>NNLIQPYSGDLIINGLKNYYTYTGSVKNNIPTFDGVVTTAT<br>YVSAPSILGQGNMLKLTQSNKDKLLFSDKVSRHGCTIGL<br>VLIPSFTGATTMTAFTLGSGYSPSGNSAVMQFIVNSSGV<br>QTIAILLSGDGITQTLTSDLTTEQALASGGVYHFAMGFAP<br>GRLWWSIIDINTGRRIRRAYRQPDLHAAFNSIFNSGTSSI<br>TAFSGPLAGDIACEGAGSHVYVGGFSSESDYAASRMYG<br>LFTPVDLDKQYSFRTLNGNINYIVRLA |
| *STF74* | p521 | MAVKISGVLKDGTGKPVQNCTIQLKARRNSTTVVVNTVG<br>SENPDEAGRYSMDVEYGQYSVILQVDGFPPSHAGTITV<br>YEDSQPGTLNDFLCAMTEDDARPEVLRRLELMVEEVAR<br>NASVVAQSTADAKKSAGDASASAAQVAALVTDATDSAR<br>AASTSAGQAASSAQEASSGAEAASAKATEAEKSAAAAE<br>SSKNAAATSAGAAKTSETNAAASQQSAATSASTAATKAS<br>EAATSARDAVASKEAAKSSETNASSSAGRAASSATAAEN<br>SARAAKTSETNARSSETAAERSASAAADAKTAAAGSAST<br>ASTKATEAAGSAVSASQSKSAAEAAAIRAKNSAKRAEDI<br>ASAVALEDADTTRKGIVQLSSATNSTSETLAATPKAVKVV<br>MDETNRKYTAQDASTAQKGLVKLSSATDSTSETLAATPK<br>AVKAVNDNANGRVPSERKVNGHSLAGDISVTSQDIFDG<br>QCVEIGPGQDLDNYQTPGLYFQPANANTSAALHYPENN<br>AGSLMVLRSAGITQVYRVYSGSRSYLRSKYSTQPWTTW<br>TPDDAFPVGAPIPWPSDIAPPAYALMQGQSFDKSAYPLL<br>AVAYPSGVIPDMRGQTIKGKPDGRAVLSYEQDGIKSHAH<br>TASISDTDLGTKYTNSFDYGSKPTTSFDYGNKSSTEGG<br>WHVHNFRYCATSAYRDTPGSGLGMHSSNISWSAGDRIE<br>GSGNHAHVTWIGPHDHWVGIGEHNHYVVMGYHGHTAT<br>VHATGNTENTVKNIAFNYIVRLA |
| *STF86* | p522 | MAVKISGVLKDGTGKPVQNCTIQLKARRNSTTVVVNTVG<br>SENPDEAGRYSMDVEYGQYSVILQVDGFPPSHAGTITV<br>YEDSQPGTLNDFLCAMTEDDARPEVLRRLELMVEEVAR<br>NASVVAQSTADAKKSAGDASASAAQVAALVTDATDSAR<br>AASTSAGQAASSAQEASSGAEAASAKATEAEKSAAAAE<br>SSKNAAATSAGAAKTSETNAAASQQSAATSASTAATKAS<br>EAATSARDAVASKEAAKSSETNASSSAGRAASSATAAEN<br>SARAAKTSETNARSSETAAERSASAAADAKTAAAGSAST<br>ASTKATEAAGSAVSASQSKSAAEAAAIRAKNSAKRAEDI<br>ASAVALEDADTTRKGIVQLSSATNSTSETLAATPKAVKVV<br>MDETNRRVPASRKVNGHALNGDINVTSRDIFDGQVIAIG |

| | | |
|---|---|---|
| | | ANKNLDDYQVPGLYFQEANNNTSAAMNYPENSAGSLM<br>VLRGAGVTQVYRVYNSSRSYSRSKYSTLAWTPWMPED<br>SYPVGAPIPWPSDVTPTGYALMQGQPFDKAVYPLLAIAY<br>PAGIIPDMRGQTIKGKPNGRAVLSYEQDGVISHTHGASIS<br>DTDLGTKYTSSFDYGSKPTTSFDYGNKSSTEGGWHAH<br>NFRYCATSAYRDTPGQGLGMHSSNVSWAAGDRIEGSG<br>NHAHVTWIGPHDHWVGIGAHNHYVVMGYHGHTATVHA<br>AGNAENTVKNIAFNYIVRLA |
| WT Lambda STF | p515 | MAVKISGVLKDGTGKPVQNCTIQLKARRNSTTVVVNTVG<br>SENPDEAGRYSMDVEYGQYSVILQVDGFPPSHAGTITV<br>YEDSQPGTLNDFLCAMTEDDARPEVLRRLELMVEEVAR<br>NASVVAQSTADAKKSAGDASASAAQVAALVTDATDSAR<br>AASTSAGQAASSAQEASSGAEAASAKATEAEKSAAAAE<br>SSKNAAATSAGAAKTSETNAAASQQSAATSASTAATKAS<br>EAATSARDAVASKEAAKSSETNASSSAGRAASSATAAEN<br>SARAAKTSETNARSSETAAERSASAAADAKTAAAGSAST<br>ASTKATEAAGSAVSASQSKSAAEAAAIRAKNSAKRAEDI<br>ASAVALEDADTTRKGIVQLSSATNSTSETLAATPKAVKVV<br>MDETNRKAPLDSPALTGTPTAPTALRGTNNTQIANTAFVL<br>AAIADVIDASPDALNTLNELAAALGNDPDFATTMTNALAG<br>KQPKNATLTALAGLSTAKNKLPYFAENDAASLTELTQVGR<br>DILAKNSVADVLEYLGAGENSAFPAGAPIPWPSDIVPSGY<br>VLMQGQAFDKSAYPKLAVAYPSGVLPDMRGWTIKGKPA<br>SGRAVLSQEQDGIKSHTHSASASGTDLGTKTTSSFDYG<br>TKTTGSFDYGTKSTNNTGAHAHSLSGSTGAAGAHAHTS<br>GLRMNSSGWSQYGTATITGSLSTVKGTSTQGIAYLSKTD<br>SQGSHSHSLSGTAVSAGAHAHTVGIGAHQHPVVIGAHA<br>HSFSIGSHGHTITVNAAGNAENTVKNIAFNYIVRLA |
| Lambda deltaSTF | --- | |

| **gpJ chimera name** | **gpJ chimera sequence** |
|---|---|
| **1A2** | MGKGSSKGHTPREAKDNLKSTQLLSVIDAISEGPIEGPVDGLKSVLLNSTPVLDTEGNTNIS<br>GVTVVFRAGEQEQTPPEGFESSGSETVLGTEVKYDTPITRTITSANIDRLRFTFGVQALVET<br>TSKGDRNPSEVRLLVQIQRNGGWVTEKDITIKGKTTSQYLASVVMGNLPPRPFNIRMRRMT<br>PDSTTDQLQNKTLWSSYTEIIDVKQCYPNTALVGVQVDSEQFGSQQVSRNYHLRGRILQVP<br>SNYNPQTRQYSGIWDGTFKPAYSNNMAWCLWDMLTHPRYGMGKRLGAADVDKWALYVIG<br>QYCDQSVPDGFGGTEPRITCNAYLTTQRKAWDVLSDFCSAMRCMPVWNGQTLTFVQDRP<br>SDKTWTYNRSNVVMPDDGAPFRYSFSALKDRHNAVEVNWIDPNNGWETATELVEDTQAIA<br>RYGRNVTKMDAFGCTSRGQAHRAGLWLIKTELLETQTVDFSVGAEGLRHVPGDVIEICDD<br>DYAGISTGGRVLAVNSQTRTLTLDREITLPSSGTALISLVDGSGNPVSVEVQSVTDGVKVKV<br>SRVPDGVAEYSVWELKLPTLRQRLFRCVSIRENDDGTYAITAVQHVPEKEAIVDNGAHFDG<br>EQSGTVNGVTPPAVQHLTAEVTADSGEYQVLARWDTPKVVKGVSFLLRLTVTADDGSERLV<br>STARTTETTYRFTQLALGNYRLTVRAVNAWGQQGDPASVSFRIAAPAAPSRIELTPGYFQIT<br>ATPHLAVYDPTVQFEFWFSEKQIADIRQVETSTRYLGTALYWIAASINIKPGHDYYFYIRSVN<br>TVGKSAFVEAVGRASDDAEGYLDFFKGKITESHLGKELLEKVELTEDNASRLEEFSKEWKD<br>ASDKWNAMWAVKIEQTKDGKHYVAGIGLSMEDTEEGKLSQFLVAANRIAFIDPANGNETPM<br>FVAQGNQIFMNDVFLKRLTAPTITSGGNPPAFSLTPDGKLTAKNADISGNVNANSGTLNNVTI<br>NENCRVLGKLSANQIEGDLVKTVGKAFPRDSRAPERWPSGTITVRVYDDQPFDRQIVIPAVA<br>FSGAKHEKEHTDIYSSCRLIVRKNGAEIYNRTALDNTLIYSGVIDMPAGHGHMTLEFSVSAW<br>LVNNWYPTASISDLLVVVMKKATAGITIS |

#### Supplementary Table 1. gpJ and STF variants

| Strain name | Genotype/Description | Supplier |
|---|---|---|
| s17465 | O157:H7 / Gene_stx : none / MLST : ST11 | This study |
| s14269 | *K-12 F- λ- ilvG- rfb-50 rph-1 rpsLK42R ompC-EDL933* | This study |
| s17466 | O157:H7 / Gene_stx : none / MLST : ST11 / *ΔthyA* | This study |
| s2072 | *BW25113 Δ9 λcI857 Sam7 Δcos gpJ-1A2 ∆stf lacZ::SrpR lamB-* | This study |
| s15816 | *BW25113 Δ9 λcI857 Sam7 Δcos gpJ-1A2 stf::V10h lacZ::SrpR lamB- ΔthyA* | This study |
| s2185 | O157:H7 / Gene_stx : none / MLST : ST11 / *rpsLK42R* | This study |
| s1592 | O157:H7 / Gene_stx : stx1a / MLST : ST11 | Pasteur Institute |
| s1594 | O157:H7 / Gene_stx : stx2a / MLST : ST11 | Pasteur Institute |
| s13699 | O157:H7 / Gene_stx: stx2a / ST11 / Phage Type 21/28 | PHE |
| s13729 | O157:H7 / Gene_stx: stx2a / ST11 / Phage Type 21/28 | PHE |
| s13791 | O157:H7 / Gene_stx: stx2a stx2c / ST11 / Phage Type 31 | PHE |
| s13715 | O157:H7 / Gene_stx: stx2a stx2c / ST11 / Phage Type 21/28 | PHE |
| s13765 | O157:H7 / Gene_stx: stx2a stx2c / ST11 / Phage Type 32 | PHE |
| s13734 | O157:H7 / Gene_stx: stx2c stx1a / ST11 / Phage Type 8 | PHE |
| s13732 | O157:H7 / Gene_stx: stx2c stx1a / ST11 / Phage Type 8 | PHE |
| s13760 | O157:H7 / Gene_stx: stx2a stx1a / ST11 / Phage Type 32 | PHE |
| s17769 | O157:H7 / Gene_stx : stx2a / MLST : ST11 / *rpsLK42R* | This study |
| s17773 | O157:H7 / Gene_stx: stx2a stx2c / ST11 / Phage Type 21/28 *rpsLK42R* | This study |
| DH10B | *F– mcrA Δ(mrr-hsdRMS-mcrBC) φ80lacZΔM15 ΔlacX74 recA1 endA1 araD139 Δ (ara-leu)7697 galU galK λ– rpsL(StrR) nupG* | Thermo Scientific |

**Supplementary Table 2. Strains used in this study**

| Note: the motif CCWGG is also found in some O157 strains which corresponds to the housekeeping dcm methylase | | | |
|---|---|---|---|
| **Strain Name** | **Motif 1** | **Motif 2** | **Motif 3** |
| 180-PT54 | CACNNNNNNNCTGG | GAAABCC | CCAYNNNNNGTTY |
| 644-PT8 | CACNNNNNNNCTGG | GAAABCC | CCAYNNNNNGTTY |
| FDAARGOS_293 | CACNNNNNNNCTGG | GAAABCC | |
| 1130 | CACNNNNNNNCTGG | GAAABCC | |
| 2149 | CACNNNNNNNCTGG | GAAABCC | |
| 2159 | CACNNNNNNNCTGG | GAAABCC | |
| 3384 | CACNNNNNNNCTGG | GAAABCC | |
| 4276 | CACNNNNNNNCTGG | GAAABCC | |
| 8368 | CACNNNNNNNCTGG | GAAABCC | |
| 9234 | CACNNNNNNNCTGG | GAAABCC | |
| EDL933 | CACNNNNNNNCTGG | GAAABCC | |

**Supplementary Table 3. Restriction-modification motifs found on REBASE PacBio for *E. coli* O157 strains**

| | | | | Total number of: | |
|---|---|---|---|---|---|
| Data source | Year of isolation | Country of origin (number of isolates) | Genotypes (number of isolates) | Isolates | stx genes |
| EnteroBase<br>NCBI<br>RefSeq<br>PHE<br>Institut Pasteur | 2015-2019 | UK (1113)<br>US (292)<br>France (128)<br>Austria (12)<br>Luxemburg (8)<br>Canada (5)<br>Norway (1)<br>South Korea (1) | stx2c (582)<br>stx1a+stx2c (314)<br>stx2a (246)<br>stx1a (106)<br>stx1a+stx2a (26)<br>stx2b (6)<br>stx1c (4)<br>stx1c+stx2b (2)<br>stx1a+stx2b (2)<br>stx2d (1)<br>stx1a+stx2d (1)<br>stx2e (1)<br>stx1d (1) | 1292 | 1640 |

**Supplementary Table 4. Isolates and genotypes of O157 strains used to find guide RNAs targeting all *stx* gene variants**

| Guide name | Guide sequence |
|---|---|
| stx2 - guide 1 | CATTCCGGAACGTTCCAGCGCTGC |
| stx2 - guide 2 | ATCATATCTGGCGTTAATGGAGTT |
| stx1 - guide 1 | CTGATTTTTCACATGTTACCTTTC |
| stx1 - guide 2 | ATCATATCTGGCGTTAATGGAGTT |

**Supplementary Table 5. Guide RNAs found in the multiplex array**

| Collection | Total Isolates | stx2a | stx2c | stx2d | stx2a stx2c | stx2a stx1a | stx2c stx1a | stx2a stx2c stx1a | stx1a |
|---|---|---|---|---|---|---|---|---|---|
| Pasteur CNR V1 | 22 | 7 | 6 | 0 | 0 | 1 | 7 | 0 | 1 |
| Pasteur CNR V2 | 102 | 22 | 29 | 2 | 7 | 18 | 23 | 0 | 1 |
| PHE | 100 | 41 | 0 | 0 | 32 | 1 | 22 | 2 | 2 |
| **Total** | 224 | 70 | 35 | 2 | 39 | 20 | 52 | 2 | 4 |
| % | | **31%** | **16%** | **1%** | **17%** | **9%** | **23%** | **1%** | **2%** |
| | | **Contains stx2a** | **Contains stx2c** | **Contains stx2d** | **Contains stx1a** | **Contains Stx1 and Stx2** | | | |
| | Total | 131 | 128 | 2 | 78 | 74 | | | |
| | % | 58% | 57% | 1% | 35% | 33% | | | |

**Supplementary Table 6. Composition of in-house O157 collections**

| Name | Sequence |
| --- | --- |
| p513 | CCTTTAGGGAAATATGCTAAGTTTTCACCGTAACACGCCACATCTTGACTATATATGTGTAGA<br>AACTGCCGGAAATCGTCGTGGTATTCTGACCAGAGCGATGAAAACGTTTCAGTTTGCTCAT<br>GGAAAACGGTGTAACAAGGGTGAACACTATCCCATATCACCAGCTCACCGTCTTTCATTGC<br>CATACGAAACTCCGGATGTGCATTCATCAGGCGGGCAAGAATGTGAATAAAGGCCGGATAA<br>AACTTGTGCTTATTTTTCTTTACGGTTTTTAAAAAGGCCGTAATATCCAGCTGAACGGTTTGG<br>TTATAGGTGCACTGAGCAACTGACTGGAATGCCTCAAAATGTTCTTTACGATGCCATTGACT<br>TATATCAACTGTAGTATATCCAGTGATTTTTTTCTCCATTTTAGCTTCCTTAGCTTGCGAAATC<br>TCGATAACTCAAAAAATAGTAGTGATCTTATTTCATTATGGTGAAAGTTGTCTTACGTGCAAC<br>ATTTTCGCAAAAAGTTGGCGCTTTATCAACACTGTCCCTCCTGTTCAGCTACTGACGGTACT<br>GCGGAACTGACTAAAGTAGTGCGTAACGGCAAAAGCACCGCCGGACATCTGCGCTAGCG<br>GAGTGTATACTGGCTTACTATGTTGGCACTGATGAGGGTGTAAGTGAAGTGCTTCATGTGG<br>CAGGAGAAAAAAGGCTGCATCGGTGCGTCAGCAGAATATGTGATACAGGATATATTCCGCT<br>TCCTCGCTCACTGACTCGCTACGCTCGGTCGTTCGACTGTGGCGAGCGGAAATGGCTTAC<br>GAACGGGGCGGAGATTTCCTGGAAGATGCCAGGAAGATACTTAACAGGGAAGTGAGAGG<br>GTCGCGGCAAAGCCGTTTTTCCATAGGCTCCGCCCCCCTGACAAGCATCACGAAATCTGA<br>CGCTCAAATCAGTGGTGGCGAAACCTGACAGGACTATAAAGATACCAGGCGTTTCCCCCT<br>GGCGGCTCCCTCGTGCGCTCTCCTGTTCCTGCCTTTCGGTTTGCCGGTGTCATTCCTCTG<br>TTACGGCCGAGTTTGTCTCATTCCACGCCTGACACTCAGTTCCGGGTAGGCAGTTCGCTC<br>CAAGCTGGACTGTATGCACGAACCCCCCGTTCAGTCCGACCGCTGCGCCTTATCCGGTAA<br>CTATCGTCTTGAGTCCAACCCGGAAAGACATGCAAAAGCACCACTGGCAGCAGCCACTGG<br>TAATTGATTTAGAGGAGTTAGTCTTGAAGTCATGCGCCGGATAAGGCTAAACTGAAAGGAC<br>AAGTTTTGGCGACTGCGCTCCTCCAAGCCAGTTACCTCGGTTCAAAGAGTTGGTAGCTCA<br>GAGAACCTTCGAAAAACCGCCCTGCAAGGCGGTTTTTTCGTTTTCAGAGCAAGAGATTAC<br>GCGCAGACCAAAACGATCTCAAGAAGATCATCTTATTAATCAGATAAAATATTTCTAGATTTC<br>AGTGCAATTTATCTCTTCAAATGTAGCACTTTATAGCTAGCTCAGCCCTTGGTACAATGCTAG<br>CGTTTTCATTAAAGAGGAGAAAGGAAGCCATGAGTAAAGGTGAGGAATTATTTACTGGTGTT<br>GTTCCGATCTTAGTTGAACTGGACGGCGATGTTAACGGTCATAAATTCAGTGTTCGTGGTG<br>AAGGTGAAGGTGATGCAACCAACGGTAAGCTGACCCTGAAATTCATCTGCACTACTGGAAA<br>ATTACCAGTACCGTGGCCTACTCTGGTGACTACCCTGACCTATGGTGTTCAGTGTTTTTCTC<br>GTTACCCTGACCACATGAAGCAACATGATTTCTTCAAATCTGCAATGCCGGAAGGTTATGTA<br>CAGGAGCGCACCATTTCTTTCAAAGACGATGGCACGTATAAAACCCGTGCAGAGGTTAAAT<br>TTGAAGGTGACACTCTGGTGAATCGTATTGAACTGAAAGGCATTGATTTCAAAGAGGACGG<br>CAATATTTTAGGCCACAAACTGGAATATAACTTCAACTCCCATAACGTTTACATCACCGCAGA<br>CAAACAAAAGAACGGTATCAAAGCTAACTTCAAAATTCGCCATAACGTTGAAGACGGTAGC<br>GTACAGCTGGCGGATCATTACCAACAGAACACTCCGATTGGAGATGCTCCTGTTTTACTGC<br>CGGATAACCACTACCTGTCCACCCAGTCTAAACTGTCGAAGGATCCGAACGAAAAGCGCG<br>ACCACATGGTGTTATTAGAGTTCGTTACCGCTAGTGGTATCACGCACGGTATGGATGAACTC<br>TACAAATAAGTCAGTTTCACCTGTTTTACGTTAAAACCCGCTTCGGCGGGTTTTTACTTTTG<br>GGTTTAGCCGAACGCCCCAAAAAGCCTCGCTTTCAGCACCTGTCGTTTCCTTTCTTTTCAG<br>AGGGTATTTTAAATAAAAACATTAAGTTATGACGAAGAAGAACGGAAACGCCTTAAACCGGA<br>AAATTTTCATAAATAGCGAAAACCCGCGAGGTCGCCGCCCCGTAACCTGTCGGATCACCG<br>GAAAGAACCTGTAAAGTGATAATGATTATCATCTACATATCACAACGTGCGTAAAGGACTAT<br>AACAAGACGCAAACGGAGGTAGGCTCACTCCTACTTCGGAAACTTAACCGAAGAACTAGG<br>ACGGTATTGTTTGCGCTTGGAATTGGCCTTGAAGTAAGTCAGGTTTTGACGGAACGATTAG<br>TTACAGGGGGGGAACAGTCGTTGGTCGCCACCAAGTCGATTTTTGGCTTACCTCTTATCTC<br>GTAGTTGGTGAGGGTTGGGATTCACGGGACGAGATCCAGCCTAAGTATATTGTCACTTCTG<br>ATTCGTTCGATCACTTACTCCCCTTACTTATCCTGCGGCTACTGTTTCCGCTGGCTCGTAAG |

| | |
|---|---|
| | CTCTACGTTCGGCAATCTACCCGCGAGGTCAGACGTGACACTCTTAAACTAAAAATTGGTA<br>GCTTCTTTGGCTGAATTGCTGGATCTTATTCGTTCACCCAATAAAACGGTACAGCTTCAAGC<br>AATATCCTCAGTAAGTTAATACCCGTTGTACTATTACTTTCACGACCGTTCGACGTTCCCGCT<br>CTATTTATTAAGAGCTGTCACTTCGAGTCTTTAGCTCACTTAGGAATTAGCTGAGTTTAGGCT<br>CAGCCCTCTTGGGTTGCTTGTACTTTCAGAGTTATTCGCACGGCTGGTTTTGTCGAGTGGG<br>GAATTGTGGTTGACCGAAAGTCCGCTATCCTTCAACGCCGAATCAGCTCTTGCCCTTTACT<br>ATCTTCAATCTCTTGGAGGCTATTACGGGCGGGGGCAAGAGATTAGAACTGCAAGACACC<br>CGTTGATAATCGAGTCGCTCGATAGATTGTCGAGAGCCGGAGAGATTAGTACGTTATTCAA<br>GGCAATACGTGCAGGGTTAATCTGGGCGCGTTGTAGTCTACGCTGGCGTAAGTCCCCAATA<br>ACACGCTCGTCCGGCGAGTCACGATCCTCTAGGCGGTGTTCAACGCGTACGCCAGCTATT<br>TGGGATACTTAGCTACGTTACACGTAAGAATATCTTAGCGGAGGATCGCCCTGCTTCCGCTT<br>GGACGGATAAACGGGAGAGTGGGCGCGTATAGCGCAGGCGGTGTGAAGGCTTTTAAGTA<br>ATTCTAGCCCTCTTTGAACGGTATTTCCCAATTTGGAGATTACCGGATAGCGCGTTTAAATG<br>AGTGTCAGAGAAACGGAAGCCGAAGTCTTTCCATTCCGGATGTTTGGAAATGCTCTGTTTA<br>TAGAAGTCGATGAACTTACGGCAGTCCTCTATATTAAATTCGAATTTTTCATACCCTTTCTGC<br>GGGCTACCGTTTTTAGTGTGCGTGCTATGATTGCGGATGCGCAGAATATCCTCAGACGGGT<br>TGTAGAATTTGATGCTTTTCGCGGAAAAGAACACTTTCGGTAACATTTTATTCGCACCCGGC<br>AGGAGCTTGTACACGATTTTCTTATAGCCTTCACCCTTGTTTTCTTTGATCGCTTTATCGTCG<br>AAGATCTTGTTGTTCTTCTTGTTCATTACGCCCAGATAGTATTTGTCGTCTTTGATGAACAGG<br>ATTGCGGTGTTGTCCGGCTCTTTGTTCTTATCCCAGCCGTTCGCCAGCGTGCTGTTTTCGA<br>AGTTCAGTTTGAATTTCTCGTCAGAGTAAGGCTTCTGCGTGATGTAGTTGCGGATTTTATTG<br>TAGAGAGGGACGATGTTTGCCAGTTCGAAGTAACATTCTTCGAACACCAGATAGAAGTGTT<br>CATCTTTATCCAGAATGTTCGCCTTGTCCTCGCTCTGGCTGATGTGGAAGATTTTGAGCTTG<br>TGTAATAAGTTATTCGTCTGATCTAATAAGTCTTTAATTGCTTTCACATCGTCCTCCGCAGAT<br>GCTTGAAGCAGATCTTTCTTACCCTGATTCTGGTACTTGATAGAGATCTGCGCCAGATTGTC<br>TTTGTTTTGAGCAATTTCGTCGAAGATCATCGGGATTGCCGCAAAGTTCGCCAGAATTTCCT<br>CAAAACGACACTGTTTATCAATATCACGATGTTTATTAAATTCCTCAAGTGCCAGTTTGATAG<br>TTTCTAAGCTCAGGTATTTAGCTTTTTCTGTTTTCTTTGCAATCAGTTCCTGTTCCTTCTTGG<br>ACGGGTTGTCCAGATTTTTCGGCGCGATTTGTTGGGTGATGTATTCCAAAACTGCCGTGCC<br>GATCACGCTATAGTCATCGAAAACTTGTTGACTGAGATCGGTCAGAGATTTGTCGTTTTTAA<br>AGTAAATCTTAGACAGATCTAGTTTCTGCGCTTTGAGGTCGTCAAAGAGCAGGGACAGAGT<br>TTCTTTAATAGATTTCTCTTCCACGGTTTTGAACGCCGCAATCTGCTCATAAAAGCTCTGCAT<br>CGTGGTGACAACGTCGCTATCATCTTCCAGTTTATCAATTACGAAGGATTTAGATTCGGTGT<br>CCGATAAAATCTGTTTAAACAGAACGGACATTTTATACTTTTTCAGGGTTTTGTCGTTGATTT<br>GTTGGCTATACAGGTTAATGTATTCGTTGATGCCCTTACGCTTGGTGTTTTCGCCGTTAACA<br>AATTTGCCACCAATAATGGTGTTGAATTTGGTGATGCCAGATTGATTCAGGTAATTGTTGAA<br>ATTAGCGATTTCGAAAACCTCGTCCAGTGAGAAAACACGCTGGTTAACTTCGGAGGTTTTA<br>TAGTCGATGTCGAAGGTCAGTTCTTCCGCCAGATCTTTCTTGATCTGTTCATAGTTAATAGC<br>TTCCGGTGCTTTGTCTTTCAGAGATTCATATTTCGCTTTGTTTTCCAGAAACTTCGGCAGGT<br>TGTCGTCCACGATACGATAAATAATAGAGGTCGGAATATCGTTGCTCGAATATACATTCTTAC<br>GGTTTTCATGAAAACCTTTGAAATACGTCGTCCAGCCTTTGAAAGACTTGATGATTTCGGTA<br>TTCGAATATGACCTGATCAAAGATAAACGTTTCACCGAAGATAAGTTCTTTTTCCACTGTCC<br>GATTACCATCAACTTCAAATCTAGCGGTGCGAACAAGTTCAACGATGAAATTAACTTATTACT<br>GAAAGAGAAAGCTAATGACGTACACATCTTATCTATTGATCGCGGTGAACGTCATTTAGCAT<br>ACTATACACTGGTAGATGGTAAAGGTAATATTATTAAACAGGATACTTTCAATATTATCGGTAAT<br>GACCGTATGAAAACCAACTATCACGATAAGCTGGCGGCGATCGAAAAAGATCGTGATTCTG<br>CGCGTAAAGATTGGAAGAAAATTAACAATATCAAAGAAATGAAAGAAGGCTATCTGAGCCAA<br>GTGGTGCACGAGATCGCAAAACTGGTGATTGAATATAACGCTATCGTGGTTTTCGAAGATC |

| | |
|---|---|
| | TGAACTTTGGTTTTAAACGTGGTCGCTTCAAAGTAGAAAAACAGGTGTACCAAAAACTGGA<br>AAAAATGCTGATTGAAAAACTGAACTATCTGGTTTTTAAAGACAACGAATTTGACAAAACGG<br>GTGGCGTACTCCGTGCCTATCAGCTGACCGCTCCGTTCGAAACGTTCAAGAAAATGGGTA<br>AACAAACGGGGATTATCTATTATGTGCCAGCTGGTTTCACCTCCAAGATTTGTCCAGTTACG<br>GGCTTCGTTAACCAGCTGTACCCGAAATACGAGAGCGTTAGCAAATCTCAAGAATTTTTCA<br>GCAAATTCGACAAGATCTGCTATAATCTGGATAAAGGCTATTTCGAGTTCAGCTTCGATTACA<br>AAAACTTCGGCGATAAAGCGGCTAAAGGTAAGTGGACTATTGCTAGCTTTGGTAGCCGTCT<br>GATTAACTTTCGCAACTCCGACAAAAACCATAATTGGGACACGCGTGAAGTGTATCCGACC<br>AAAGAACTGGAAAAATTACTGAAAGACTATTCCGGACACTCAGAAGGGTTATAGGAATAGTC<br>ACTACTGGGGTAAGCACTTCGGAAATTATATTATTCTCGCTTCTTATTGCGGTAACGTGATCC<br>TGAACGATACTTATTACTTTGTAATTTACTTAACGTCGGAGTCCCTGCAATCTTCTAGTACCC<br>GCTTCCCGAATACAGGAGATAACTTTTTAACACTCAAGAGTTGCTTCGTGCTTAGCCAGTCT<br>TGGATTTGATTGCTCTAATCCTTCAACGTGTCAAAGACAGTGTATCTGGTCAAGTAAAGTCT<br>AGAGAAAGGCGTAGTCAGTTACGGAGTTATCCCACCTTAGTGTTACTCCGATTTAATTTCTG<br>CTTTCTTTGATTTCTACCCGACTTTCGCCGTGACTTCAATAGAGAGGCAGGCTCTTGCTATT<br>TCTTTCAAGGGCTTGTCCAACTACCTAATTAAGATAAAGATACGGCAGTTGACGCACTGCC<br>GATAATTTCTTTACGTCAGCGAAATTAAATCGAGCACCAGTCGTAGAGTCGCGGTTGCCTA<br>GCAGTTTATCTCGCGTACGGGCCTTCGCTACTTACACGATACCTAGTACGTGGATTCGGGT<br>AGCACCAGAAGTCTATAGCATGTGCATACCTTTGGTCGAAAAAAAAAGCCCGCACTGTCAG<br>GTGCGGGCTTTTTTCAGTGTTTCCTTGCCGGATTACGCCCCGCCCTGCCACTCATCGCAG<br>TATTGTTGTAATTCATTAAGCATTCTGCCGACATGGAAGCCATCACAAACGGCATGATGAAC<br>TTGGATCGCCAGTGGCATTAACACCTTGTCGCCTTGCGTATAATATTTTCCCATAGTGAAAA<br>CGGGGGCGAAGAAGTTGTCCATATTTGCTACGTTTAAATCAAAACTGGTGAAACTCACCCA<br>GGGATTGGCACTGACGAAAAACATATTTTCGATAAAC |
| p1392 | GTTTGCAATAAGGGACAAGTTACGAGTGTAGACACGCAGAATTATCCAGCCTTTAGTCTTTA<br>GGAAGGCAAAGCTATTGTACGCGGTAGCCGTCGTAGCAATTTACCAACTGTAGAATTATTG<br>GACACACGTAACAAGGGCTTACAGTTGAAGTTTAATAAGGTCACACGCAAAACCGCTAAGG<br>AATAATCGCACCGTTAGCGAAAGAATATTTCAGAGCGGTTAGTAAAGGTTGAGTAAAGTGAG<br>ATTCCAAAGTGAGCCTTTATAAAAAGTAAAGAGCTATAATAAAACCGTCGATCGGAAAACAAT<br>CGCCTGAAATCTCAAGCACGTTGCCCTTTCTAACGTCGCTAAGGTTTCGTAAACCCGTTTG<br>ATTAGGAAGAAGAATAAGTAACCCGATTAGGTTTGAGATCGCGGGTTATCGGTTTGGATTAA<br>AAGTGGATACCAGCGGAGTCAACGCCGACGCAAACGTACAGTGATCCAATCCTGTTCCAC<br>GGTCAAGCACAATCAGCTAGCAAGATCTTGGAATAGAGTCGTTGCACCGCTTTGATTTACAT<br>GCTCTCCATTGCACAACATTCCGGAAGGACTGGCTTCTCTGCCATGATCGGATAATGAAAA<br>ACATCAGTATGCCCTGTCATTTTTCTTTGGGTGTCCTCAAATAATTGCCCTCACGTTATCGTA<br>TGTGACGCGCTCATCTATGCTCGAAGTATTCCTTGTTCTCCCATCTTTTAATAGAAAGTCTTT<br>AATGAACGTGTCGTTACGCAGTGTATGAACTCTTGTTTTATAGGGCAGACTTTGGCGTGGC<br>CTAAGTGTGTTCGATAAGAAGGCAAGGACAACTAGCTGACGCGCTGTAATACGGATATTATG<br>GCACGGTTGATACAAACGCTGATATCCTGATTTGCTAATGTGCCCAACACTTTAGTTGAGTG<br>CCACGTTCCGACTACAAGTTGCTTCAAGAGGGGAATTTGGATTTGGCAATAGCCCCCCGTT<br>TCTACCTCAAGAGGCGACGAGTATTAACCGCGCCAGCTTTCGGCACAAGGGCCAAAGAAG<br>ATTCCAATTTCTTATTCCCGAATAACCTCCGAATCCCTGCGGGAAAATCACCGACCGAATAG<br>CCTAGAAGCAAGGGGGAACAGATAGGTATAATTAGCTTAAGAGAGTACCAGCCGTGACAAC<br>ACCGTAGTAACCACAAACTTACGCTGGGGCTTCTTTGGCGGATTTTTACAGATACTAACAAG<br>GTGATTTGAAGTACCTTAGTTGAGGATTTAAACGCGCTATCCGGTAGTCTACAAATTGGGAA<br>ATACCGTTCAAAGAGGGCTAGAATTACTTAAAAGCCTTCACACCGCCTGCGCTATACGCGC<br>CCACTCTCCCGTTTATCCGTCCAAGCGGAAGCAGGGCGAACTTCCGCTAAGATATTCTTAC<br>GTGTAACGTAGCTAAGTATCCCAAATAGCTGGCGTACGCGTTGAACACCGCCTAGAGGATC |

| | |
|---|---|
| | GGGAGTCGCCGGACGAGCGTGTTATTGGGGACTTACGCCAGCGTAGACTACAACGCGCC<br>CAGATTAACCCTGCACGTATTGCCTTGAATAACGTACTAATCTCTCCGGCTCTCGACAATCT<br>ATCGAGCGACTCGATTATCAACGGGTGTCTTGCAGTTCTAATCTCTTGCCCCCGCCCGTAA<br>TAGCCTCCAAGTGATTCAAGATAGTAAAGGGCAAGAGCTTATTCGGCGTTGAAGGATAGCG<br>GACTTTCGGTCAACCACAATTCCCCACTCGACAAAACCAGCCGTGCGAAGAACTCTGAAA<br>GTACAAGCAACCCAAGAGGGCTGAGCCTAAACTCAGCTAATTCCTAAGTGAGCTAAAGACT<br>CGAAGTGACAGCTATTAATAAATAGAGCGGGAACGTCGAACGGTCGTGAAAGTAATAGTAC<br>AACGGGTATTAACTTACTGAGGATATTGCTTGAAGCTGTACCGTTTTATTGGGTGAACGAAT<br>AAGATCCAGCAATTCAGCCAAAGAAGCTACCAATTTTTAGTTTAAGAGTGTCACGTCTGACC<br>TCGCGGGTGGATAGCCGAACGTAGAGCTTACGAGCCAGCGGAAACAGTAGCCGCAGGAT<br>AAGTAAGGGGAGTAAGTGATCGAACGAATCAGAAGTGACAATATACTTAGGCTGGATCTCG<br>TCCCGTGAATCCCAACCCTCACCAACTACGAGATAAGAGGTAAGCCAGAAATCGGCATGGT<br>GGCGACCAACGACTGTTCCCCCCCTGTAACTAATCGTTCCGTCAAAACCTGACTTACTTCA<br>AGGCCAATTCCAAGCGCAAACAATACCGTCCTAGTTCTTCGGTTAAGTTTCCGAAGTAGGA<br>GTGAGCCTACCTCCGTTTGCGTCTTGTTACCACTGACCCAGCTATTTACTTTGTATTGCCTG<br>CAATCGAATTTCTGAACTCTCAGATAGTGGGGATAACGGGAAAGTTCCTATATTTGCGAACT<br>AACTTAGCCGTCCACCTCGAAGCTACCTACTCACACCCACCCCGCGCGGGGTAAATAAGG<br>CACTAATCCCAGCTTAGAGCTTGCGTAGCACTTAGCCACAAGTTAATTAACAGTTGTCTGGT<br>AGTTTGGCGGTATTAGCGAGATCCTAGAAGCAAGGCAGAGTTAGTTCTAACCTAAAGCCAC<br>AAATAAGACAGGTTGCCAAAGCCCGCCGGAAATTAAATCTTGCTCAGTTCGGTAACGGAGT<br>TTCCCTCCCGCGTACTTAATTCCCAATAAGAAACGCGCCCAAGTCCTATCAGGCAAAATTCA<br>GCCCCTTCCCGTGTTAGAACGAGGGTAAAAATACAAGCCGATTGAACAAGGGTTGGGGGC<br>TTCAAATCGTCGTTTACCCCACTTTACAACGGAGGGTAAGTAGTTCACCCTATAGTACGAAG<br>CAGAACTATTTCGAGGGGCGTGCAATAATCGAATCTTCTGCGGTTGACTTAACACGCTAGG<br>GACGTGCCCTCGATTCAGTCGCAGGTACTCCTACTCAGACTGCCTCACACCCAGCTAGTC<br>ACTGAGCGATAAAATTGACCCGCCCTCTAAGGTAGCGAGTACGTCCCAAAGGGCTCCGGA<br>CAGGGCTATATAGGAGAGTTTGATCTCGCCCCGACAACTGCAACCCTCAACTCCCTTAGAT<br>AATATTGTTAGCCGAAGTTGCACGACCCGCCGTCCACGGACTGCTCTTAGGGTGTGGCTC<br>CTTAATCTGACAACGTGCAACCCCTATCGAGGGCGATTGTTTCTGCGAAAGGTGTTGTCCT<br>AATAGTCGCGACATTTGGCCCTTGTAGGTGTGAAACCACTTAGCTTCGCGCCGTAGTCCTA<br>AAGGCCCACCTATTGACTTTGTTTCGGGTAGCACTAGGAATCTTAACAATTTGAATTTGGAC<br>GTGGAACGCGTACACCTTGATCTTCGAATAATTCTAGGGATTTGGAAGTCCTCTACGTTGAC<br>ACACCTACAATGCTCCAAGTAAATATACGAATAACGCGGGCCTCGCGGAGCCGTTCCGAAT<br>CGTCACGTGTTCGTTTACTGTTAATTGGTGGCAAATAAGCAATATCGTAGTCCGTCAGGCCC<br>AGCCCTGTTATCCACGGCGTTATTTGTCAAATTGCGTAGAACTGGATTGACTGCCTGACAAT<br>ACCTAATTATCGGTACGAAGTCCCCGAATCTGTCCGGCTATTTCACTAATACTTTCCAAACG<br>CCCCGTATCCAAGAAGAACGAATTTATCCACGCTCCCGTCTTTGGGACGAATACCGCTACA<br>AGTGGACAGAGGATCGGTACGGGCCTCTAATAAATCCAACACTCTACGCCCTCTTCAAGAG<br>CTAGAAGAACAGGGTGCAGTTGGAAAGGGAATTATTTCGTAAGGCGAGCCAATACCGTAAT<br>TAATTCGGAAGAGTTAACACGATTGGAAGTAGGAATAGTTTCTAACCACGGTTACTAATCCT<br>AATAACGGAACGCTGTCTGATAGATTAGTGTCAGCGCTCACTACCAAAGAAAAATAAAAAGA<br>CGCTGAAAAGCGTCTTTTTATTTTTCGGTCCAGTGTAACTCAGGCAAAAGCACGTAATATTC<br>GTACTCACCAAACGAAACTCATCCGGCGCATCGCGCTTCTTCCTCCGTAAGCGTCACCCC<br>CATTACTTAAAGAGTGCATGTGCATATTTTGTTATCAATAAAAAAGGCCGCGATTTGCGGCCT<br>TATTGTTCGTCTTGCCGGATTAGATAGCTACCGGTGCTTTAATACCCGGATGCGGATCATAG<br>CCTTCGATTTCGAAGTCCTCAAAACGATAATCGAAGATGCTTTCCGGTTTGCGTTTGATAAT<br>CAGTTTCGGGAGCGGGCGTGGCTCACGGCTTAATTGTAAATGCGTCTGATCCATGTGATTT<br>GAGTACAGGTGAGTATCCCCACCAGTCCAAACAAAGTCACCAACTTCCAGATCACACTGCT |

| | |
|---|---|
| | GTGCCATCATATGAACTAATAAGGCGTAGGAGGCAATGTTAAACGGTAAGCCCAGAAACAC<br>GTCGCAAGAACGCTGGTACAGTTGGCACGATAACTTACCATCCGCAACATAGAATTGAAAG<br>AAGGCATGACACGGTGCTAAAGCCATTTTGTCTAATTCCCCCACGTTCCATGCGGACACGA<br>TAATCCGGCGAGAGTCCGGATCATTTTTCAGTTGGTTAAGAACGGTAGTGATCTGATCAATA<br>TGCCGACCATCCGGCGTAGGCCATGCACGCCATTGCTTACCATACACTGGCCCTAAGTCA<br>CCGTTTTCATCTGCCCACTCATCCCAGATGGTAACGTTATTCTCGTGCAGGTACGCAATGTT<br>CGTATCGCCTTGCAGAAACCATAATAACTCGTGAATAATAGAACGGAGGTGGCAACGCTTG<br>GTAGTGACCAGCGGGAAACCGTCTTGCAGGTTGAAACGCATCTGATGACCAAAGATAGAC<br>AGCGTACCAGTGCCAGTACGATCATTCTTCTGAGTGCCTTCGTCCAGCACTTTTTGCATCA<br>GTTCCAGATACTGTTTCATTTTAGCTTCCTTAGCTTGCGAAATCTCGATAACTCAAAAAATAG<br>TAGTGATCTTATTTCATTATGGTGAAAGTTGTCTTACGTGCAACATTTTCGCAAAAAGTTGGC<br>GCTTTATCAACACTGTCCGAATGACAAATGGTTACAATTATTGAACACCCTTCGGGGTGTTT<br>TTTTGTTTCTGGTTTCCCGAGGCCGAACTTTTGTTGCAATGGCTGTCTACCCTGTCTACCT<br>GAGTAAAGAAAAATACATTTAATTCAGTATATTAACTTGGGTAGACAGCCTTTTTTTACTGTCT<br>ACCTTCTGTCTACCCTCTCTACCTGATTTTACCTGAATCAGACAGGGAGGTAGACACGGGG<br>TAGACAGTGGATAAAAGCACTCTACCCCACTGAAAGCAGTGCCATTACTGGCATGGTTGCC<br>AGTAAGGTTGATAAGGTAGACAAGGGGAGGGACAACTCAAAACTTTTTAAACGAGGGGGT<br>AAAACGCAGATCAAAACGATCTCAAGAAGATCATCTTATTAATCAGATAAAATATTTCTAGATT<br>TCAGTGCAATTTATCTCTTCAAATGTAGCACCGGCGCGCCGTGACCAATTATTGAAGGCCG<br>CTAACGCGGCCTTTTTTTGTTTCTGGTTTCCCGAATAGAGCGACTTCTCCCCAAAAAGCCT<br>CGCTTTCAGCACCTGTCGTTTCCTTTCTTTTCAGAGGGTATTTTAAATAAAAACATTAAGTTA<br>TGACGAAGAAGAACGGAAACGCCTTAAACCGGAAAATTTTCATAAATAGCGAAAACCCGCG<br>AGGTCGCCGCCCCGTAACCTGTCGGATCACCGGAAAGAACCTGTAAAGTGATAATGATTAT<br>CATCTACATATCACAACGTGCGTAAAGGGTAAGTATGAAGGTCGTGTACTCCATCGCTACCA<br>AATTCCAGAAAACAGACGCTTTCGAGCGTCTTTTTTCGTTTTGGTCACGACGTACGGTGGA<br>AGATTCGTTACCAATTGACAGCTAGCTCAGTCCTAGGTATATACATACATGCTTGTTTGTTTG<br>TAAACTACTGTTTTCATTAAAGAGGAGAAAGGAAGCCATGTCCATCTATCAGGAGTTTGTTA<br>ACAAGTATTCCCTGTCTAAAACCCTGCGTTTTGAACTGATCCCGCAGGGCAAAACTTTGGA<br>AAACATTAAAGCGCGTGGCCTGATTCTGGATGACGAAAAACGTGCAAAGGATTACAAGAAA<br>GCTAAACAGATCATCGACAAATATCACCAGTTCTTTATCGAAGAAATTCTGTCCTCGGTGTG<br>CATCAGTGAGGATCTGTTACAGAATTATTCTGATGTATACTTTAAACTTAAAAAGTCCGATGA<br>CGATAATCTGCAAAAAGATTTCAAGTCAGCCAAAGATACCATCAAGAAACAGATCTCAGAAT<br>ATATTAAAGATAGCGAAAAGTTCAAAAACCTGTTTAACCAAAACCTCATTGATGCTAAGAAAG<br>GCCAAGAATCTGACCTGATCTTATGGCTGAAACAGAGCAAAGATAACGGCATTGAACTGTT<br>CAAAGCTAATAGCGACATCACCGATATTGATGAAGCGCTCGAAATCATCAAGTCTTTCAAAG<br>GCTGGACGACGTATTTCAAAGGTTTTCATGAAAACCGTAAGAATGTATATTCGAGCAACGAT<br>ATTCCGACCTCTATTATTTATCGTATCGTGGACGACAACCTGCCGAAGTTTCTGGAAAACAA<br>AGCGAAATATGAATCTCTGAAAGACAAAGCACCGGAAGCTATTAACTATGAACAGATCAAGA<br>AAGATCTGGCGGAAGAACTGACCTTCGACATCGACTATAAAACCTCCGAAGTTAACCAGCG<br>TGTTTTCTCACTGGACGAGGTTTTCGAAATCGCTAATTTCAACAATTACCTGAATCAATCTG<br>GCATCACCAAATTCAACACCATTATTGGTGGCAAATTTGTTAACGGCGAAAACACCAAGCG<br>TAAGGGCATCAACGAATACATTAACCTCTATAGCCAACAAATCAACGACAAAACCCTGAAAA<br>AGTATAAAATGTCCGTTCTGTTTAAACAGATTTTATCGGACACCGAATCTAAATCCTTCGTAA<br>TTGATAAACTGGAAGATGATAGCGACGTTGTCACCACGATGCAGAGCTTTTATGAGCAGATT<br>GCGGCGTTCAAAACCGTCGAAGAGAAATCTATTAAAGAAACTCTGTCCCTGCTCTTTGACG<br>ACCTCAAAGCGCAGAAACTAGATCTGTCTAAGATTTACTTTAAAAACGACAAATCTCTGACC<br>GATCTCAGTCAACAAGTTTTCGATGACTATAGCGTGATCGGCACGGCAGTTTTGGAATACAT<br>CACCCAACAAATCGCGCCGAAAAATCTGGACAACCCGTCCAAGAAGGAACAGGAACTGAT |

| | |
|---|---|
| | TGCAAAGAAAACAGAAAAAGCTAAATACCTGAGCTTAGAAACTATCAAACTGGCACTTGAG |
| | GAATTTAATAAACATCGTGATATTGATAAACAGTGTCGTTTTGAGGAAATTCTGGCGAACTTT |
| | GCGGCAATCCCGATGATCTTCGACGAAATTGCTCAAAACAAAGACAATCTGGCGCAGATCT |
| | CTATCAAGTACCAGAATCAGGGTAAGAAAGATCTGCTTCAAGCATCTGCGGAGGACGATGT |
| | CAAAGCAATTAAAGACTTATTAGATCAGACGAATAACTTATTACACAAGCTCAAAATCTTCCA |
| | CATCAGCCAGAGCGAGGACAAGGCGAACATTCTGGATAAAGATGAACACTTCTATCTGGTG |
| | TTCGAAGAATGTTACTTCGAACTGGCAAACATCGTACCTCTCTACAATAAAATCCGCAACTA |
| | CATCACGCAGAAGCCTTACAGTGACGAGAAATTCAAACTGAACTTCGAAAACAGCACGCT |
| | GGCGAACGGCTGGGATAAGAACAAAGAGCCGGACAACACCGCAATCCTGTTCATCAAAGA |
| | CGACAAATACTATCTGGGCGTAATGAACAAGAAGAACAACAAGATCTTCGACGATAAAGCG |
| | ATCAAAGAAAACAAGGGTGAAGGCTATAAGAAAATCGTGTACAAGCTCCTGCCGGGTGCG |
| | AACAAAATGTTACCGAAAGTGTTCTTTTCCGCGAAAAGCATCAAATTCTACAACCCGTCTGA |
| | GGATATTCTGCGCATCCGCAATCATAGCACGCACACTAAAAACGGTAGCCCGCAGAAAGG |
| | GTATGAAAAATTCGAATTTAATATAGAGGACTGCCGTAAATTCATCGACTTCTATAAACAGAG |
| | CATTTCCAAACATCCGGAATGGAAAGACTTCGGCTTCCGTTTCTCTGACACTCAGCGCTAT |
| | AATAGCATCGACGAGTTCTACCGCGAAGTGGAGAATCAGGGCTATAAACTGACCTTCGAGA |
| | ACATTAGTGAGTCGTACATCGACTCCGTTGTGAATCAGGGTAAACTGTACCTGTTTCAGATC |
| | TATAATAAAGACTTTAGCGCGTACAGCAAAGGCCGCCCGAATCTGCACACCCTTTACTGGA |
| | AAGCATTATTTGACGAACGTAACCTGCAAGATGTGGTGTATAAACTGAACGGTGAGGCGGA |
| | ACTTTTCTACCGTAAACAGAGTATCCCGAAGAAAATCACGCATCCGGCAAAAGAAGCTATTG |
| | CCAACAAAAACAAAGACAACCCGAAGAAAGAAAGTGTATTCGAATATGACCTGATCAAAGA |
| | TAAACGTTTCACCGAAGATAAGTTCTTTTTCCACTGTCCGATTACCATCAACTTCAAATCTAG |
| | CGGTGCGAACAAGTTCAACGATGAAATTAACTTATTACTGAAAGAGAAAGCTAATGACGTAC |
| | ACATCTTATCTATTGATCGCGGTGAACGTCATTTAGCATACTATACACTGGTAGACGGTAAAG |
| | GTAATATTATTAAACAGGATACTTTCAATATTATCGGTAATGACCGTATGAAAACCAACTATCA |
| | CGATAAGCTGGCGGCGATCGAAAAAGATCGTGATTCTGCGCGTAAAGATTGGAAGAAAATT |
| | AACAATATCAAAGAAATGAAAGAAGGCTATCTGAGCCAAGTGGTGCACGAGATCGCAAAAC |
| | TGGTGATTGAATATAACGCTATCGTGGTTTTCGAAGATCTGAACTTTGGTTTTAAACGTGGT |
| | CGCTTCAAAGTAGAAAAACAGGTGTACCAAAAACTGGAAAAAATGCTGATTGAAAAACTGA |
| | ACTATCTGGTTTTTAAAGACAACGAATTTGACAAAACGGGTGGCGTACTCCGTGCCTATCA |
| | GCTTACCGCTCCGTTCGAAACGTTTAAGAAAATGGGTAAACAAACGGGGATTATCTATTATG |
| | TGCCAGCCGGTTTCACCTCCAAGATTTGTCCAGTTACGGGCTTCGTTAACCAGCTTTACCC |
| | GAAATACGAGAGCGTTAGCAAATCTCAAGAATTTTTCAGCAAATTCGACAAGATCTGCTATA |
| | ATCTGGATAAAGGCTATTTCGAGTTCAGCTTTGATTACAAAAACTTCGGCGATAAAGCGGCT |
| | AAAGGTAAGTGGACTATTGCTAGCTTTGGTAGCCGTCTGATTAACTTTCGCAACTCCGACA |
| | AAAACCATAATTGGGACACGCGTGAAGTGTATCCGACCAAAGAACTGGAAAAATTACTGAA |
| | AGACTATTCCATCGAATATGGTCATGGGGAGTGCATTAAAGCGGCGATTTGCGGTGAATCC |
| | GATAAGAAATTTTTCGCCAAACTGACCAGCGTGCTTAACACCATTCTCCAAATGCGTAATTC |
| | TAAAACGGGTACGGAGCTTGACTACCTGATTTCTCCGGTAGCCGACGTTAACGGCAACTTC |
| | TTCGATTCTCGTCAAGCACCGAAAAATATGCCACAAGACGCGGATGCCAACGGTGCATACC |
| | ATATCGGCCTTAAAGGCTTAATGTTATTAGGCCGTATCAAGAATAATCAGGAGGGCAAGAAA |
| | TTAAATCTGGTTATCAAAAACGAAGAATACTTCGAGTTCGTTCAGAATCGTAACAATTAATGT |
| | ATGCTTAAGCAGATCGGTAATAAAGACGAACAATAAGACGCTGAAAAGCGTCTTTTTTCGTT |
| | TTGGTCCTGTTCCGGCGCGATAGTGTGAACATGCTATAGACTTCTGGTGCTACCCGACTGA |
| | CAATTAATCATCCGGCTCGTATAATGCTAGCAATTTCTACTGTTGTAGATCATTCCGGAACGT |
| | TCCAGCGCTGCAATTTCTACTGTTGTAGATCTGATTTTTCACATGTTACCTTTCAATTTCTAC |
| | TGTTGTAGATCCGAAAACGTAAAGCTTCAGCTGTAATTTCTACTGTTGTAGATATCATATCTG |
| | GCGTTAATGGAGTTTCGTGACGAACAATAAGTCCTCCCTAACGGGGGGCAATTTTTATTGAT |

| | |
|---|---|
| | AACAAAAGTAACTTCGAGCTTGTCTACCTCCTAGCTCGTAAATTGCACGCTGATAGTCTCCC<br>AATTGCGAAGGACCAAAACGAAAAAACACCCTTTCGGGTGTCTTTTCTGGAATTTGGTACG<br>CAGTACTAGGTATCGTGTAAGTAGCGAAGGCCCGTACGCGAGATAAACTGCTAGGCAACC<br>GCGACTCTACGACTGGTGCTCGATTTAATTTCGCTGACGTAAAGAAATTATCGGCAGTGCG<br>TCAACTGCCGTATCTTTATCTTAATTAGGTAGTTGGACAAGCCCTTGAAAGAAATAGCAAGA<br>GCCTGCCTCTCTATTGAAGTCACGGCGAAAGTCGGGTAGAAATCAAAGAAAGCAGAAATTA<br>AATCGGAGTAATACTAAGTTGGGATAACTCCGTAACTGACTACGCCTTTCTCTAGACTTTACT<br>TGACCAGATACACTGTCTTTGACACGTTGAAGGATTAGAGCAATCAAATCCAAGACTGGCT<br>AAGCACGAAGCAACTCTTGAGTGTTAAAAAGTTACTTCCTGTATTCGGGACGAGGGTACTA<br>GAAGATTGCAGGGACTCCGACGTTAAGTAAATTACAAAGTAATAAGTATCGTTCAGGATCAC<br>GTTACCGCAATAAGAAGCGAGAATAATATAATTTCCGAAGTGCTTACCCCAGTAGTGACTATT<br>CCTATAACCCTTCTGAGTGTCCGGAGGCGGAAATTTGCCACGAAAGAGAAAGTATTTCCCC<br>GACAATAATAAAGGGGCGCTCCTCAGCTTTTCCACTTGGTTGGGTAAGCTAGGCAACTCTG<br>AAAGGAGTTTCGGCGAAGTGAAGCCGACACCTTTGAATTGTTTTAGGGGCGTTATTCGAG<br>GGCAATCGGAGCTAACTTCAAGACTACTTCTTTGTTGAATACTAAATAGTGCAAAGGTCGTG<br>TTTCCTCAAGGATACTCCGCTAACAATATAGGATTCCAATCAGATTCAGCACTGGCGGTACG<br>GGTGTTGCGGTGAGGCGTTCGGGTTTACGGCTCGAAGCTAGCACGGTAGGAAGCCTGAC<br>AATCACCAAGCAAAAGGGCCGTCGAAGGCCCACAAGATACGAAAGCTCTCGAAGCCTTAT<br>CCTTGACCGATCCACCTATTTAGGCAGTTACGCACAAAAGCTACCCAATAATCCGTGACAG<br>GCACAATATCACGGAACAAAACCGAAAACTCTCGTACACGGTTAGGTTTTCGCTAGGAAGA<br>ATAAACCTCTATCTTGATTATAAGAAGGCTCCCCAAGCACCCCCAAAACCGAAATAGCG |
| p1399 | TCAGATCCTTCCGTATTTAGCCAGTATGTTCTCTAGTGTGGTTCGTTGTTTTTGCGTGAGCC<br>ATGAGAACGAACCATTGAGATCATACTTACTTTGCATGTCACTCAAAAATTTTGCCTCAAAAC<br>TGGTGAGCTGAATTTTTGCAGTTAAAGCATCGTGTAGTGTTTTTCTTAGTCCGTTACGTAGG<br>TAGGAATCTGATGTAATGGTTGTTGGTATTTTGTCACCATTCATTTTTATCTGGTTGTTCTCAA<br>GTTCGGTTACGAGATCCATTTGTCTATCTAGTTCAACTTGGAAAATCAACGTATCAGTCGGG<br>CGGCCTCGCTTATCAACCACCAATTTCATATTGCTGTAAGTGTTTAAATCTTTACTTATTGGT<br>TTCAAAACCCATTGGTTAAGCCTTTTAAACTCATGGTAGTTATTTTCAAGCATTAACATGAAC<br>TTAAATTCATCAAGGCTAATCTCTATATTTGCCTTGTGAGTTTTCTTTTGTGTTAGTTCTTTTA<br>ATAACCACTCATAAATCCTCATAGAGTATTTGTTTTCAAAAGACTTAACATGTTCCAGATTATA<br>TTTTATGAATTTTTTTAACTGGAAAAGATAAGGCAATATCTCTTCACTAAAAACTAATTCTAATT<br>TTTCGCTTGAGAACTTGGCATAGTTTGTCCACTGGAAAATCTCAAAGCCTTTAACCAAAGG<br>ATTCCTGATTTCCACAGTTCTCGTCATCAGCTCTCTGGTTGCTTTAGCTAATACACCATAAG<br>CATTTTCCCTACTGATGTTCATCATCTGAGCGTATTGGTTATAAGTGAACGATACCGTCCGTT<br>CTTTCCTTGTAGGGTTTTCAATCGTGGGGTTGAGTAGTGCCACACAGCATAAAATTAGCTTG<br>GTTTCATGCTCCGTTAAGTCATAGCGACTAATCGCTAGTTCATTTGCTTTGAAAACAACTAAT<br>TCAGACATACATCTCAATTGGTCTAGGTGATTTTAATCACTATACCAATTGAGATGGGCTAGT<br>CAATGATAATTACTAGTCCTTTTCCTTTGAGTTGTGGGTATCTGTAAATTCTGCTAGACCTTT<br>GCTGGAAAACTTGTAAATTCTGCTAGACCCTCTGTAAATTCCGCTAGACCTTTGTGTGTTTT<br>TTTTGTTTATATTCAAGTGGTTATAATTTATAGAATAAAGAAAGAATAAAAAAAGATAAAAAGAA<br>TAGATCCCAGCCCTGTGTATAACTCACTACTTTAGTCAGTTCCGCAGTATTACAAAAGGATG<br>TCGCAAACGCTGTTTGCTCCTCTACAAAACAGACCTTAAAACCCTAAAGGCTTAAGTAGCA<br>CCCTCGCAAGCTCGGTTGCGGCCGCAATCGGGCAAATCGCTGAATATTCCTTTTGTCTCC<br>GACCATCAGGCACCTGAGTCGCTGTCTTTTTCGTGACATTCAGTTCGCTGCGCTCACGGC<br>TCTGGCAGTGAATGGGGGTAAATGGCACTACAGGCGCCTTTTATGGATTCATGCAAGGAAA<br>CTACCCATAATACAAGAAAAGCCCGTCACGGGCTTCTCAGGGCGTTTTATGGCGGGTCTG<br>CTATGTGGTGCTATCTGACTTTTTGCTGTTCAGCAGTTCCTGCCCTCTGATTTTCCAGTCTG<br>ACCACTTCGGATTATCCCGTGACAGGTCATTCAGACTGGCTAATGCACCCAGTAAGGCAGC |

| | |
|---|---|
| | GGTATCATCAACGGGGTCTGACGCTCAGTGGAACGAAAACTCACGTTAAGGGATTTTGGT<br>CATGAGATTATCAAAAAGGATCTTCACCTAGATCCTTTTAAATTAAAAATGAAGTTTTAAATCA<br>ATCTAAAGTATATATGAGTAAACTTGGTCTGACAGTTACGTTTCCACAACCAATTAACCAATT<br>CTGATTTAGAAAAACTCATCGAGCATCAAATGAAACTGCAATTTATTCATATCAGGATTATCAA<br>TACCATATTTTTGAAAAAGCCGTTTCTGTAATGAAGGAGAAAACTCACCGAGGCAGTTCCAT<br>AGGATGGCAAGATCCTGGTATCGGTCTGCGATTCCGACTCGTCCAACATCAATACAACCTA<br>TTAATTTCCCCTCGTCAAAAATAAGGTTATCAAGTGAGAAATCACCATGAGTGACGACTGAA<br>TCCGGTGAGAATGGCAAAAGCTTATGCATTTCTTTCCAGACTTGTTCAACAGGCCAGCCAT<br>TACGCTCGTCATCAAAATCACTCGCATCAACCAAACCGTTATTCATTCGTGATTGCGCCTGA<br>GCGAGACGAAATACGCGATCGCTGTTAAAAGGACAATTACAAACAGGAATCGAATGCAACC<br>GGCGCAGGAACACTGCCAGCGCATCAACAATATTTTCACCTGAATCAGGATATTCTTCTAAT<br>ACCTGGAATGCTGTTTTCCCGGGGATCGCAGTGGTGAGTAACCATGCATCATCAGGAGTA<br>CGGATAAAATGCTTGATGGTCGGAAGAGGCATAAATTCCGTCAGCCAGTTTAGTCTGACCA<br>TCTCATCTGTAACATCATTGGCAACGCTACCTTTGCCATGTTTCAGAAACAACTCTGGCGCA<br>TCGGGCTTCCCATACAATCGATAGATTGTCGCACCTGATTGCCCGACATTATCGCGAGCCC<br>ATTTATACCCATATAAATCAGCATCCATGTTGGAATTTAATCGCGGCCTCGAGCAAGACGTTT<br>CCCGTTGAATATGGCTCATAACACCCCTTGTATTACTGTTTATGTAAGCAGACAGTTTTATTG<br>TTCATGATGATATATTTTTATCTTGTGCAATGTAACATCAGAGATTTTGAGACacaacgtggaggcct<br>gCGTTATCCCCTGATTCTGTGGATAACCGTATTACCGCCTTTGAGTGAGCTGATACCGCTCG<br>CCGCAGCCGAACGCCGACTAGTGGATTTTACGGCTAGCTCAGTCCTAGGTACAATGCTAG<br>CgaattcattaaagaggagaaaggtacccATGGCACGTACCCCGAGCCGTAGCAGCATTGGTAGCCTG<br>CGTAGTCCGCATACCCATAAAGCAATTCTGACCAGCACCATTGAgATCCTGAAAGAATGTGG<br>TTATAGCGGTCTGAGCATTGAAAGCGTTGCACGTCGTGCCGGTGCAAGCAAACCGACCAT<br>TTATCGTTGGTGGACCAATAAAGCAGCACTGATTGCCGAAGTGTATGAAAATGAAAGCGAA<br>CAGGTGCGTAAATTTCCGGATCTGGGTAGCTTTAAAGCCGATCTGGATTTTCTGCTGCGTA<br>ATCTGTGGAAAGTTTGGCGTGAAACCATTTGTGGTGAAGCATTTCGTTGTGTTATTGCAGA<br>AGCACAGCTGGACCCTGCAACCCTGACCCAGCTGAAAGATCAGTTTATGGAACGTCGTCG<br>TGAGATGCCGAAAAAACTGGTTGAAAATGCCATTAGCAATGGTGAACTGCCGAAAGATACC<br>AATCGTGAACTGCTGCTGGATATGATTTTTGGTTTTTGTTGGTATCGCCTGCTGACCGAACA<br>GCTGACCGTTGAACAGGATATTGAAGAATTTACCTTCCTGCTaATTAATGGTGTTTGTCCGG<br>GTACACAGCGTTAACTAGGGCCCATACCCCCAATTATTGAAGGCCGCTAACGCGGCCTTTT<br>TTTGTTTCTGGTCTGCCCGACGTACGGTGAAtctgattcgttaccaattgacATGATACGAAACGTACC<br>GTATCGTTAAGGTCGTGATTAAACGATCCCGTTTTCAAAAatgaaactggcaccgaacgtaaaacagca<br>gtcacgcggcataaaacacaaagaaacagaagtcattatttttgcgggtagtgatgcctggtcacacgcaaaacaatggcaggaa<br>catgacgcgcgtatggccggagataatgagcctcctgtgtggcttggggagcagcagttatccgaactggataagctgcaaattgtg<br>ccggaaggcagaaaatccgtgcgcatattcagggccggatatcttgcgccagtaatgataaaggcgattggtcagaagctggcgg<br>cggcaggcgtacaggatgcaaatttttaccctgatggtatgcacggtcagaaggtggagaactggcgcgaatatctggcccgtgag<br>cgccagaatctttctgatggtctggtcattgagcttccggtaaagcaaaaggcgcaactttcgcagatggcggacagtgagcgcgcg<br>cagctgcttgccgatcgctttgatggcgtttgcgtacatcctgaaagtgaaatcgttcacgtatggcgcggcggggtatggtgtccggtc<br>agcacaatggagctgagccgcgaaatggtggcgatctattcagagcacagggccactttcagcaagcgcgtaatcaataacgcc<br>gtggaagcgttaaaagttattgccgaaccaatgggcgagccgtccggcgatttgctgccgttcgccaatggtgcgcttgacctgaaa<br>acgggggaattttccccgcacacgccggagaactggatcaccacgcacaacggcattgagtacacgccaccagcacccgggga<br>gaacatccgcgataacgcgccaaactttcataaatggcttgagcacgcagccggaaaagacccgcgcaagatgatgcgtatatgt<br>gccgcgctgtacatgattatggcgaaccggtacgactggcagatgtttattgaggccaccggagacggcgggagcggtaaaagta<br>cattcacacacatagccagccttctggcagggaaacaaaacacggtaagcgctgaaatgacatcgcttgatgatgctggtgggcgt<br>gcgcaggttgtcgggagtcgtcttatcgtcctggcagaccagccgaaatatacaggcgaaggaacgggcatcaagaaaatcacg<br>ggcggcgaccccgtggaaattaacccgaaatatgaaaagcgttttacggcggtaatcagggcggtggtgctggcaaccaataaca<br>atccgatgatattcaccgaacggccggaggtgtggcacgtcgtcgggtgatattccggttcgataacatcgtaagcgaggcagaa |

| | |
|---|---|
| | aaagacagggagctaccggaaaagatcgcggctgaGatccctgtcattatccgccgcttgctggcgaactttgccgaccctgaaa<br>aggcacgggctttactcattgaacagcgtgacggtgatgaagcactggcaataaagcaacagacggatccggttattgagttttgcc<br>agttcctgaattttctggaggaagcacgcggcctgatgatgggcggcggtggcgattcagtgaagtacacgaccagaaacagccttt<br>accgcgtctatctggcgtttatggcgtacgcaggcaggagcaaaccgctaaacgtaaatgactttggcaaggctatgaagccagcc<br>gcgaaagtttacggacatgaatatattacgcggaaagttaaaggagtaacgcagactaacgcaataacaacagacgattgcgac<br>gcgttttttaTAAtgacgcatcctcacgataatatccgggtagGACGAACAATAAGGCCGCAAATCGCGGCCTTT<br>TTTATTGATAACAAAAGGACAGTTTTCCCTTTGATATGTAACGGTGAACAGTTGTTCTACTTT<br>TGTTTGTTAGTCTTGATGCTTCACTGATAGATACAAGAGCCATAAGAACC |
| p521 | TCAGATCCTTCCGTATTTAGCCAGTATGTTCTCTAGTGTGGTTCGTTGTTTTTGCGTGAGCC<br>ATGAGAACGAACCATTGAGATCATACTTACTTTGCATGTCACTCAAAAATTTTGCCTCAAAAC<br>TGGTGAGCTGAATTTTTGCAGTTAAAGCATCGTGTAGTGTTTTTCTTAGTCCGTTACGTAGG<br>TAGGAATCTGATGTAATGGTTGTTGGTATTTTGTCACCATTCATTTTTATCTGGTTGTTCTCAA<br>GTTCGGTTACGAGATCCATTTGTCTATCTAGTTCAACTTGGAAAATCAACGTATCAGTCGGG<br>CGGCCTCGCTTATCAACCACCAATTTCATATTGCTGTAAGTGTTTAAATCTTTACTTATTGGT<br>TTCAAAACCCATTGGTTAAGCCTTTTAAACTCATGGTAGTTATTTTCAAGCATTAACATGAAC<br>TTAAATTCATCAAGGCTAATCTCTATATTTGCCTTGTGAGTTTTCTTTTGTGTTAGTTCTTTTA<br>ATAACCACTCATAAATCCTCATAGAGTATTTGTTTTCAAAAGACTTAACATGTTCCAGATTATA<br>TTTTATGAATTTTTTTAACTGGAAAAGATAAGGCAATATCTCTTCACTAAAAACTAATTCTAATT<br>TTTCGCTTGAGAACTTGGCATAGTTTGTCCACTGGAAAATCTCAAAGCCTTTAACCAAAGG<br>ATTCCTGATTTCCACAGTTCTCGTCATCAGCTCTCTGGTTGCTTTAGCTAATACACCATAAG<br>CATTTTCCCTACTGATGTTCATCATCTGAGCGTATTGGTTATAAGTGAACGATACCGTCCGTT<br>CTTTCCTTGTAGGGTTTTCAATCGTGGGGTTGAGTAGTGCCACACAGCATAAAATTAGCTTG<br>GTTTCATGCTCCGTTAAGTCATAGCGACTAATCGCTAGTTCATTTGCTTTGAAAACAACTAAT<br>TCAGACATACATCTCAATTGGTCTAGGTGATTTTAATCACTATACCAATTGAGATGGGCTAGT<br>CAATGATAATTACTAGTCCTTTTCCTTTGAGTTGTGGGTATCTGTAAATTCTGCTAGACCTTT<br>GCTGGAAAACTTGTAAATTCTGCTAGACCCTCTGTAAATTCCGCTAGACCTTTGTGTGTTTT<br>TTTTGTTTATATTCAAGTGGTTATAATTTATAGAATAAAGAAAGAATAAAAAAAGATAAAAAGAA<br>TAGATCCCAGCCCTGTGTATAACTCACTACTTTAGTCAGTTCCGCAGTATTACAAAAGGATG<br>TCGCAAACGCTGTTTGCTCCTCTACAAAACAGACCTTAAAACCCTAAAGGCTTAAGTAGCA<br>CCCTCGCAAGCTCGGTTGCGGCCGCAATCGGGCAAATCGCTGAATATTCCTTTTGTCTCC<br>GACCATCAGGCACCTGAGTCGCTGTCTTTTTCGTGACATTCAGTTCGCTGCGCTCACGGC<br>TCTGGCAGTGAATGGGGGTAAATGGCACTACAGGCGCCTTTTATGGATTCATGCAAGGAAA<br>CTACCCATAATACAAGAAAAGCCCGTCACGGGCTTCTCAGGGCGTTTTATGGCGGGTCTG<br>CTATGTGGTGCTATCTGACTTTTTGCTGTTCAGCAGTTCCTGCCCTCTGATTTTCCAGTCTG<br>ACCACTTCGGATTATCCCGTGACAGGTCATTCAGACTGGCTAATGCACCCAGTAAGGCAGC<br>GGTATCATCAACGGGGTCTGACGCTCAGTGGAACGAAAACTCACGTTAAGGGATTTTGGT<br>CATGAGATTATCAAAAAGGATCTTCACCTAGATCCTTTTAAATTAAAAATGAAGTTTTAAATCA<br>ATCTAAAGTATATATGAGTAAACTTGGTCTGACAGTTACGTTTCCACAACCAATTAACCAATT<br>CTGATTTAGAAAAACTCATCGAGCATCAAATGAAACTGCAATTTATTCATATCAGGATTATCAA<br>TACCATATTTTTGAAAAAGCCGTTTCTGTAATGAAGGAGAAAACTCACCGAGGCAGTTCCAT<br>AGGATGGCAAGATCCTGGTATCGGTCTGCGATTCCGACTCGTCCAACATCAATACAACCTA<br>TTAATTTCCCCTCGTCAAAAATAAGGTTATCAAGTGAGAAATCACCATGAGTGACGACTGAA<br>TCCGGTGAGAATGGCAAAAGCTTATGCATTTCTTTCCAGACTTGTTCAACAGGCCAGCCAT<br>TACGCTCGTCATCAAAATCACTCGCATCAACCAAACCGTTATTCATTCGTGATTGCGCCTGA<br>GCGAGACGAAATACGCGATCGCTGTTAAAAGGACAATTACAAACAGGAATCGAATGCAACC<br>GGCGCAGGAACACTGCCAGCGCATCAACAATATTTTCACCTGAATCAGGATATTCTTCTAAT<br>ACCTGGAATGCTGTTTTCCCGGGGATCGCAGTGGTGAGTAACCATGCATCATCAGGAGTA<br>CGGATAAAATGCTTGATGGTCGGAAGAGGCATAAATTCCGTCAGCCAGTTTAGTCTGACCA |

| | |
|---|---|
| | TCTCATCTGTAACATCATTGGCAACGCTACCTTTGCCATGTTTCAGAAACAACTCTGGCGCA<br>TCGGGCTTCCCATACAATCGATAGATTGTCGCACCTGATTGCCCGACATTATCGCGAGCCC<br>ATTTATACCCATATAAATCAGCATCCATGTTGGAATTTAATCGCGGCCTCGAGCAAGACGTTT<br>CCCGTTGAATATGGCTCATAACACCCCTTGTATTACTGTTTATGTAAGCAGACAGTTTTATTG<br>TTCATGATGATATATTTTTATCTTGTGCAATGTAACATCAGAGATTTTGAGACACAACGTGGC<br>TTTCCCTGCAGGATTTCGGAGGCCTGCGTTATCCCCTGATTCTGTGGATAACCGTATTACC<br>GCCTTTGAGTGAGCTGATACCGCTCGCCGCAGCCGAACGCCGACTAGTGGATTTTACGGC<br>TAGCTCAGTCCTAGGTACAATGCTAGCGAATTCATTAAAGAGGAGAAAGGTACCCATGGCA<br>CGTACCCCGAGCCGTAGCAGCATTGGTAGCCTGCGTAGTCCGCATACCCATAAAGCAATTC<br>TGACCAGCACCATTGAAATCCTGAAAGAATGTGGTTATAGCGGTCTGAGCATTGAAAGCGT<br>TGCACGTCGTGCCGGTGCAAGCAAACCGACCATTTATCGTTGGTGGACCAATAAAGCAGC<br>ACTGATTGCCGAAGTGTATGAAAATGAAAGCGAACAGGTGCGTAAATTTCCGGATCTGGGT<br>AGCTTTAAAGCCGATCTGGATTTTCTGCTGCGTAATCTGTGGAAAGTTTGGCGTGAAACCA<br>TTTGTGGTGAAGCATTTCGTTGTGTTATTGCAGAAGCACAGCTGGACCCTGCAACCCTGAC<br>CCAGCTGAAAGATCAGTTTATGGAACGTCGTCGTGAGATGCCGAAAAAACTGGTTGAAAAT<br>GCCATTAGCAATGGTGAACTGCCGAAAGATACCAATCGTGAACTGCTGCTGGATATGATTTT<br>TGGTTTTTGTTGGTATCGCCTGCTGACCGAACAGCTGACCGTTGAACAGGATATTGAAGAA<br>TTTACCTTCCTGCTAATTAATGGTGTTTGTCCGGGTACACAGCGTTAACTAGGGCCCATACC<br>CCCAATTATTGAAGGCCGCTAACGCGGCCTTTTTTTGTTTCTGGTCTGCCCGACGTACGGT<br>GAATCTGATTCGTTACCAATTGACATGATACGAAACGTACCGTATCGTTAAGGTTACTAGATT<br>AAAGAGGAGAAATACTAGATGGCAGTAAAGATTTCAGGAGTCCTGAAAGACGGCACAGGA<br>AAACCGGTACAGAACTGCACCATTCAGCTGAAAGCCAGACGTAACAGCACCACGGTGGTG<br>GTGAACACGGTGGGCTCAGAGAATCCGGATGAAGCCGGGCGTTACAGCATGGATGTGGA<br>GTACGGTCAGTACAGTGTCATCCTGCAGGTTGACGGTTTTCCACCATCGCACGCCGGGAC<br>CATCACCGTGTATGAAGATTCACAACCGGGGACGCTGAATGATTTTCTCTGTGCCATGACG<br>GAGGATGATGCCCGGCCGGAGGTGCTGCGTCGTCTTGAACTGATGGTGGAAGAGGTGGC<br>GCGTAACGCGTCCGTGGTGGCACAGAGTACGGCAGACGCGAAGAAATCAGCCGGCGATG<br>CCAGTGCATCAGCTGCTCAGGTCGCGGCCCTTGTGACTGATGCAACTGACTCAGCACGC<br>GCCGCCAGCACGTCCGCCGGACAGGCTGCATCGTCAGCTCAGGAAGCGTCCTCCGGCG<br>CAGAAGCGGCATCAGCAAAGGCCACTGAAGCGGAAAAAAGTGCCGCAGCCGCAGAGTCC<br>TCAAAAAACGCGGCGGCCACCAGTGCCGGTGCGGCGAAAACGTCAGAAACGAATGCTGC<br>AGCGTCACAACAATCAGCCGCCACGTCTGCCTCCACCGCGGCCACGAAAGCGTCAGAGG<br>CCGCCACTTCAGCACGAGATGCGGTGGCCTCAAAAGAGGCAGCAAAATCATCAGAAACGA<br>ACGCATCATCAAGTGCCGGTCGTGCAGCTTCCTCGGCAACGGCGGCAGAAAATTCTGCCA<br>GGGCGGCAAAAACGTCCGAGACGAATGCCAGGTCATCTGAAACAGCAGCGGAACGGAGC<br>GCCTCTGCCGCGGCAGACGCAAAAACAGCGGCGGCGGGGAGTGCGTCAACGGCATCCA<br>CGAAGGCGACAGAGGCTGCGGGAAGTGCGGTATCAGCATCGCAGAGCAAAAGTGCGGCA<br>GAAGCGGCGGCAATACGTGCAAAAAATTCGGCAAAACGTGCAGAAGATATAGCTTCAGCT<br>GTCGCGCTTGAGGATGCGGACACAACGAGAAAGGGGATAGTGCAGCTCAGCAGTGCAAC<br>CAACAGCACGTCTGAAACGCTTGCTGCAACGCCAAAGGCGGTTAAGGTGGTAATGGATGA<br>GACTAATCGTAAATATACGGCTCAGGACGCGAGCACGGCGCAGAAAGGTCTGGTGAAACT<br>GAGCAGCGCCACCGACAGCACATCTGAGACCCTCGCCGCGACACCGAAAGCGGTTAAGG<br>CGGTGAATGATAATGCGAATGGTCGCGTCCCGTCTGAGCGAAAAGTTAACGGACATTCGC<br>TGGCCGGTGATATCAGTGTCACCTCACAGGATATTTTTGACGGTCAGTGTGTTGAAATTGG<br>TCCGGGTCAGGATCTGGATAATTACCAGACGCCGGGTCTGTATTTTCAGCCCGCAAATGCC<br>AATACCAGTGCTGCTCTGCATTACCCGGAAAATAATGCCGGTTCCCTGATGGTTTTAAGAAG<br>CGCAGGGATAACGCAGGTTTATCGCGTGTACAGCGGTTCGCGAAGTTATTTGCGGAGCAA<br>ATATTCCACGCAGCCATGGACGACGTGGACACCCGATGATGCTTTTCCTGTCGGCGCGCC |

| | |
|---|---|
| | GATTCCGTGGCCATCTGACATCGCCCCGCCCGCTTACGCCTTAATGCAGGGGCAGTCATT<br>TGATAAATCTGCATATCCATTGCTTGCTGTAGCGTATCCCTCTGGTGTTATCCCGGATATGCG<br>TGGTCAGACGATAAAGGGCAAGCCGGACGGACGAGCGGTACTCTCGTATGAACAGGACG<br>GTATTAAATCGCACGCTCATACAGCCAGTATTTCCGATACCGATTTGGGAACGAAATATACCA<br>ACTCTTTTGATTATGGTTCAAAACCAACAACCAGTTTTGACTACGGCAATAAGTCCTCCACT<br>GAGGGGGGATGGCACGTACATAACTTTCGTTATTGTGCTACGTCTGCATACCGGGATACTC<br>CTGGCTCAGGGCTGGGGATGCACTCGTCGAATATTTCGTGGTCAGCCGGGGATCGCATTG<br>AGGGGAGTGGTAATCATGCACATGTTACGTGGATTGGTCCCCATGATCACTGGGTTGGTAT<br>CGGTGAGCATAACCATTATGTGGTTATGGGGTATCACGGACATACAGCGACCGTTCATGCA<br>ACCGGGAATACAGAAAACACCGTTAAAAATATTGCGTTTAACTACATTGTGAGGCTTGCATA<br>ATGGCTTTTGAAATGACCGGAGAAAACCGGACAATTATTCTTTATAACCTTCGTTCAGATAC<br>AAATGAATTTATTGGGAAATCTGATGGGTTTATCCCTGCTAATACGGGCTTGCCTGCTTACA<br>GTACCGATATCGCGCCCCCAAAAGTGACGGCAGGTTTTGTGGCTGTTTTCGATGCACAGA<br>CGAATAAATGGTCGCGGGTGGAGGACTACCGCGGGACAACCGTCTATGACATCAGCACCG<br>GTAAGCCCGCTGTTATTGAAAAACTTGGCGCTCTGCCTGATAACGTTGTGTCGGTTGCTCC<br>TGACGGGGAGTATGTAAAATGGGATGGCGCTAAGTGGATCCACGATGCCGAAGCGGAAAA<br>AACATTTCGTCAGGGGCAGGCGGCGCAGGAAAAATCAAACCTGCTGATGATTGCAACATC<br>GGCTATTGCCCCCCTGCAGGATGCCGTTGATCTGGATATGGCAACGGAAGACGAAGCGAC<br>CGCGCTTAATGAATGGAAAAAATACCGGGTCATGCTCAACAGAGTCAAACCCGAAGATGCC<br>CCCGATATCACATGGCCGGAACTGCCCGCATAAAACTATATTGTGAGGCTTGCATAATGGCA<br>TTCAGAATGAGTGAACAACCACGGACCATAAAAATTTATAATCTGCTGGCCGGAACTAATGA<br>ATTTATTGGTGAAGGTGACGCATATATTCCGCCTCATACCGGTCTGCCTGCAAACAGTACCG<br>ATATTGCACCGCCAGATATTCCGGCTGGCTTTGTGGCTGTTTTCAACAGTGATGAGGCATC<br>GTGGCATCTCGTTGAAGACCATCGGGGTAAAACCGTCTATGACGTGGCTTCCGGCGACGC<br>GTTATTTATTTCTGAACTCGGTCCGTTACCGGAAAATTTTACCTGGTTATCGCCGGGAGGGG<br>AATATCAGAAGTGGAACGGCACAGCCTGGGTGAAGGATACGGAAGCAGAAAAACTGTTCC<br>GGATCCGGGAGGCGGAAGAAACAAAAAAAAGCCTGATGCAGGTAGCCAGTGAGCATATTG<br>CGCCGCTTCAGGATGCTGCAGATCTGGAAATTGCAACGAAGGAAGAAACCTCGTTGCTGG<br>AAGCCTGGAAGAAGTATCGGGTGTTGCTGAACCGTGTTGATACATCAACTGCACCTGATAT<br>TGAGTGGCCTGCTGTCCCTGTTATGGAGTAATGACGCATCCTCACGATAATATCCGGGTAG<br>GACGAACAATAAGGCCGCAAATCGCGGCCTTTTTTATTGATAACAAAAGGACAGTTTTCCCT<br>TTGATATGTAACGGTGAACAGTTGTTCTACTTTTGTTTGTTAGTCTTGATGCTTCACTGATAG<br>ATACAAGAGCCATAAGAACC |
| p522 | TCAGATCCTTCCGTATTTAGCCAGTATGTTCTCTAGTGTGGTTCGTTGTTTTTGCGTGAGCC<br>ATGAGAACGAACCATTGAGATCATACTTACTTTGCATGTCACTCAAAAATTTTGCCTCAAAAC<br>TGGTGAGCTGAATTTTTGCAGTTAAAGCATCGTGTAGTGTTTTTCTTAGTCCGTTACGTAGG<br>TAGGAATCTGATGTAATGGTTGTTGGTATTTTGTCACCATTCATTTTTATCTGGTTGTTCTCAA<br>GTTCGGTTACGAGATCCATTTGTCTATCTAGTTCAACTTGGAAAATCAACGTATCAGTCGGG<br>CGGCCTCGCTTATCAACCACCAATTTCATATTGCTGTAAGTGTTTAAATCTTTACTTATTGGT<br>TTCAAAACCCATTGGTTAAGCCTTTTAAACTCATGGTAGTTATTTTCAAGCATTAACATGAAC<br>TTAAATTCATCAAGGCTAATCTCTATATTTGCCTTGTGAGTTTTCTTTTGTGTTAGTTCTTTTA<br>ATAACCACTCATAAATCCTCATAGAGTATTTGTTTTCAAAAGACTTAACATGTTCCAGATTATA<br>TTTTATGAATTTTTTTAACTGGAAAAGATAAGGCAATATCTCTTCACTAAAAACTAATTCTAATT<br>TTTCGCTTGAGAACTTGGCATAGTTTGTCCACTGGAAAATCTCAAAGCCTTTAACCAAAGG<br>ATTCCTGATTTCCACAGTTCTCGTCATCAGCTCTCTGGTTGCTTTAGCTAATACACCATAAG<br>CATTTTCCCTACTGATGTTCATCATCTGAGCGTATTGGTTATAAGTGAACGATACCGTCCGTT<br>CTTTCCTTGTAGGGTTTTCAATCGTGGGGTTGAGTAGTGCCACACAGCATAAAATTAGCTTG<br>GTTTCATGCTCCGTTAAGTCATAGCGACTAATCGCTAGTTCATTTGCTTTGAAAACAACTAAT |

| | |
|---|---|
| | TCAGACATACATCTCAATTGGTCTAGGTGATTTTAATCACTATACCAATTGAGATGGGCTAGT<br>CAATGATAATTACTAGTCCTTTTCCTTTGAGTTGTGGGTATCTGTAAATTCTGCTAGACCTTT<br>GCTGGAAAACTTGTAAATTCTGCTAGACCCTCTGTAAATTCCGCTAGACCTTTGTGTGTTTT<br>TTTTGTTTATATTCAAGTGGTTATAATTTATAGAATAAAGAAAGAATAAAAAAAGATAAAAAGAA<br>TAGATCCCAGCCCTGTGTATAACTCACTACTTTAGTCAGTTCCGCAGTATTACAAAAGGATG<br>TCGCAAACGCTGTTTGCTCCTCTACAAAACAGACCTTAAAACCCTAAAGGCTTAAGTAGCA<br>CCCTCGCAAGCTCGGTTGCGGCCGCAATCGGGCAAATCGCTGAATATTCCTTTTGTCTCC<br>GACCATCAGGCACCTGAGTCGCTGTCTTTTTCGTGACATTCAGTTCGCTGCGCTCACGGC<br>TCTGGCAGTGAATGGGGGTAAATGGCACTACAGGCGCCTTTTATGGATTCATGCAAGGAAA<br>CTACCCATAATACAAGAAAAGCCCGTCACGGGCTTCTCAGGGCGTTTTATGGCGGGTCTG<br>CTATGTGGTGCTATCTGACTTTTTGCTGTTCAGCAGTTCCTGCCCTCTGATTTTCCAGTCTG<br>ACCACTTCGGATTATCCCGTGACAGGTCATTCAGACTGGCTAATGCACCCAGTAAGGCAGC<br>GGTATCATCAACGGGGTCTGACGCTCAGTGGAACGAAAACTCACGTTAAGGGATTTTGGT<br>CATGAGATTATCAAAAAGGATCTTCACCTAGATCCTTTTAAATTAAAAATGAAGTTTTAAATCA<br>ATCTAAAGTATATATGAGTAAACTTGGTCTGACAGTTACGTTTCCACAACCAATTAACCAATT<br>CTGATTTAGAAAAACTCATCGAGCATCAAATGAAACTGCAATTTATTCATATCAGGATTATCAA<br>TACCATATTTTTGAAAAAGCCGTTTCTGTAATGAAGGAGAAAACTCACCGAGGCAGTTCCAT<br>AGGATGGCAAGATCCTGGTATCGGTCTGCGATTCCGACTCGTCCAACATCAATACAACCTA<br>TTAATTTCCCCTCGTCAAAAATAAGGTTATCAAGTGAGAAATCACCATGAGTGACGACTGAA<br>TCCGGTGAGAATGGCAAAAGCTTATGCATTTCTTTCCAGACTTGTTCAACAGGCCAGCCAT<br>TACGCTCGTCATCAAAATCACTCGCATCAACCAAACCGTTATTCATTCGTGATTGCGCCTGA<br>GCGAGACGAAATACGCGATCGCTGTTAAAAGGACAATTACAAACAGGAATCGAATGCAACC<br>GGCGCAGGAACACTGCCAGCGCATCAACAATATTTTCACCTGAATCAGGATATTCTTCTAAT<br>ACCTGGAATGCTGTTTTCCCGGGGATCGCAGTGGTGAGTAACCATGCATCATCAGGAGTA<br>CGGATAAAATGCTTGATGGTCGGAAGAGGCATAAATTCCGTCAGCCAGTTTAGTCTGACCA<br>TCTCATCTGTAACATCATTGGCAACGCTACCTTTGCCATGTTTCAGAAACAACTCTGGCGCA<br>TCGGGCTTCCCATACAATCGATAGATTGTCGCACCTGATTGCCCGACATTATCGCGAGCCC<br>ATTTATACCCATATAAATCAGCATCCATGTTGGAATTTAATCGCGGCCTCGAGCAAGACGTTT<br>CCCGTTGAATATGGCTCATAACACCCCTTGTATTACTGTTTATGTAAGCAGACAGTTTTATTG<br>TTCATGATGATATATTTTTATCTTGTGCAATGTAACATCAGAGATTTTGAGACACAACGTGGC<br>TTTCCCTGCAGGATTTCGGAGGCCTGCGTTATCCCCTGATTCTGTGGATAACCGTATTACC<br>GCCTTTGAGTGAGCTGATACCGCTCGCCGCAGCCGAACGCCGACTAGTGGATTTTACGGC<br>TAGCTCAGTCCTAGGTACAATGCTAGCGAATTCATTAAAGAGGAGAAAGGTACCCATGGCA<br>CGTACCCCGAGCCGTAGCAGCATTGGTAGCCTGCGTAGTCCGCATACCCATAAAGCAATTC<br>TGACCAGCACCATTGAAATCCTGAAAGAATGTGGTTATAGCGGTCTGAGCATTGAAAGCGT<br>TGCACGTCGTGCCGGTGCAAGCAAACCGACCATTTATCGTTGGTGGACCAATAAAGCAGC<br>ACTGATTGCCGAAGTGTATGAAAATGAAAGCGAACAGGTGCGTAAATTTCCGGATCTGGGT<br>AGCTTTAAAGCCGATCTGGATTTTCTGCTGCGTAATCTGTGGAAAGTTTGGCGTGAAACCA<br>TTTGTGGTGAAGCATTTCGTTGTGTTATTGCAGAAGCACAGCTGGACCCTGCAACCCTGAC<br>CCAGCTGAAAGATCAGTTTATGGAACGTCGTCGTGAGATGCCGAAAAAACTGGTTGAAAAT<br>GCCATTAGCAATGGTGAACTGCCGAAAGATACCAATCGTGAACTGCTGCTGGATATGATTTT<br>TGGTTTTTGTTGGTATCGCCTGCTGACCGAACAGCTGACCGTTGAACAGGATATTGAAGAA<br>TTTACCTTCCTGCTAATTAATGGTGTTTGTCCGGGTACACAGCGTTAACTAGGGCCCATACC<br>CCCAATTATTGAAGGCCGCTAACGCGGCCTTTTTTTGTTTCTGGTCTGCCCGACGTACGGT<br>GAATCTGATTCGTTACCAATTGACATGATACGAAACGTACCGTATCGTTAAGGTTACTAGATT<br>AAAGAGGAGAAATACTAGATGGCAGTAAAGATTTCAGGAGTCCTGAAAGACGGCACAGGA<br>AAACCGGTACAGAACTGCACCATTCAGCTGAAAGCCAGACGTAACAGCACCACGGTGGTG<br>GTGAACACGGTGGGCTCAGAGAATCCGGATGAAGCCGGGCGTTACAGCATGGATGTGGA |

| | |
|---|---|
| | GTACGGTCAGTACAGTGTCATCCTGCAGGTTGACGGTTTTCCACCATCGCACGCCGGGAC<br>CATCACCGTGTATGAAGATTCACAACCGGGGACGCTGAATGATTTTCTCTGTGCCATGACG<br>GAGGATGATGCCCGGCCGGAGGTGCTGCGTCGTCTTGAACTGATGGTGGAAGAGGTGGC<br>GCGTAACGCGTCCGTGGTGGCACAGAGTACGGCAGACGCGAAGAAATCAGCCGGCGATG<br>CCAGTGCATCAGCTGCTCAGGTCGCGGCCCTTGTGACTGATGCAACTGACTCAGCACGC<br>GCCGCCAGCACGTCCGCCGGACAGGCTGCATCGTCAGCTCAGGAAGCGTCCTCCGGCG<br>CAGAAGCGGCATCAGCAAAGGCCACTGAAGCGGAAAAAAGTGCCGCAGCCGCAGAGTCC<br>TCAAAAAACGCGGCGGCCACCAGTGCCGGTGCGGCGAAAACGTCAGAAACGAATGCTGC<br>AGCGTCACAACAATCAGCCGCCACGTCTGCCTCCACCGCGGCCACGAAAGCGTCAGAGG<br>CCGCCACTTCAGCACGAGATGCGGTGGCCTCAAAAGAGGCAGCAAAATCATCAGAAACGA<br>ACGCATCATCAAGTGCCGGTCGTGCAGCTTCCTCGGCAACGGCGGCAGAAAATTCTGCCA<br>GGGCGGCAAAAACGTCCGAGACGAATGCCAGGTCATCTGAAACAGCAGCGGAACGGAGC<br>GCCTCTGCCGCGGCAGACGCAAAAACAGCGGCGGCGGGGAGTGCGTCAACGGCATCCA<br>CGAAGGCGACAGAGGCTGCGGGAAGTGCGGTATCAGCATCGCAGAGCAAAAGTGCGGCA<br>GAAGCGGCGGCAATACGTGCAAAAAATTCGGCAAAACGTGCAGAAGATATAGCTTCAGCT<br>GTCGCGCTTGAGGATGCGGACACAACGAGAAAGGGGATAGTGCAGCTCAGCAGTGCAAC<br>CAACAGCACGTCTGAAACGCTTGCTGCAACGCCAAAGGCGGTTAAGGTGGTAATGGATGA<br>GACTAATCGTCGCGTTCCGGCATCACGAAAAGTGAACGGCCATGCCCTGAATGGAGATATC<br>AATGTCACTTCACGGGATATTTTTGACGGCCAGGTTATAGCGATTGGTGCAAATAAGAATCT<br>GGATGATTACCAGGTACCGGGGCTTTATTTTCAGGAAGCGAACAACAATACCAGTGCAGCA<br>ATGAATTACCCGGAGAATAGCGCGGGTTCTCTGATGGTACTGAGAGGTGCCGGAGTCACT<br>CAGGTTTATCGTGTGTACAACAGCTCGCGCAGTTATTCGCGCAGCAAGTATTCAACGCTGG<br>CATGGACGCCGTGGATGCCAGAAGATTCTTACCCTGTCGGCGCACCTATCCCCTGGCCAT<br>CGGATGTTACCCCGACAGGGTACGCCTTAATGCAGGGGCAGCCCTTTGATAAAGCGGTCT<br>ATCCATTGCTAGCGATTGCCTATCCTGCGGGGATTATCCCGGACATGCGAGGCCAGACGAT<br>TAAGGGTAAACCGAACGGTCGCGCGGTACTCTCGTATGAACAGGATGGTGTTATATCGCAT<br>ACCCACGGAGCCAGTATTTCCGATACCGATTTGGGGACGAAATACACCAGCTCTTTTGATT<br>ATGGTTCAAAACCAACAACCAGTTTTGACTACGGCAATAAATCCTCCACTGAGGGTGGGTG<br>GCACGCACATAACTTTCGTTATTGCGCAACGTCTGCATACCGGGATACCCCCGGTCAGGG<br>GCTGGGGATGCATTCGTCTAATGTTTCATGGGCGGCGGGAGATCGCATTGAGGGAAGCGG<br>TAATCATGCTCATGTGACATGGATCGGCCCTCATGATCACTGGGTGGGTATTGGTGCGCAT<br>AACCATTATGTGGTTATGGGCTATCACGGACATACAGCGACCGTTCATGCCGCAGGAAATG<br>CGGAAAATACCGTTAAAAATATTGCGTTTAACTACATTGTGAGGCTTGCCTGATGACTTTTG<br>AAATGACCGGAGAAAACCGGACAATTACCATCTATAACCTGCGTGCTGATACAAATGAATTT<br>ATCGGGAAAAGTGATGGGTTTATCCCTGCTAATACCGGTTTGCCTGCTAACAGTACCAATAT<br>TGCGCCACCGCCGATGAAAGCCGGTTTTGTCGCTGTATTTAATTCTGCGTCAGAAAAATGG<br>TCACTTGTTGAAGACCATCGCGGGAAAATTGTTTACGACATTCTCACCGGGAAATCCATCA<br>CGATTGATGAATTAGGTCAGTTACCTGACGACGTTGTTTCCGTTGCGCCGGAAGGCCATTT<br>TGTTAAATGGAATGGTAAAAAATGGGTGCATGATGCTGACGCAGAAAAAACGGCACAGATT<br>ACACAGGCTACACAGCAAAAAGACAGTCTTCTGGCGCTGGCTGCATCAAAAATTGCCCCA<br>TTACAGGATGCTGTTGATCTGGATATTGCAACGGAAGAGGAAACAGCGCTTTTGCTGGCGT<br>GGAAAAAATACAGGGTTTTGATTAATCGTATTAAGCCAGAAGATGCGCCAGATATTGACTGG<br>CCGGAGGTTCCGGGCGATGTGGCGTGAAACTATATTGTGAGGCTTGCATAATGGCATTCAG<br>AATGAGTGAACAACCACGGACCATAAAAATTTATAATCTGCTGGCCGGAACTAATGAATTTAT<br>TGGTGAAGGTGACGCATATATTCCGCCTCATACCGGTCTGCCTGCAAACAGTACCGATATT<br>GCACCGCCAGATATTCCGGCTGGCTTTGTGGCTGTTTTCAACAGTGATGAGGCATCGTGG<br>CATCTCGTTGAAGACCATCGGGGTAAAACCGTCTATGACGTGGCTTCCGGCGACGCGTTA<br>TTTATTTCTGAACTCGGTCCGTTACCGGAAAATTTTACCTGGTTATCGCCGGGAGGGGAATA |

| | |
|---|---|
| | TCAGAAGTGGAACGGCACAGCCTGGGTGAAGGATACGGAAGCAGAAAAACTGTTCCGGA<br>TCCGGGAGGCGGAAGAAACAAAAAAAAGCCTGATGCAGGTAGCCAGTGAGCATATTGCGC<br>CGCTTCAGGATGCTGCAGATCTGGAAATTGCAACGAAGGAAGAAACCTCGTTGCTGGAAG<br>CCTGGAAGAAGTATCGGGTGTTGCTGAACCGTGTTGATACATCAACTGCACCTGATATTGA<br>GTGGCCTGCTGTCCCTGTTATGGAGTAATGACGCATCCTCACGATAATATCCGGGTAGGAC<br>GAACAATAAGGCCGCAAATCGCGGCCTTTTTATTGATAACAAAAGGACAGTTTTCCCTTTG<br>ATATGTAACGGTGAACAGTTGTTCTACTTTTGTTTGTTAGTCTTGATGCTTCACTGATAGATA<br>CAAGAGCCATAAGAACC |
| p363 | TCAGATCCTTCCGTATTTAGCCAGTATGTTCTCTAGTGTGGTTCGTTGTTTTTGCGTGAGCC<br>ATGAGAACGAACCATTGAGATCATACTTACTTTGCATGTCACTCAAAAATTTTGCCTCAAAAC<br>TGGTGAGCTGAATTTTTGCAGTTAAAGCATCGTGTAGTGTTTTTCTTAGTCCGTTACGTAGG<br>TAGGAATCTGATGTAATGGTTGTTGGTATTTTGTCACCATTCATTTTTATCTGGTTGTTCTCAA<br>GTTCGGTTACGAGATCCATTTGTCTATCTAGTTCAACTTGGAAAATCAACGTATCAGTCGGG<br>CGGCCTCGCTTATCAACCACCAATTTCATATTGCTGTAAGTGTTTAAATCTTTACTTATTGGT<br>TTCAAAACCCATTGGTTAAGCCTTTTAAACTCATGGTAGTTATTTTCAAGCATTAACATGAAC<br>TTAAATTCATCAAGGCTAATCTCTATATTTGCCTTGTGAGTTTTCTTTTGTGTTAGTTCTTTTA<br>ATAACCACTCATAAATCCTCATAGAGTATTTGTTTTCAAAAGACTTAACATGTTCCAGATTATA<br>TTTTATGAATTTTTTTAACTGGAAAAGATAAGGCAATATCTCTTCACTAAAAACTAATTCTAATT<br>TTTCGCTTGAGAACTTGGCATAGTTTGTCCACTGGAAAATCTCAAAGCCTTTAACCAAAGG<br>ATTCCTGATTTCCACAGTTCTCGTCATCAGCTCTCTGGTTGCTTTAGCTAATACACCATAAG<br>CATTTTCCCTACTGATGTTCATCATCTGAGCGTATTGGTTATAAGTGAACGATACCGTCCGTT<br>CTTTCCTTGTAGGGTTTTCAATCGTGGGGTTGAGTAGTGCCACACAGCATAAAATTAGCTTG<br>GTTTCATGCTCCGTTAAGTCATAGCGACTAATCGCTAGTTCATTTGCTTTGAAAACAACTAAT<br>TCAGACATACATCTCAATTGGTCTAGGTGATTTTAATCACTATACCAATTGAGATGGGCTAGT<br>CAATGATAATTACTAGTCCTTTTCCTTTGAGTTGTGGGTATCTGTAAATTCTGCTAGACCTTT<br>GCTGGAAAACTTGTAAATTCTGCTAGACCCTCTGTAAATTCCGCTAGACCTTTGTGTGTTTT<br>TTTTGTTTATATTCAAGTGGTTATAATTTATAGAATAAAGAAAGAATAAAAAAAGATAAAAAGAA<br>TAGATCCCAGCCCTGTGTATAACTCACTACTTTAGTCAGTTCCGCAGTATTACAAAAGGATG<br>TCGCAAACGCTGTTTGCTCCTCTACAAAACAGACCTTAAAACCCTAAAGGCTTAAGTAGCA<br>CCCTCGCAAGCTCGGTTGCGGCCGCAATCGGGCAAATCGCTGAATATTCCTTTTGTCTCC<br>GACCATCAGGCACCTGAGTCGCTGTCTTTTTCGTGACATTCAGTTCGCTGCGCTCACGGC<br>TCTGGCAGTGAATGGGGGTAAATGGCACTACAGGCGCCTTTTATGGATTCATGCAAGGAAA<br>CTACCCATAATACAAGAAAAGCCCGTCACGGGCTTCTCAGGGCGTTTTATGGCGGGTCTG<br>CTATGTGGTGCTATCTGACTTTTTGCTGTTCAGCAGTTCCTGCCCTCTGATTTTCCAGTCTG<br>ACCACTTCGGATTATCCCGTGACAGGTCATTCAGACTGGCTAATGCACCCAGTAAGGCAGC<br>GGTATCATCAACGGGGTCTGACGCTCAGTGGAACGAAAACTCACGTTAAGGGATTTTGGT<br>CATGAGATTATCAAAAAGGATCTTCACCTAGATCCTTTTAAATTAAAAATGAAGTTTTAAATCA<br>ATCTAAAGTATATATGAGTAAACTTGGTCTGACAGTTACGTTTCCACAACCAATTAACCAATT<br>CTGATTTAGAAAAACTCATCGAGCATCAAATGAAACTGCAATTTATTCATATCAGGATTATCAA<br>TACCATATTTTTGAAAAAGCCGTTTCTGTAATGAAGGAGAAAACTCACCGAGGCAGTTCCAT<br>AGGATGGCAAGATCCTGGTATCGGTCTGCGATTCCGACTCGTCCAACATCAATACAACCTA<br>TTAATTTCCCCTCGTCAAAAATAAGGTTATCAAGTGAGAAATCACCATGAGTGACGACTGAA<br>TCCGGTGAGAATGGCAAAAGCTTATGCATTTCTTTCCAGACTTGTTCAACAGGCCAGCCAT<br>TACGCTCGTCATCAAAATCACTCGCATCAACCAAACCGTTATTCATTCGTGATTGCGCCTGA<br>GCGAGACGAAATACGCGATCGCTGTTAAAAGGACAATTACAAACAGGAATCGAATGCAACC<br>GGCGCAGGAACACTGCCAGCGCATCAACAATATTTTCACCTGAATCAGGATATTCTTCTAAT<br>ACCTGGAATGCTGTTTTCCCGGGGATCGCAGTGGTGAGTAACCATGCATCATCAGGAGTA<br>CGGATAAAATGCTTGATGGTCGGAAGAGGCATAAATTCCGTCAGCCAGTTTAGTCTGACCA |

| | |
|---|---|
| | TCTCATCTGTAACATCATTGGCAACGCTACCTTTGCCATGTTTCAGAAACAACTCTGGCGCA<br>TCGGGCTTCCCATACAATCGATAGATTGTCGCACCTGATTGCCCGACATTATCGCGAGCCC<br>ATTTATACCCATATAAATCAGCATCCATGTTGGAATTTAATCGCGGCCTCGAGCAAGACGTTT<br>CCCGTTGAATATGGCTCATAACACCCCTTGTATTACTGTTTATGTAAGCAGACAGTTTTATTG<br>TTCATGATGATATATTTTTATCTTGTGCAATGTAACATCAGAGATTTTGAGACACAACGTGGC<br>TTTCCCTGCAGGATTTCGGAGGCCTGCGTTATCCCCTGATTCTGTGGATAACCGTATTACC<br>GCCTTTGAGTGAGCTGATACCGCTCGCCGCAGCCGAACGCCGACTAGTGGATTTTACGGC<br>TAGCTCAGTCCTAGGTACAATGCTAGCGAATTCATTAAAGAGGAGAAAGGTACCCATGGCA<br>CGTACCCCGAGCCGTAGCAGCATTGGTAGCCTGCGTAGTCCGCATACCCATAAAGCAATTC<br>TGACCAGCACCATTGAAATCCTGAAAGAATGTGGTTATAGCGGTCTGAGCATTGAAAGCGT<br>TGCACGTCGTGCCGGTGCAAGCAAACCGACCATTTATCGTTGGTGGACCAATAAAGCAGC<br>ACTGATTGCCGAAGTGTATGAAAATGAAAGCGAACAGGTGCGTAAATTTCCGGATCTGGGT<br>AGCTTTAAAGCCGATCTGGATTTTCTGCTGCGTAATCTGTGGAAAGTTTGGCGTGAAACCA<br>TTTGTGGTGAAGCATTTCGTTGTGTTATTGCAGAAGCACAGCTGGACCCTGCAACCCTGAC<br>CCAGCTGAAAGATCAGTTTATGGAACGTCGTCGTGAGATGCCGAAAAAACTGGTTGAAAAT<br>GCCATTAGCAATGGTGAACTGCCGAAAGATACCAATCGTGAACTGCTGCTGGATATGATTTT<br>TGGTTTTTGTTGGTATCGCCTGCTGACCGAACAGCTGACCGTTGAACAGGATATTGAAGAA<br>TTTACCTTCCTGCTAATTAATGGTGTTTGTCCGGGTACACAGCGTTAACTAGGGCCCATACC<br>CCCAATTATTGAAGGCCGCTAACGCGGCCTTTTTTTGTTTCTGGTCTGCCCGACGTACGGT<br>GAATCTGATTCGTTACCAATTGACATGATACGAAACGTACCGTATCGTTAAGGTTACTAGATT<br>AAAGAGGAGAAATACTAGATGGCAGTAAAGATTTCAGGAGTCCTGAAAGACGGCACAGGA<br>AAACCGGTACAGAACTGCACCATTCAGCTGAAAGCCAGACGTAACAGCACCACGGTGGTG<br>GTGAACACGGTGGGCTCAGAGAATCCGGATGAAGCCGGGCGTTACAGCATGGATGTGGA<br>GTACGGTCAGTACAGTGTCATCCTGCAGGTTGACGGTTTTCCACCATCGCACGCCGGGAC<br>CATCACCGTGTATGAAGATTCACAACCGGGGACGCTGAATGATTTTCTCTGTGCCATGACG<br>GAGGATGATGCCCGGCCGGAGGTGCTGCGTCGTCTTGAACTGATGGTGGAAGAGGTGGC<br>GCGTAACGCGTCCGTGGTGGCACAGAGTACGGCAGACGCGAAGAAATCAGCCGGCGATG<br>CCAGTGCATCAGCTGCTCAGGTCGCGGCCCTTGTGACTGATGCAACTGACTCAGCACGC<br>GCCGCCAGCACGTCCGCCGGACAGGCTGCATCGTCAGCTCAGGAAGCGTCCTCCGGCG<br>CAGAAGCGGCATCAGCAAAGGCCACTGAAGCGGAAAAAAGTGCCGCAGCCGCAGAGTCC<br>TCAAAAAACGCGGCGGCCACCAGTGCCGGTGCGGCGAAAACGTCAGAAACGAATGCTGC<br>AGCGTCACAACAATCAGCCGCCACGTCTGCCTCCACCGCGGCCACGAAAGCGTCAGAGG<br>CCGCCACTTCAGCACGAGATGCGGTGGCCTCAAAAGAGGCAGCAAAATCATCAGAAACGA<br>ACGCATCATCAAGTGCCGGTCGTGCAGCTTCCTCGGCAACGGCGGCAGAAAATTCTGCCA<br>GGGCGGCAAAAACGTCCGAGACGAATGCCAGGTCATCTGAAACAGCAGCGGAACGGAGC<br>GCCTCTGCCGCGGCAGACGCAAAAACAGCGGCGGCGGGGAGTGCGTCAACGGCATCCA<br>CGAAGGCGACAGAGGCTGCGGGAAGTGCGGTATCAGCATCGCAGAGCAAAAGTGCGGCA<br>GAAGCGGCGGCAATACGTGCAAAAAATTCGGCAAAACGTGCAGAAGATATAGCTTCAGCT<br>GTCGCGCTTGAGGATGCGGACACAACGAGAAAGGGGATAGTGCAGCTCAGCAGTGCAAC<br>CAACAGCACGTCTGAAACGCTTGCTGCAACGCCAAAGGCGGTTAAGGTGGTAATGGATGA<br>GACTAATCGTAAGGCACCTCTGGACAGTCCGGCACTGACCGGAACGCCAACAGCACCAA<br>CCGCGCTCAGGGGAACAAACAATACCCAGATTGCGAACACCGCTTTTGTACTGGCCGCGA<br>TTGCAGATGTTATCGACGCGTCACCTGACGCACTGAATACGCTGAATGAACTGGCCGCAG<br>CGCTCGGGAATGATCCAGATTTTGCTACCACCATGACTAACGCGCTTGCGGGTAAACAACC<br>GAAGAATGCGACACTGACGGCGCTGGCAGGGCTTTCCACGGCGAAAAATAAATTACCGTA<br>TTTTGCGGAAAATGATGCCGCCAGCCTGACTGAACTGACTCAGGTTGGCAGGGATATTCT<br>GGCAAAAAATTCCGTTGCAGATGTTCTTGAATACCTTGGGGCCGGTGAGAATTCGGCGGC<br>AAATGATGGCTTCAAATTCATCGGTCAGTGCCCAGACATCTTGACCCTGCGTACTATCGAG |

| | |
|---|---|
| | CCGGAAAAAAACGGTCAGCGTATCACCTTACGTCAACATACGATTGGCACTGGCTTAGGCG<br>GTGGCGTTTTCCGTGCAGTTCTGGACGGCACTGGCTATACCGATGACGACGGTGTGGTGA<br>TCAAAACCGCTGGGGGCAGCGTTTGGCTGCGTGTCAACGCTGACAAAGTTAACCCGTTCA<br>TGTTCGGTGCAACCGGAGTAGCGGACGACACCGCCGCCCTGCAAAAAATGCTGGAATGC<br>GGTCGTGCGGCGGAACTGGGGACTAACGTATGGAAAGCAAGCAATCTGGAACTGAACAA<br>CAAATCTTGCTCTCTGTCCGGCAGTGGCCTGCACGTTTCTCGTATTGAACAGATTTCCGGT<br>GCAACCGGAGCATTGTTAACCATCACCCAAGACTGTTCGCTGATTTACCTGTCCGATTGTG<br>GCCTGTACGGCGATGGCATCACCGCAGGCACGAGCGGTGTTACTATGGAAACGGGTAATC<br>CGGGTGGCGCTCCGTCTTACCCTTTCAATACCGCTCCGGACGTTCGTCGTGACCTGTACA<br>TCTCTAACGTGCACATCACGGGCTTCGACGAGCTGGGTTTTGATTATCCGGAAACCAATTT<br>CTCTGTTTCGACGCATGGCCTCTTCATCCGTAACATCAAAAAAACGGGTGCAAAGATTGGT<br>ACTACGGACTTCACTTGGACTAACCTGCAAATTGATACTTGCGGTCAGGAATGTCTGGTGC<br>TGGACGGTGCGGGTAACTGCCGTATTATTGGTGCAAAACTGATTTGGGCAGGTAGCGAAA<br>ACGAAACGCCATACTCTGGCCTGCGTATTAGCAACTCTCAAAATGTAAATATGACTGGCGTA<br>GAGTTACAAGACTGCGCGTATGATGGTTTATACATCAAGAACTCTACGGTTGCAATTTCAGG<br>CTTAAACACCAATCGCAATAGCGCATCCTCTAATCTGTCCTACCATAACATGGTATTCGAAAA<br>TTCTATTGTAACTGTTGATGGTTATGTGTGTCGTAACTACGCGGCGACTTCGCTGTACGACC<br>TGAACAGCCAAGCAGGCAACGTCCGTTGCATCGGTAGCGACAGCACCGTTTTAATCAACG<br>GCATCTACGAAAGCGAAGTCAATAGCGAGCGCCTGATGGGTGATAACAACCTGATCCAGC<br>CGTATAGTGGTGATCTGATCATTAACGGCCTGAAAAATTACTACACCTATACTGGTAGCGTAA<br>AAAACAACATTCCGACCTTCGACGGCGTTGTTACTACGGCAACCTATGTGAGCGCACCGTC<br>TATTCTGGGTCAGGGCAATATGCTCAAACTGACCCAGTCTAATAAAGACAAACTGTTATTTA<br>GCGATAAAGTTAGCCGTCATGGCTGTACCATCGGCTTAGTTCTGATTCCGTCCTTTACGGG<br>CGCGACCACTATGACGGCGTTCACGCTGGGTAGCGGTTACTCTCCATCCGGTAACTCCGC<br>CGTGATGCAGTTCATTGTTAACAGTTCCGGTGTACAAACCATTGCGATTTTATTATCCGGCG<br>ACGGTATTACCCAAACCCTGACCAGCGATCTGACCACGGAACAAGCACTGGCGAGCGGT<br>GGCGTGTATCATTTTGCAATGGGTTTTGCGCCGGGTCGTTTATGGTGGAGCATTATCGATAT<br>TAACACGGGCAGGCGTATTCGTCGCGCCTACCGTCAGCCGGATCTGCACGCGGCGTTCA<br>ACTCTATCTTCAACTCCGGCACGTCGTCTATTACCGCATTTAGCGGGCCACTGGCGGGCGA<br>CATTGCTTGCGAAGGTGCAGGTAGCCATGTATACGTTGGCGGTTTTTCGTCGGAATCTGAT<br>TACGCGGCTAGCCGTATGTATGGCCTGTTCACTCCGGTCGATCTGGACAAGCAGTATAGCT<br>TCCGTACCCTGAACGGTAACATTAACTATATTGTGAGGCTTGCATAATGGCATTCAGAATGA<br>GTGAACAACCACGGACCATAAAAATTTATAATCTGCTGGCCGGAACTAATGAATTTATTGGT<br>GAAGGTGACGCATATATTCCGCCTCATACCGGTCTGCCTGCAAACAGTACCGATATTGCAC<br>CGCCAGATATTCCGGCTGGCTTTGTGGCTGTTTTCAACAGTGATGAGGCATCGTGGCATCT<br>CGTTGAAGACCATCGGGGTAAAACCGTCTATGACGTGGCTTCCGGCGACGCGTTATTTATT<br>TCTGAACTCGGTCCGTTACCGGAAAATTTTACCTGGTTATCGCCGGGAGGGGAATATCAGA<br>AGTGGAACGGCACAGCCTGGGTGAAGGATACGGAAGCAGAAAAACTGTTCCGGATCCGG<br>GAGGCGGAAGAAACAAAAAAAAGCCTGATGCAGGTAGCCAGTGAGCATATTGCGCCGCTT<br>CAGGATGCTGCAGATCTGGAAATTGCAACGAAGGAAGAAACCTCGTTGCTGGAAGCCTGG<br>AAGAAGTATCGGGTGTTGCTGAACCGTGTTGATACATCAACTGCACCTGATATTGAGTGGC<br>CTGCTGTCCCTGTTATGGAGTAATGACGCATCCTCACGATAATATCCGGGTAGGACGAACA<br>ATAAGGCCGCAAATCGCGGCCTTTTTTATTGATAACAAAAGGACAGTTTTCCCTTTGATATGT<br>AACGGTGAACAGTTGTTCTACTTTTGTTTGTTAGTCTTGATGCTTCACTGATAGATACAAGA<br>GCCATAAGAACC |
| p871 | TCAGATCCTTCCGTATTTAGCCAGTATGTTCTCTAGTGTGGTTCGTTGTTTTTGCGTGAGCC<br>ATGAGAACGAACCATTGAGATCATACTTACTTTGCATGTCACTCAAAAATTTTGCCTCAAAAC<br>TGGTGAGCTGAATTTTTGCAGTTAAAGCATCGTGTAGTGTTTTTCTTAGTCCGTTACGTAGG |

| | |
|---|---|
| | TAGGAATCTGATGTAATGGTTGTTGGTATTTTGTCACCATTCATTTTTATCTGGTTGTTCTCAA<br>GTTCGGTTACGAGATCCATTTGTCTATCTAGTTCAACTTGGAAAATCAACGTATCAGTCGGG<br>CGGCCTCGCTTATCAACCACCAATTTCATATTGCTGTAAGTGTTTAAATCTTTACTTATTGGT<br>TTCAAAACCCATTGGTTAAGCCTTTTAAACTCATGGTAGTTATTTTCAAGCATTAACATGAAC<br>TTAAATTCATCAAGGCTAATCTCTATATTTGCCTTGTGAGTTTTCTTTTGTGTTAGTTCTTTTA<br>ATAACCACTCATAAATCCTCATAGAGTATTTGTTTTCAAAAGACTTAACATGTTCCAGATTATA<br>TTTTATGAATTTTTTTAACTGGAAAAGATAAGGCAATATCTCTTCACTAAAAACTAATTCTAATT<br>TTTCGCTTGAGAACTTGGCATAGTTTGTCCACTGGAAAATCTCAAAGCCTTTAACCAAAGG<br>ATTCCTGATTTCCACAGTTCTCGTCATCAGCTCTCTGGTTGCTTTAGCTAATACACCATAAG<br>CATTTTCCCTACTGATGTTCATCATCTGAGCGTATTGGTTATAAGTGAACGATACCGTCCGTT<br>CTTTCCTTGTAGGGTTTTCAATCGTGGGGTTGAGTAGTGCCACACAGCATAAAATTAGCTTG<br>GTTTCATGCTCCGTTAAGTCATAGCGACTAATCGCTAGTTCATTTGCTTTGAAAACAACTAAT<br>TCAGACATACATCTCAATTGGTCTAGGTGATTTTAATCACTATACCAATTGAGATGGGCTAGT<br>CAATGATAATTACTAGTCCTTTTCCTTTGAGTTGTGGGTATCTGTAAATTCTGCTAGACCTTT<br>GCTGGAAAACTTGTAAATTCTGCTAGACCCTCTGTAAATTCCGCTAGACCTTTGTGTGTTTT<br>TTTTGTTTATATTCAAGTGGTTATAATTTATAGAATAAAGAAAGAATAAAAAAAGATAAAAAGAA<br>TAGATCCCAGCCCTGTGTATAACTCACTACTTTAGTCAGTTCCGCAGTATTACAAAAGGATG<br>TCGCAAACGCTGTTTGCTCCTCTACAAAACAGACCTTAAAACCCTAAAGGCTTAAGTAGCA<br>CCCTCGCAAGCTCGGTTGCGGCCGCAATCGGGCAAATCGCTGAATATTCCTTTTGTCTCC<br>GACCATCAGGCACCTGAGTCGCTGTCTTTTTCGTGACATTCAGTTCGCTGCGCTCACGGC<br>TCTGGCAGTGAATGGGGGTAAATGGCACTACAGGCGCCTTTTATGGATTCATGCAAGGAAA<br>CTACCCATAATACAAGAAAAGCCCGTCACGGGCTTCTCAGGGCGTTTTATGGCGGGTCTG<br>CTATGTGGTGCTATCTGACTTTTTGCTGTTCAGCAGTTCCTGCCCTCTGATTTTCCAGTCTG<br>ACCACTTCGGATTATCCCGTGACAGGTCATTCAGACTGGCTAATGCACCCAGTAAGGCAGC<br>GGTATCATCAACGGGGTCTGACGCTCAGTGGAACGAAAACTCACGTTAAGGGATTTTGGT<br>CATGAGATTATCAAAAAGGATCTTCACCTAGATCCTTTTAAATTAAAAATGAAGTTTTAAATCA<br>ATCTAAAGTATATATGAGTAAACTTGGTCTGACAGTTACGTTTCCACAACCAATTAACCAATT<br>CTGATTTAGAAAAACTCATCGAGCATCAAATGAAACTGCAATTTATTCATATCAGGATTATCAA<br>TACCATATTTTTGAAAAAGCCGTTTCTGTAATGAAGGAGAAAACTCACCGAGGCAGTTCCAT<br>AGGATGGCAAGATCCTGGTATCGGTCTGCGATTCCGACTCGTCCAACATCAATACAACCTA<br>TTAATTTCCCCTCGTCAAAAATAAGGTTATCAAGTGAGAAATCACCATGAGTGACGACTGAA<br>TCCGGTGAGAATGGCAAAAGCTTATGCATTTCTTTCCAGACTTGTTCAACAGGCCAGCCAT<br>TACGCTCGTCATCAAAATCACTCGCATCAACCAAACCGTTATTCATTCGTGATTGCGCCTGA<br>GCGAGACGAAATACGCGATCGCTGTTAAAAGGACAATTACAAACAGGAATCGAATGCAACC<br>GGCGCAGGAACACTGCCAGCGCATCAACAATATTTTCACCTGAATCAGGATATTCTTCTAAT<br>ACCTGGAATGCTGTTTTCCCGGGGATCGCAGTGGTGAGTAACCATGCATCATCAGGAGTA<br>CGGATAAAATGCTTGATGGTCGGAAGAGGCATAAATTCCGTCAGCCAGTTTAGTCTGACCA<br>TCTCATCTGTAACATCATTGGCAACGCTACCTTTGCCATGTTTCAGAAACAACTCTGGCGCA<br>TCGGGCTTCCCATACAATCGATAGATTGTCGCACCTGATTGCCCGACATTATCGCGAGCCC<br>ATTTATACCCATATAAATCAGCATCCATGTTGGAATTTAATCGCGGCCTCGAGCAAGACGTTT<br>CCCGTTGAATATGGCTCATAACACCCCTTGTATTACTGTTTATGTAAGCAGACAGTTTTATTG<br>TTCATGATGATATATTTTTATCTTGTGCAATGTAACATCAGAGATTTTGAGACACAACGTGGC<br>TTTCCCTGCAGGATTTCGGAGGCCTGCGTTATCCCCTGATTCTGTGGATAACCGTATTACC<br>GCCTTTGAGTGAGCTGATACCGCTCGCCGCAGCCGAACGCCGACTAGTGGATTTTACGGC<br>TAGCTCAGTCCTAGGTACAATGCTAGCGAATTCATTAAAGAGGAGAAAGGTACCCATGGCA<br>CGTACCCCGAGCCGTAGCAGCATTGGTAGCCTGCGTAGTCCGCATACCCATAAAGCAATTC<br>TGACCAGCACCATTGAAATCCTGAAAGAATGTGGTTATAGCGGTCTGAGCATTGAAAGCGT<br>TGCACGTCGTGCCGGTGCAAGCAAACCGACCATTTATCGTTGGTGGACCAATAAAGCAGC |

| | |
|---|---|
| | ACTGATTGCCGAAGTGTATGAAAATGAAAGCGAACAGGTGCGTAAATTTCCGGATCTGGGT<br>AGCTTTAAAGCCGATCTGGATTTTCTGCTGCGTAATCTGTGGAAAGTTTGGCGTGAAACCA<br>TTTGTGGTGAAGCATTTCGTTGTGTTATTGCAGAAGCACAGCTGGACCCTGCAACCCTGAC<br>CCAGCTGAAAGATCAGTTTATGGAACGTCGTCGTGAGATGCCGAAAAAACTGGTTGAAAAT<br>GCCATTAGCAATGGTGAACTGCCGAAAGATACCAATCGTGAACTGCTGCTGGATATGATTTT<br>TGGTTTTTGTTGGTATCGCCTGCTGACCGAACAGCTGACCGTTGAACAGGATATTGAAGAA<br>TTTACCTTCCTGCTAATTAATGGTGTTTGTCCGGGTACACAGCGTTAACTAGGGCCCATACC<br>CCCAATTATTGAAGGCCGCTAACGCGGCCTTTTTTTGTTTCTGGTCTGCCCGACGTACGGT<br>GAATCTGATTCGTTACCAATTGACATGATACGAAACGTACCGTATCGTTAAGGTTACTAGATT<br>AAAGAGGAGAAATACTAGATGGCAGTAAAGATTTCAGGAGTCCTGAAAGACGGCACAGGA<br>AAACCGGTACAGAACTGCACCATTCAGCTGAAAGCCAGACGTAACAGCACCACGGTGGTG<br>GTGAACACGGTGGGCTCAGAGAATCCGGATGAAGCCGGGCGTTACAGCATGGATGTGGA<br>GTACGGTCAGTACAGTGTCATCCTGCAGGTTGACGGTTTTCCACCATCGCACGCCGGGAC<br>CATCACCGTGTATGAAGATTCACAACCGGGGACGCTGAATGATTTTCTCTGTGCCATGACG<br>GAGGATGATGCCCGGCCGGAGGTGCTGCGTCGTCTTGAACTGATGGTGGAAGAGGTGGC<br>GCGTAACGCGTCCGTGGTGGCACAGAGTACGGCAGACGCGAAGAAATCAGCCGGCGATG<br>CCAGTGCATCAGCTGCTCAGGTCGCGGCCCTTGTGACTGATGCAACTGACTCAGCACGC<br>GCCGCCAGCACGTCCGCCGGACAGGCTGCATCGTCAGCTCAGGAAGCGTCCTCCGGCG<br>CAGAAGCGGCATCAGCAAAGGCCACTGAAGCGGAAAAAAGTGCCGCAGCCGCAGAGTCC<br>TCAAAAAACGCGGCGGCCACCAGTGCCGGTGCGGCGAAAACGTCAGAAACGAATGCTGC<br>AGCGTCACAACAATCAGCCGCCACGTCTGCCTCCACCGCGGCCACGAAAGCGTCAGAGG<br>CCGCCACTTCAGCACGAGATGCGGTGGCCTCAAAAGAGGCAGCAAAATCATCAGAAACGA<br>ACGCATCATCAAGTGCCGGTCGTGCAGCTTCCTCGGCAACGGCGGCAGAAAATTCTGCCA<br>GGGCGGCAAAAACGTCCGAGACGAATGCCAGGTCATCTGAAACAGCAGCGGAACGGAGC<br>GCCTCTGCCGCGGCAGACGCAAAAACAGCGGCGGCGGGGAGTGCGTCAACGGCATCCA<br>CGAAGGCGACAGAGGCTGCGGGAAGTGCGGTATCAGCATCGCAGAGCAAAAGTGCGGCA<br>GAAGCGGCGGCAATACGTGCAAAAAATTCGGCAAAACGTGCAGAAGATATAGCTTCAGCT<br>GTCGCGCTTGAGGATGCGGACACAACGAGAAAGGGGATAGTGCAGCTCAGCAGTGCAAC<br>CAACAGCACGTCTGAAACGCTTGCTGCAACGCCAAAGGCGGTTAAGGTGGTAATGGATGA<br>GACTAATCGTAAGGCACCTCTGGACAGTCCGGCACTGACCGGAACGCCAACAGCACCAA<br>CCGCGCTCAGGGGAACAAACAATACCCAGATTGCGAACACCGCTTTTGTACTGGCCGCGA<br>TTGCAGATGTTATCGACGCGTCACCTGACGCACTGAATACGCTGAATGAACTGGCCGCAG<br>CGCTCGGGAATGATCCAGATTTTGCTACCACCATGACTAACGCGCTTGCGGGTAAACAACC<br>GAAGAATGCGACACTGACGGCGCTGGCAGGGCTTTCCACGGCGAAAAATAAATTACCGTA<br>TTTTGCGGAAAATGATGCCGCCAGCCTGACTGAACTGACTCAGGTTGGCAGGGATATTCT<br>GGCAAAAAATTCCGTTGCAGATGTTCTTGAATACCTTGGGGCCGGTGAGAATTCGGCGGC<br>AAATGATGGCTTCGCATTCATCGGTCAGTGCCCAGACATCTTGACCCTGCGTACTATCGAG<br>CCGGAAAAAAACGGTCAGCGTATCACCTTACGTCAACATACGATTGGCACTGGCTTAGGCG<br>GTGGCGTTTTCCGTGCAGTTCTGGACGGCACTGGCTATACCGATGACGACGGTGTGGTGA<br>TCAAAACCGCTGGGGGCAGCGTTTGGCTGCGTGTCAACGCTGACAAAGTTAACCCGTTCA<br>TGTTCGGTGCAACCGGAGTAGCGGACGACACCGCCGCCCTGCAAAAAATGCTGGAATGC<br>GGTCGTGCGGCGGAACTGGGGACTAACGTATGGAAAGCAAGCAATCTGGAACTGAACAA<br>CAAATCTTGCTCTCTGTCCGGCAGTGGCCTGCACGTTTCTCGTATTGAACAGATTTCCGGT<br>GCAACCGGAGCATTGTTAACCATCACCCAAGACTGTTCGCTGATTTACCTGTCCGATTGTG<br>GCCTGTACGGCGATGGCATCACCGCAGGCACGAGCGGTGTTACTATGGAAACGGGTAATC<br>CGGGTGGCGCTCCGTCTTACCCTTTCAATACCGCTCCGGACGTTCGTCGTGACCTGTACA<br>TCTCTAACGTGCACATCACGGGCTTCGACGAGCTGGGTTTTGATTATCCGGAAACCAATTT<br>CTCTGTTTCGACGCATGGCCTCTTCATCCGTAACATCAAAAAAACGGGTGCAAAGATTGGT |

| | |
|---|---|
| | ACTACGGACTTCACTTGGACTAACCTGCAAATTGATACTTGCGGTCAGGAATGTCTGGTGC<br>TGGACGGTGCGGGTAACTGCCGTATTATTGGTGCAAAACTGATTTGGGCAGGTAGCGAAA<br>ACGAAACGCCATACTCTGGCCTGCGTATTAGCAACTCTCAAAATGTAAATATGACTGGCGTA<br>GAGTTACAAGACTGCGCGTATGATGGTTTATACATCAAGAACTCTACGGTTGCAATTTCAGG<br>CTTAAACACCAATCGCAATAGCGCATCCTCTAATCTGTCCTACCATAACATGGTATTCGAAAA<br>TTCTATTGTAACTGTTGATGGTTATGTGTGTCGTAACTACGCGGCGACTTCGCTGTACGACC<br>TGAACAGCCAAGCAGGCAACGTCCGTTGCATCGGTAGCGACAGCACCGTTTTAATCAACG<br>GCATCTACGAAAGCGAAGTCAATAGCGAGCGCCTGATGGGTGATAACAACCTGATCCAGC<br>CGTATAGTGGTGATCTGATCATTAACGGCCTGAAAAATTACTACACCTATACTGGTAGCGTAA<br>AAAACAACATTCCGACCTTCGACGGCGTTGTTACTACGGCAACCTATGTGAGCGCACCGTC<br>TATTCTGGGTCAGGGCAATATGCTCAAACTGACCCAGTCTAATAAAGACAAACTGTTATTTA<br>GCGATAAAGTTAGCCGTCATGGCTGTACCATCGGCTTAGTTCTGATTCCGTCCTTTACGGG<br>CGCGACCACTATGACGGCGTTCACGCTGGGTAGCGGTTACTCTCCATCCGGTAACTCCGC<br>CGTGATGCAGTTCATTGTTAACAGTTCCGGTGTACAAACCATTGCGATTTTATTATCCGGCG<br>ACGGTATTACCCAAACCCTGACCAGCGATCTGACCACGGAACAAGCACTGGCGAGCGGT<br>GGCGTGTATCATTTTGCAATGGGTTTTGCGCCGGGTCGTTTATGGTGGAGCATTATCGATAT<br>TAACACGGGCAGGCGTATTCGTCGCGCCTACCGTCAGCCGGATCTGCACGCGGCGTTCA<br>ACTCTATCTTCAACTCCGGCACGTCGTCTATTACCGCATTTAGCGGGCCACTGGCGGGCGA<br>CATTGCTTGCGAAGGTGCAGGTAGCCATGTATACGTTGGCGGTTTTTCGTCGGAATCTGAT<br>TACGCGGCTAGCCGTATGTATGGCCTGTTCACTCCGGTCGATCTGGACAAGCAGTATAGCT<br>TCCGTACCCTGAACGGTAACATTAACTATATTGTGAGGCTTGCATAATGGCATTCAGAATGA<br>GTGAACAACCACGGACCATAAAAATTTATAATCTGCTGGCCGGAACTAATGAATTTATTGGT<br>GAAGGTGACGCATATATTCCGCCTCATACCGGTCTGCCTGCAAACAGTACCGATATTGCAC<br>CGCCAGATATTCCGGCTGGCTTTGTGGCTGTTTTCAACAGTGATGAGGCATCGTGGCATCT<br>CGTTGAAGACCATCGGGGTAAAACCGTCTATGACGTGGCTTCCGGCGACGCGTTATTTATT<br>TCTGAACTCGGTCCGTTACCGGAAAATTTTACCTGGTTATCGCCGGGAGGGGAATATCAGA<br>AGTGGAACGGCACAGCCTGGGTGAAGGATACGGAAGCAGAAAAACTGTTCCGGATCCGG<br>GAGGCGGAAGAAACAAAAAAAAGCCTGATGCAGGTAGCCAGTGAGCATATTGCGCCGCTT<br>CAGGATGCTGCAGATCTGGAAATTGCAACGAAGGAAGAAACCTCGTTGCTGGAAGCCTGG<br>AAGAAGTATCGGGTGTTGCTGAACCGTGTTGATACATCAACTGCACCTGATATTGAGTGGC<br>CTGCTGTCCCTGTTATGGAGTAATGACGCATCCTCACGATAATATCCGGGTAGGACGAACA<br>ATAAGGCCGCAAATCGCGGCCTTTTTTATTGATAACAAAAGGACAGTTTTCCCTTTGATATGT<br>AACGGTGAACAGTTGTTCTACTTTTGTTTGTTAGTCTTGATGCTTCACTGATAGATACAAGA<br>GCCATAAGAACC |
| p3606 | TCAGATCCTTCCGTATTTAGCCAGTATGTTCTCTAGTGTGGTTCGTTGTTTTTGCGTGAGCC<br>ATGAGAACGAACCATTGAGATCATACTTACTTTGCATGTCACTCAAAAATTTTGCCTCAAAAC<br>TGGTGAGCTGAATTTTTGCAGTTAAAGCATCGTGTAGTGTTTTTCTTAGTCCGTTACGTAGG<br>TAGGAATCTGATGTAATGGTTGTTGGTATTTTGTCACCATTCATTTTTATCTGGTTGTTCTCAA<br>GTTCGGTTACGAGATCCATTTGTCTATCTAGTTCAACTTGGAAAATCAACGTATCAGTCGGG<br>CGGCCTCGCTTATCAACCACCAATTTCATATTGCTGTAAGTGTTTAAATCTTTACTTATTGGT<br>TTCAAAACCCATTGGTTAAGCCTTTTAAACTCATGGTAGTTATTTTCAAGCATTAACATGAAC<br>TTAAATTCATCAAGGCTAATCTCTATATTTGCCTTGTGAGTTTTCTTTTGTGTTAGTTCTTTTA<br>ATAACCACTCATAAATCCTCATAGAGTATTTGTTTTCAAAAGACTTAACATGTTCCAGATTATA<br>TTTTATGAATTTTTTTAACTGGAAAAGATAAGGCAATATCTCTTCACTAAAAACTAATTCTAATT<br>TTTCGCTTGAGAACTTGGCATAGTTTGTCCACTGGAAAATCTCAAAGCCTTTAACCAAAGG<br>ATTCCTGATTTCCACAGTTCTCGTCATCAGCTCTCTGGTTGCTTTAGCTAATACACCATAAG<br>CATTTTCCCTACTGATGTTCATCATCTGAGCGTATTGGTTATAAGTGAACGATACCGTCCGTT<br>CTTTCCTTGTAGGGTTTTCAATCGTGGGGTTGAGTAGTGCCACACAGCATAAAATTAGCTTG |

| | |
|---|---|
| | GTTTCATGCTCCGTTAAGTCATAGCGACTAATCGCTAGTTCATTTGCTTTGAAAACAACTAAT<br>TCAGACATACATCTCAATTGGTCTAGGTGATTTTAATCACTATACCAATTGAGATGGGCTAGT<br>CAATGATAATTACTAGTCCTTTTCCTTTGAGTTGTGGGTATCTGTAAATTCTGCTAGACCTTT<br>GCTGGAAAACTTGTAAATTCTGCTAGACCCTCTGTAAATTCCGCTAGACCTTTGTGTGTTTT<br>TTTTGTTTATATTCAAGTGGTTATAATTTATAGAATAAAGAAAGAATAAAAAAAGATAAAAAGAA<br>TAGATCCCAGCCCTGTGTATAACTCACTACTTTAGTCAGTTCCGCAGTATTACAAAAGGATG<br>TCGCAAACGCTGTTTGCTCCTCTACAAAACAGACCTTAAAACCCTAAAGGCTTAAGTAGCA<br>CCCTCGCAAGCTCGGTTGCGGCCGCAATCGGGCAAATCGCTGAATATTCCTTTTGTCTCC<br>GACCATCAGGCACCTGAGTCGCTGTCTTTTTCGTGACATTCAGTTCGCTGCGCTCACGGC<br>TCTGGCAGTGAATGGGGGTAAATGGCACTACAGGCGCCTTTTATGGATTCATGCAAGGAAA<br>CTACCCATAATACAAGAAAAGCCCGTCACGGGCTTCTCAGGGCGTTTTATGGCGGGTCTG<br>CTATGTGGTGCTATCTGACTTTTTGCTGTTCAGCAGTTCCTGCCCTCTGATTTTCCAGTCTG<br>ACCACTTCGGATTATCCCGTGACAGGTCATTCAGACTGGCTAATGCACCCAGTAAGGCAGC<br>GGTATCATCAACGGGGTCTGACGCTCAGTGGAACGAAAACTCACGTTAAGGGATTTTGGT<br>CATGAGATTATCAAAAAGGATCTTCACCTAGATCCTTTTAAATTAAAAATGAAGTTTTAAATCA<br>ATCTAAAGTATATATGAGTAAACTTGGTCTGACAGTTACGTTTCCACAACCAATTAACCAATT<br>CTGATTTAGAAAAACTCATCGAGCATCAAATGAAACTGCAATTTATTCATATCAGGATTATCAA<br>TACCATATTTTTGAAAAAGCCGTTTCTGTAATGAAGGAGAAAACTCACCGAGGCAGTTCCAT<br>AGGATGGCAAGATCCTGGTATCGGTCTGCGATTCCGACTCGTCCAACATCAATACAACCTA<br>TTAATTTCCCCTCGTCAAAAATAAGGTTATCAAGTGAGAAATCACCATGAGTGACGACTGAA<br>TCCGGTGAGAATGGCAAAAGCTTATGCATTTCTTTCCAGACTTGTTCAACAGGCCAGCCAT<br>TACGCTCGTCATCAAAATCACTCGCATCAACCAAACCGTTATTCATTCGTGATTGCGCCTGA<br>GCGAGACGAAATACGCGATCGCTGTTAAAAGGACAATTACAAACAGGAATCGAATGCAACC<br>GGCGCAGGAACACTGCCAGCGCATCAACAATATTTTCACCTGAATCAGGATATTCTTCTAAT<br>ACCTGGAATGCTGTTTTCCCGGGGATCGCAGTGGTGAGTAACCATGCATCATCAGGAGTA<br>CGGATAAAATGCTTGATGGTCGGAAGAGGCATAAATTCCGTCAGCCAGTTTAGTCTGACCA<br>TCTCATCTGTAACATCATTGGCAACGCTACCTTTGCCATGTTTCAGAAACAACTCTGGCGCA<br>TCGGGCTTCCCATACAATCGATAGATTGTCGCACCTGATTGCCCGACATTATCGCGAGCCC<br>ATTTATACCCATATAAATCAGCATCCATGTTGGAATTTAATCGCGGCCTCGAGCAAGACGTTT<br>CCCGTTGAATATGGCTCATAACACCCCTTGTATTACTGTTTATGTAAGCAGACAGTTTTATTG<br>TTCATGATGATATATTTTTATCTTGTGCAATGTAACATCAGAGATTTTGAGACACAACGTGGC<br>TTTCCCTGCAGGATTTCGGAGGCCTGCGTTATCCCCTGATTCTGTGGATAACCGTATTACC<br>GCCTTTGAGTGAGCTGATACCGCTCGCCGCAGCCGAACGCCGACTAGTGGATTTTACGGC<br>TAGCTCAGTCCTAGGTACAATGCTAGCGAATTCATTAAAGAGGAGAAAGGTACCCATGGCA<br>CGTACCCCGAGCCGTAGCAGCATTGGTAGCCTGCGTAGTCCGCATACCCATAAAGCAATTC<br>TGACCAGCACCATTGAAATCCTGAAAGAATGTGGTTATAGCGGTCTGAGCATTGAAAGCGT<br>TGCACGTCGTGCCGGTGCAAGCAAACCGACCATTTATCGTTGGTGGACCAATAAAGCAGC<br>ACTGATTGCCGAAGTGTATGAAAATGAAAGCGAACAGGTGCGTAAATTTCCGGATCTGGGT<br>AGCTTTAAAGCCGATCTGGATTTTCTGCTGCGTAATCTGTGGAAAGTTTGGCGTGAAACCA<br>TTTGTGGTGAAGCATTTCGTTGTGTTATTGCAGAAGCACAGCTGGACCCTGCAACCCTGAC<br>CCAGCTGAAAGATCAGTTTATGGAACGTCGTCGTGAGATGCCGAAAAAACTGGTTGAAAAT<br>GCCATTAGCAATGGTGAACTGCCGAAAGATACCAATCGTGAACTGCTGCTGGATATGATTTT<br>TGGTTTTTGTTGGTATCGCCTGCTGACCGAACAGCTGACCGTTGAACAGGATATTGAAGAA<br>TTTACCTTCCTGCTAATTAATGGTGTTTGTCCGGGTACACAGCGTTAACTAGGGCCCATACC<br>CCCAATTATTGAAGGCCGCTAACGCGGCCTTTTTTTGTTTCTGGTCTGCCCGACGTACGGT<br>GAATCTGATTCGTTACCAATTGACATGATACGAAACGTACCGTATCGTTAAGGTTACTAGATT<br>AAAGAGGAGAAATACTAGATGGCAGTAAAGATTTCAGGAGTCCTGAAAGACGGCACAGGA<br>AAACCGGTACAGAACTGCACCATTCAGCTGAAAGCCAGACGTAACAGCACCACGGTGGTG |

| | |
|---|---|
| | GTGAACACGGTGGGCTCAGAGAATCCGGATGAAGCCGGGCGTTACAGCATGGATGTGGA<br>GTACGGTCAGTACAGTGTCATCCTGCAGGTTGACGGTTTTCCACCATCGCACGCCGGGAC<br>CATCACCGTGTATGAAGATTCACAACCGGGGACGCTGAATGATTTTCTCTGTGCCATGACG<br>GAGGATGATGCCCGGCCGGAGGTGCTGCGTCGTCTTGAACTGATGGTGGAAGAGGTGGC<br>GCGTAACGCGTCCGTGGTGGCACAGAGTACGGCAGACGCGAAGAAATCAGCCGGCGATG<br>CCAGTGCATCAGCTGCTCAGGTCGCGGCCCTTGTGACTGATGCAACTGACTCAGCACGC<br>GCCGCCAGCACGTCCGCCGGACAGGCTGCATCGTCAGCTCAGGAAGCGTCCTCCGGCG<br>CAGAAGCGGCATCAGCAAAGGCCACTGAAGCGGAAAAAAGTGCCGCAGCCGCAGAGTCC<br>TCAAAAAACGCGGCGGCCACCAGTGCCGGTGCGGCGAAAACGTCAGAAACGAATGCTGC<br>AGCGTCACAACAATCAGCCGCCACGTCTGCCTCCACCGCGGCCACGAAAGCGTCAGAGG<br>CCGCCACTTCAGCACGAGATGCGGTGGCCTCAAAAGAGGCAGCAAAATCATCAGAAACGA<br>ACGCATCATCAAGTGCCGGTCGTGCAGCTTCCTCGGCAACGGCGGCAGAAAATTCTGCCA<br>GGGCGGCAAAAACGTCCGAGACGAATGCCAGGTCATCTGAAACAGCAGCGGAACGGAGC<br>GCCTCTGCCGCGGCAGACGCAAAAACAGCGGCGGCGGGGAGTGCGTCAACGGCATCCA<br>CGAAGGCGACAGAGGCTGCGGGAAGTGCGGTATCAGCATCGCAGAGCAAAAGTGCGGCA<br>GAAGCGGCGGCAATACGTGCAAAAAATTCGGCAAAACGTGCAGAAGATATAGCTTCAGCT<br>GTCGCGCTTGAGGATGCGGACACAACGAGAAAGGGGATAGTGCAGCTCAGCAGTGCAAC<br>CAACAGCACGTCTGAAACGCTTGCTGCAACGCCAAAGGCGGTTAAGGTGGTAATGGATGA<br>GACTAATCGTAAGGCACCTCTGGACAGTCCGGCACTGACCGGAACGCCAACAGCACCAA<br>CCGCGCTCAGGGGAACAAACAATACCCAGATTGCGAACACCGCTTTTGTACTGGCCGCGA<br>TTGCAGATGTTATCGACGCGTCACCTGACGCACTGAATACGCTGAATGAACTGGCCGCAG<br>CGCTCGGGAATGATCCAGATTTTGCTACCACCATGACTAACGCGCTTGCGGGTAAACAACC<br>GAAGAATGCGACACTGACGGCGCTGGCAGGGCTTTCCACGGCGAAAAATAAATTACCGTA<br>TTTTGCGGAAAATGATGCCGCCAGCCTGACTGAACTGACTCAGGTTGGCAGGGATATTCT<br>GGCAAAAAATTCCGTTGCAGATGTTCTTGAATACCTTGGGGCCGGTGAGAATTCGGCGGC<br>AAATGATGGCTTCGCATTCATCGGTCAGTGCCCAGACATCTTGACCCTGCGTACTATCGAG<br>CCGGAAAAAAACGGTCAGCGTATCACCTTACGTCAACATACGATTGGCACTGGCTTAGGCG<br>GTGGCGTTTTCCGTGCAGTTCTGGACGGCACTGGCTATACCGATGACGACGGTGTGGTGA<br>TCAAAACCGCTGGGGGCAGCGTTTGGCTGCGTGTCAACGCTGACAAAGTTAACCCGTTCA<br>TGTTCGGTGCAACCGGAGTAGCGGACGACACCGCCGCCCTGCAAAAAATGCTGGAATGC<br>GGTCGTGCGGCGGAACTGGGGACTAACGTATGGAAAGCAAGCAATCTGGAACTGAACAA<br>CAAATCTTGCTCTCTGTCCGGCAGTGGCCTGCACGTTTCTCGTATTGAACAGATTTCCGGT<br>GCAACCGGAGCATTGTTAACCATCACCCAAGACTGTTCGCTGATTTACCTGTCCGATTGTG<br>GCCTGTACGGCGATGGCATCACCGCAGGCACGAGCGGTGTTACTATGGAAACGGGTAATC<br>CGGGTGGCGCTCCGTCTTACCCTTTCAATACCGCTCCGGACGTTCGTCGTGACCTGTACA<br>TCTCTAACGTGCACATCACGGGCTTCGACGAGCTGGGTTTTGATTATCCGGAAACCAATTT<br>CTCTGTTTCGACGCATGGCCTCTTCATCCGTAACATCAAAAAAACGGGTGCAAAGATTGGT<br>ACTACGGACTTCACTTGGACTAACCTGCAAATTGATACTTGCGGTCAGGAATGTCTGGTGC<br>TGGACGGTGCGGGTAACTGCCGTATTATTGGTGCAAAACTGATTTGGGCAGGTAGCGAAA<br>ACGAAACGCCATACTCTGGCCTGCGTATTAGCAACTCTCAAAATGTAAATATGACTGGCGTA<br>GAGTTACAAGACTGCGCGTATGATGGTTTATACATCAAGAACTCTACGGTTGCAATTTCAGG<br>CTTAAACACCAATCGCAATAGCGCATCCTCTAATCTGTCCTACCATAACATGGTATTCGAAAA<br>TTCTATTGTAACTGTTGATGGTTATGTGTGTCGTAACTACGCGGCGACTTCGCTGTACGACC<br>TGAACAGCCAAGCAGGCAACGTCCGTTGCATCGGTAGCGACAGCACCGTTTTAATCAACG<br>GCATCTACGAAAGCGAAGTCAATAGCGAGCGCCTGATGGGTGATAACAACCTGATCCAGC<br>CGTATAGTGGTGATCTGATCATTAACGGCCTGAAAAATTACTACACCTATACTGGTAGCGTAA<br>AAAACAACATTCCGACCTTCGACGGCGTTGTTACTACGGCAACCTATGTGAGCGCACCGTC<br>TATTCTGGGTCAGGGCAATATGCTCAAACTGACCCAGTCTAATAAAGACAAACTGTTATTTA |

| | |
|---|---|
| | GCGATAAAGTTAGCCGTCATGGCTGTACCATCGGCTTAGTTCTGATTCCGTCCTTTACGGG<br>CGCGACCACTATGACGGCGTTCACGCTGGGTAGCGGTTACTCTCCATCCGGTAACTCCGC<br>CGTGATGCAGTTCATTGTTAACAGTTCCGGTGTACAAACCATTGCGATTTTATTATCCGGCG<br>ACGGTATTACCCAAACCCTGACCAGCGATCTGACCACGGAACAAGCACTGGCGAGCGGT<br>GGCGTGTATCATTTTGCAATGGGTTTTGCGCCGGGTCGTTTATGGTGGAGCATTATCGATAT<br>TAACACGGGCAGGCGTATTCGTCGCGCCTACCGTCAGCCGGATCTGCACGCGGCGTTCA<br>ACTCTATCTTCAACTCCGGCACGTCGTCTATTACCGCATTTAGCGGGCCACTGGCGGGCGA<br>CATTGCTTGCGAAGGTGCAGGTAGCCATGTATACGTTGGCGGTTTTTCGTCGGAATCTGAT<br>TACGCGGCTAGCCGTATGTATGGCCTGTTCACTCCGGTCGATCTGGACAAGCAGTATAGCT<br>TCCGTACCCTGAACGGTAACATTAACTATATTGTGAGGCTTGCATAATGGCATTCAGAATGA<br>GTGAACAACCACGGACCATAAAAATTTATAATCTGCTGGCCGGAACTAATGAATTTATTGGT<br>GAAGGTGACGCATATATTCCGCCTCATACCGGTCTGCCTGCAAACAGTACCGATATTGCAC<br>CGCCAGATATTCCGGCTGGCTTTGTGGCTGTTTTCAACAGTGATGAGGCATCGTGGCATCT<br>CGTTGAAGACCATCGGGGTAAAACCGTCTATGACGTGGCTTCCGGCGACGCGTTATTTATT<br>TCTGAACTCGGTCCGTTACCGGAAAATTTTACCTGGTTATCGCCGGGAGGGGAATATCAGA<br>AGTGGAACGGCACAGCCTGGGTGAAGGATACGGAAGCAGAAAAACTGTTCCGGATCCGG<br>GAGGCGGAAGAAACAAAAAAAAGCCTGATGCAGGTAGCCAGTGAGCATATTGCGCCGCTT<br>CAGGATGCTGCAGATCTGGAAATTGCAACGAAGGAAGAAACCTCGTTGCTGGAAGCCTGG<br>AAGAAGTATCGGGTGTTGCTGAACCGTGTTGATACATCAACTGCACCTGATATTGAGTGGC<br>CTGCTGTCCCTGTTATGGAGTAATGACGCATCCTCACGATAATATCCGGGTAGGACGAACA<br>ATAAGGCCGCAAATCGCGGCCTTTTTTATTGATAACAAAAGGACAGTTTTCCCTTTGATATGT<br>AACGGTGAACAGTTGTTCTACTTTTGTTTGTTAGTCTTGATGCTTCACTGATAGATACAAGA<br>GCCATAAGAACC |
| p766 | TACGGTTATCCACAGAATCAGGGGATAACGCAGGCCTCCGAAATCCTGCAGGGAAAGCCA<br>CGTTGTGTCTCAAAATCTCTGATGTTACATTGCACAAGATAAAAATATATCATCATGAACAATA<br>AAACTGTCTGCTTACATAAACAGTAATACAAGGGGTGTTATGAGCCATATTCAACGGGAAAC<br>GTCTTGCTCGAGGCCGCGATTAAATTCCAACATGGATGCTGATTTATATGGGTATAAATGGG<br>CTCGCGATAATGTCGGGCAATCAGGTGCGACAATCTATCGATTGTATGGGAAGCCCGATGC<br>GCCAGAGTTGTTTCTGAAACATGGCAAAGGTAGCGTTGCCAATGATGTTACAGATGAGATG<br>GTCAGACTAAACTGGCTGACGGAATTTATGCCTCTTCCGACCATCAAGCATTTTATCCGTAC<br>TCCTGATGATGCATGGTTACTCACCACTGCGATCCCCGGGAAAACAGCATTCCAGGTATTA<br>GAAGAATATCCTGATTCAGGTGAAAATATTGTTGATGCGCTGGCAGTGTTCCTGCGCCGGT<br>TGCATTCGATTCCTGTTTGTAATTGTCCTTTTAACAGCGATCGCGTATTTCGTCTCGCTCAG<br>GCGCAATCACGAATGAATAACGGTTTGGTTGATGCGAGTGATTTTGATGACGAGCGTAATG<br>GCTGGCCTGTTGAACAAGTCTGGAAAGAAATGCATAAGCTTTTGCCATTCTCACCGGATTC<br>AGTCGTCACTCATGGTGATTTCTCACTTGATAACCTTATTTTTGACGAGGGGAAATTAATAG<br>GTTGTATTGATGTTGGACGAGTCGGAATCGCAGACCGATACCAGGATCTTGCCATCCTATG<br>GAACTGCCTCGGTGAGTTTTCTCCTTCATTACAGAAACGGCTTTTTCAAAAATATGGTATTG<br>ATAATCCTGATATGAATAAATTGCAGTTTCATTTGATGCTCGATGAGTTTTTCTAAATCAGAAT<br>TGGTTAATTGGTTGTGGAAACGTAACTGTCAGACCAAGTTTACTCATATATACTTTAGATTGA<br>TTTAAAACTTCATTTTTAATTTAAAAGGATCTAGGTGAAGATCCTTTTTGATAATCTCATGACC<br>AAAATCCCTTAACGTGAGTTTTCGTTCCACTGAGCGTCAGACCCCGTTGACAGTAAGACGG<br>GTAAGCCTGTTGATGATACCGCTGCCTTACTGGGTGCATTAGCCAGTCTGAATGACCTGTC<br>ACGGGATAATCCGAAGTGGTCAGACTGGAAAATCAGAGGGCAGGAACTGCTGAACAGCAA<br>AAAGTCAGATAGCACCACATAGCAGACCCGCCATAAAACGCCCTGAGAAGCCCGTGACGG<br>GCTTTTCTTGTATTATGGGTAGTTTCCTTGCATGAATCCATAAAAGGCGCCTGTAGTGCCATT<br>TACCCCCATTCACTGCCAGAGCCGTGAGCGCAGCGAACTGAATGTCACGAAAAAGACAGC<br>GACTCAGGTGCCTGATGGTCGGAGACAAAAGGAATATTCAGCGATTTGCCCGATTGCGGC |

| | |
|---|---|
| | CGCAACCGAGCTTGCGAGGGTGCTACTTAAGCCTTTAGGGTTTTAAGGTCTGTTTTGTAGA<br>GGAGCAAACAGCGTTTGCGACATCCTTTTGTAATACTGCGGAACTGACTAAAGTAGTGAGT<br>TATACACAGGGCTGGGATCTATTCTTTTTATCTTTTTTTATTCTTTCTTTATTCTATAAATTATAA<br>CCACTTGAATATAAACAAAAAAAACACACAAAGGTCTAGCGGAATTTACAGAGGGTCTAGCA<br>GAATTTACAAGTTTTCCAGCAAAGGTCTAGCAGAATTTACAGATACCCACAACTCAAAGGAA<br>AAGGACTAGTAATTATCATTGACTAGCCCATCTCAATTGGTATAGTGATTAAAATCACCTAGA<br>CCAATTGAGATGTATGTCTGAATTAGTTGTTTTCAAAGCAAATGAACTAGCGATTAGTCGCTA<br>TGACTTAACGGAGCATGAAACCAAGCTAATTTTATGCTGTGTGGCACTACTCAACCCCACG<br>ATTGAAAACCCTACAAGGAAAGAACGGACGGTATCGTTCACTTATAACCAATACGCTCAGAT<br>GATGAACATCAGTAGGGAAAATGCTTATGGTGTATTAGCTAAAGCAACCAGAGAGCTGATGA<br>CGAGAACTGTGGAAATCAGGAATCCTTTGGTTAAAGGCTTTGAGATTTTCCAGTGGACAAA<br>CTATGCCAAGTTCTCAAGCGAAAAATTAGAATTAGTTTTTAGTGAAGAGATATTGCCTTATCT<br>TTTCCAGTTAAAAAAATTCATAAAATATAATCTGGAACATGTTAAGTCTTTTGAAAACAAATAC<br>TCTATGAGGATTTATGAGTGGTTATTAAAAGAACTAACACAAAAGAAAACTCACAAGGCAAA<br>TATAGAGATTAGCCTTGATGAATTTAAGTTCATGTTAATGCTTGAAAATAACTACCATGAGTTT<br>AAAAGGCTTAACCAATGGGTTTTGAAACCAATAAGTAAAGATTTAAACACTTACAGCAATATG<br>AAATTGGTGGTTGATAAGCGAGGCCGCCCGACTGATACGTTGATTTTCCAAGTTGAACTAG<br>ATAGACAAATGGATCTCGTAACCGAACTTGAGAACAACCAGATAAAAATGAATGGTGACAAA<br>ATACCAACAACCATTACATCAGATTCCTACCTACGTAACGGACTAAGAAAAACACTACACGA<br>TGCTTTAACTGCAAAAATTCAGCTCACCAGTTTTGAGGCAAAATTTTTGAGTGACATGCAAA<br>GTAAGTATGATCTCAATGGTTCGTTCTCATGGCTCACGCAAAAACAACGAACCACACTAGA<br>GAACATACTGGCTAAATACGGAAGGATCTGAGGTTCTTATGGCTCTTGTATCTATCAGTGAA<br>GCATCAAGACTAACAAACAAAAGTAGAACAACTGTTCACCGTTACATATCAAAGGGAAAACT<br>GTCCATATGCACAGATGAAAACGGTGTAAAAAAGATAGATACATCAGAGCTTTTACGAGTTT<br>TTGGTGCATTTAAAGCTGTTCACCATGAACAGATCGACAATGTAACAGATGAACAGCATGTA<br>ACACCTAATAGAACAGGTGAAACCAGTAAAACAAAGCAACTAGAACATGAAATTGAACACCT<br>GAGACAACTTGTTACAGCTCAACAGTCACACATAGACAGCCTGAAACAGGCGATGCTGCTT<br>ATCGAATCAAAGCTGCCGACAACACGGGAGCCAGTGACGCCTCCCGTGGGGAAAAAATC<br>ATGGCAATTCTGGAAGAAATAGCGCTTTCATCCGGCAAACCTGAAACCGGATCTGCGATTC<br>TGATAACAAACTAGCAACACCAGAACAGCCCGTTTGCGGGCAGTAAAACCCGTTTTGCTGA<br>TTTTTCACATGTTACCTTTCTTTACGGCTAGCTCAGTCCTAGGTACAATGCTAGCGTTTTCAT<br>TAAAGAGGAGAAAGGAAGCCATGGAGGACGTTATCAAAGAGTTCATGCGTTTCAAAGTTCG<br>TATGGAAGGTAGCGTTAACGGTCACGAGTTCGAAATCGAAGGTGAGGGTGAAGGTCGTCC<br>GTACGAAGGTACTCAAACCGCTAAACTGAAAGTTACCAAAGGTGGCCCTCTGCCGTTCGC<br>TTGGGACATCCTGTCCCCGCAGTTCCAGTACGGTAGCAAGGCTTACGTTAAACATCCGGCT<br>GACATTCCGGACTACTTAAAGCTGTCCTTTCCGGAAGGTTTCAAATGGGAGCGTGTTATGA<br>ACTTCGAGGACGGCGGTGTTGTTACCGTTACGCAGGACTCCTCCCTGCAAGACGGTGAGT<br>TCATCTACAAAGTTAAACTGCGTGGTACGAACTTCCCGTCCGACGGCCCGGTTATGCAGAA<br>AAAAACTATGGGTTGGGAGGCTTCCACCGAACGTATGTATCCGGAGGACGGTGCGCTGAA<br>AGGTGAAATCAAAATGCGTCTGAAACTGAAAGACGGCGGTCACTACGACGCGGAAGTTAA<br>AACTACCTACATGGCTAAAAAACCTGTTCAACTGCCGGGCGCTTACAAAACCGACATCAAA<br>CTGGACATCACCTCCCACAACGAGGACTATACCATCGTTGAACAGTACGAACGCGCGGAA<br>GGTCGTCACTCCACTGGTGCTTAAGTTTAGCCGAACGCCCGCAAAAAACCCCGCTTCGGC<br>GGGGTTTTTTCGC |
| p767 | TACGGTTATCCACAGAATCAGGGGATAACGCAGGCCTCCGAAATCCTGCAGGGAAAGCCA<br>CGTTGTGTCTCAAAATCTCTGATGTTACATTGCACAAGATAAAAATATATCATCATGAACAATA<br>AAACTGTCTGCTTACATAAACAGTAATACAAGGGGTGTTATGAGCCATATTCAACGGGAAAC<br>GTCTTGCTCGAGGCCGCGATTAAATTCCAACATGGATGCTGATTTATATGGGTATAAATGGG |

| | |
|---|---|
| | CTCGCGATAATGTCGGGCAATCAGGTGCGACAATCTATCGATTGTATGGGAAGCCCGATGC<br>GCCAGAGTTGTTTCTGAAACATGGCAAAGGTAGCGTTGCCAATGATGTTACAGATGAGATG<br>GTCAGACTAAACTGGCTGACGGAATTTATGCCTCTTCCGACCATCAAGCATTTTATCCGTAC<br>TCCTGATGATGCATGGTTACTCACCACTGCGATCCCCGGGAAAACAGCATTCCAGGTATTA<br>GAAGAATATCCTGATTCAGGTGAAAATATTGTTGATGCGCTGGCAGTGTTCCTGCGCCGGT<br>TGCATTCGATTCCTGTTTGTAATTGTCCTTTTAACAGCGATCGCGTATTTCGTCTCGCTCAG<br>GCGCAATCACGAATGAATAACGGTTTGGTTGATGCGAGTGATTTTGATGACGAGCGTAATG<br>GCTGGCCTGTTGAACAAGTCTGGAAAGAAATGCATAAGCTTTTGCCATTCTCACCGGATTC<br>AGTCGTCACTCATGGTGATTTCTCACTTGATAACCTTATTTTTGACGAGGGGAAATTAATAG<br>GTTGTATTGATGTTGGACGAGTCGGAATCGCAGACCGATACCAGGATCTTGCCATCCTATG<br>GAACTGCCTCGGTGAGTTTTCTCCTTCATTACAGAAACGGCTTTTTCAAAAATATGGTATTG<br>ATAATCCTGATATGAATAAATTGCAGTTTCATTTGATGCTCGATGAGTTTTTCTAAATCAGAAT<br>TGGTTAATTGGTTGTGGAAACGTAACTGTCAGACCAAGTTTACTCATATATACTTTAGATTGA<br>TTTAAAACTTCATTTTTAATTTAAAAGGATCTAGGTGAAGATCCTTTTTGATAATCTCATGACC<br>AAAATCCCTTAACGTGAGTTTTCGTTCCACTGAGCGTCAGACCCCGTTGACAGTAAGACGG<br>GTAAGCCTGTTGATGATACCGCTGCCTTACTGGGTGCATTAGCCAGTCTGAATGACCTGTC<br>ACGGGATAATCCGAAGTGGTCAGACTGGAAAATCAGAGGGCAGGAACTGCTGAACAGCAA<br>AAAGTCAGATAGCACCACATAGCAGACCCGCCATAAAACGCCCTGAGAAGCCCGTGACGG<br>GCTTTTCTTGTATTATGGGTAGTTTCCTTGCATGAATCCATAAAAGGCGCCTGTAGTGCCATT<br>TACCCCCATTCACTGCCAGAGCCGTGAGCGCAGCGAACTGAATGTCACGAAAAAGACAGC<br>GACTCAGGTGCCTGATGGTCGGAGACAAAAGGAATATTCAGCGATTTGCCCGATTGCGGC<br>CGCAACCGAGCTTGCGAGGGTGCTACTTAAGCCTTTAGGGTTTTAAGGTCTGTTTTGTAGA<br>GGAGCAAACAGCGTTTGCGACATCCTTTTGTAATACTGCGGAACTGACTAAAGTAGTGAGT<br>TATACACAGGGCTGGGATCTATTCTTTTTATCTTTTTTTATTCTTTCTTTATTCTATAAATTATAA<br>CCACTTGAATATAAACAAAAAAAACACACAAAGGTCTAGCGGAATTTACAGAGGGTCTAGCA<br>GAATTTACAAGTTTTCCAGCAAAGGTCTAGCAGAATTTACAGATACCCACAACTCAAAGGAA<br>AAGGACTAGTAATTATCATTGACTAGCCCATCTCAATTGGTATAGTGATTAAAATCACCTAGA<br>CCAATTGAGATGTATGTCTGAATTAGTTGTTTTCAAAGCAAATGAACTAGCGATTAGTCGCTA<br>TGACTTAACGGAGCATGAAACCAAGCTAATTTTATGCTGTGTGGCACTACTCAACCCCACG<br>ATTGAAAACCCTACAAGGAAAGAACGGACGGTATCGTTCACTTATAACCAATACGCTCAGAT<br>GATGAACATCAGTAGGGAAAATGCTTATGGTGTATTAGCTAAAGCAACCAGAGAGCTGATGA<br>CGAGAACTGTGGAAATCAGGAATCCTTTGGTTAAAGGCTTTGAGATTTTCCAGTGGACAAA<br>CTATGCCAAGTTCTCAAGCGAAAAATTAGAATTAGTTTTTAGTGAAGAGATATTGCCTTATCT<br>TTTCCAGTTAAAAAAATTCATAAAATATAATCTGGAACATGTTAAGTCTTTTGAAAACAAATAC<br>TCTATGAGGATTTATGAGTGGTTATTAAAAGAACTAACACAAAAGAAAACTCACAAGGCAAA<br>TATAGAGATTAGCCTTGATGAATTTAAGTTCATGTTAATGCTTGAAAATAACTACCATGAGTTT<br>AAAAGGCTTAACCAATGGGTTTTGAAACCAATAAGTAAAGATTTAAACACTTACAGCAATATG<br>AAATTGGTGGTTGATAAGCGAGGCCGCCCGACTGATACGTTGATTTTCCAAGTTGAACTAG<br>ATAGACAAATGGATCTCGTAACCGAACTTGAGAACAACCAGATAAAAATGAATGGTGACAAA<br>ATACCAACAACCATTACATCAGATTCCTACCTACGTAACGGACTAAGAAAAACACTACACGA<br>TGCTTTAACTGCAAAAATTCAGCTCACCAGTTTTGAGGCAAAATTTTTGAGTGACATGCAAA<br>GTAAGTATGATCTCAATGGTTCGTTCTCATGGCTCACGCAAAAACAACGAACCACACTAGA<br>GAACATACTGGCTAAATACGGAAGGATCTGAGGTTCTTATGGCTCTTGTATCTATCAGTGAA<br>GCATCAAGACTAACAAACAAAAGTAGAACAACTGTTCACCGTTACATATCAAAGGGAAAACT<br>GTCCATATGCACAGATGAAAACGGTGTAAAAAAGATAGATACATCAGAGCTTTTACGAGTTT<br>TTGGTGCATTTAAAGCTGTTCACCATGAACAGATCGACAATGTAACAGATGAACAGCATGTA<br>ACACCTAATAGAACAGGTGAAACCAGTAAAACAAAGCAACTAGAACATGAAATTGAACACCT<br>GAGACAACTTGTTACAGCTCAACAGTCACACATAGACAGCCTGAAACAGGCGATGCTGCTT |

| | |
|---|---|
| | ATCGAATCAAAGCTGCCGACAACACGGGAGCCAGTGACGCCTCCCGTGGGGAAAAAATC<br>ATGGCAATTCTGGAAGAAATAGCGCTTTCATCCGGCAAACCTGAAACCGGATCTGCGATTC<br>TGATAACAAACTAGCAACACCAGAACAGCCCGTTTGCGGGCAGTAAAACCCGTTTTGCATT<br>CCGGAACGTTCCAGCGCTGCTTTACGGCTAGCTCAGTCCTAGGTACAATGCTAGCGTTTTC<br>ATTAAAGAGGAGAAAGGAAGCCATGGAGGACGTTATCAAAGAGTTCATGCGTTTCAAAGTT<br>CGTATGGAAGGTAGCGTTAACGGTCACGAGTTCGAAATCGAAGGTGAGGGTGAAGGTCGT<br>CCGTACGAAGGTACTCAAACCGCTAAACTGAAAGTTACCAAAGGTGGCCCTCTGCCGTTC<br>GCTTGGGACATCCTGTCCCCGCAGTTCCAGTACGGTAGCAAGGCTTACGTTAAACATCCG<br>GCTGACATTCCGGACTACTTAAAGCTGTCCTTTCCGGAAGGTTTCAAATGGGAGCGTGTTA<br>TGAACTTCGAGGACGGCGGTGTTGTTACCGTTACGCAGGACTCCTCCCTGCAAGACGGTG<br>AGTTCATCTACAAAGTTAAACTGCGTGGTACGAACTTCCCGTCCGACGGCCCGGTTATGCA<br>GAAAAAAACTATGGGTTGGGAGGCTTCCACCGAACGTATGTATCCGGAGGACGGTGCGCT<br>GAAAGGTGAAATCAAAATGCGTCTGAAACTGAAAGACGGCGGTCACTACGACGCGGAAGT<br>TAAAACTACCTACATGGCTAAAAAACCTGTTCAACTGCCGGGCGCTTACAAAACCGACATC<br>AAACTGGACATCACCTCCCACAACGAGGACTATACCATCGTTGAACAGTACGAACGCGCG<br>GAAGGTCGTCACTCCACTGGTGCTTAAGTTTAGCCGAACGCCCGCAAAAAACCCCGCTTC<br>GGCGGGGTTTTTTCGC |
| p768 | TACGGTTATCCACAGAATCAGGGGATAACGCAGGCCTCCGAAATCCTGCAGGGAAAGCCA<br>CGTTGTGTCTCAAAATCTCTGATGTTACATTGCACAAGATAAAAATATATCATCATGAACAATA<br>AAACTGTCTGCTTACATAAACAGTAATACAAGGGGTGTTATGAGCCATATTCAACGGGAAAC<br>GTCTTGCTCGAGGCCGCGATTAAATTCCAACATGGATGCTGATTTATATGGGTATAAATGGG<br>CTCGCGATAATGTCGGGCAATCAGGTGCGACAATCTATCGATTGTATGGGAAGCCCGATGC<br>GCCAGAGTTGTTTCTGAAACATGGCAAAGGTAGCGTTGCCAATGATGTTACAGATGAGATG<br>GTCAGACTAAACTGGCTGACGGAATTTATGCCTCTTCCGACCATCAAGCATTTTATCCGTAC<br>TCCTGATGATGCATGGTTACTCACCACTGCGATCCCCGGGAAAACAGCATTCCAGGTATTA<br>GAAGAATATCCTGATTCAGGTGAAAATATTGTTGATGCGCTGGCAGTGTTCCTGCGCCGGT<br>TGCATTCGATTCCTGTTTGTAATTGTCCTTTTAACAGCGATCGCGTATTTCGTCTCGCTCAG<br>GCGCAATCACGAATGAATAACGGTTTGGTTGATGCGAGTGATTTTGATGACGAGCGTAATG<br>GCTGGCCTGTTGAACAAGTCTGGAAAGAAATGCATAAGCTTTTGCCATTCTCACCGGATTC<br>AGTCGTCACTCATGGTGATTTCTCACTTGATAACCTTATTTTTGACGAGGGGAAATTAATAG<br>GTTGTATTGATGTTGGACGAGTCGGAATCGCAGACCGATACCAGGATCTTGCCATCCTATG<br>GAACTGCCTCGGTGAGTTTTCTCCTTCATTACAGAAACGGCTTTTTCAAAAATATGGTATTG<br>ATAATCCTGATATGAATAAATTGCAGTTTCATTTGATGCTCGATGAGTTTTTCTAAATCAGAAT<br>TGGTTAATTGGTTGTGGAAACGTAACTGTCAGACCAAGTTTACTCATATATACTTTAGATTGA<br>TTTAAAACTTCATTTTTAATTTAAAAGGATCTAGGTGAAGATCCTTTTTGATAATCTCATGACC<br>AAAATCCCTTAACGTGAGTTTTCGTTCCACTGAGCGTCAGACCCCGTTGACAGTAAGACGG<br>GTAAGCCTGTTGATGATACCGCTGCCTTACTGGGTGCATTAGCCAGTCTGAATGACCTGTC<br>ACGGGATAATCCGAAGTGGTCAGACTGGAAAATCAGAGGGCAGGAACTGCTGAACAGCAA<br>AAAGTCAGATAGCACCACATAGCAGACCCGCCATAAAACGCCCTGAGAAGCCCGTGACGG<br>GCTTTTCTTGTATTATGGGTAGTTTCCTTGCATGAATCCATAAAAGGCGCCTGTAGTGCCATT<br>TACCCCCATTCACTGCCAGAGCCGTGAGCGCAGCGAACTGAATGTCACGAAAAAGACAGC<br>GACTCAGGTGCCTGATGGTCGGAGACAAAAGGAATATTCAGCGATTTGCCCGATTGCGGC<br>CGCAACCGAGCTTGCGAGGGTGCTACTTAAGCCTTTAGGGTTTTAAGGTCTGTTTTGTAGA<br>GGAGCAAACAGCGTTTGCGACATCCTTTTGTAATACTGCGGAACTGACTAAAGTAGTGAGT<br>TATACACAGGGCTGGGATCTATTCTTTTTATCTTTTTTTATTCTTTCTTTATTCTATAAATTATAA<br>CCACTTGAATATAAACAAAAAAAACACACAAAGGTCTAGCGGAATTTACAGAGGGTCTAGCA<br>GAATTTACAAGTTTTCCAGCAAAGGTCTAGCAGAATTTACAGATACCCACAACTCAAAGGAA<br>AAGGACTAGTAATTATCATTGACTAGCCCATCTCAATTGGTATAGTGATTAAAATCACCTAGA |

| | |
|---|---|
| | CCAATTGAGATGTATGTCTGAATTAGTTGTTTTCAAAGCAAATGAACTAGCGATTAGTCGCTA<br>TGACTTAACGGAGCATGAAACCAAGCTAATTTTATGCTGTGTGGCACTACTCAACCCCACG<br>ATTGAAAACCCTACAAGGAAAGAACGGACGGTATCGTTCACTTATAACCAATACGCTCAGAT<br>GATGAACATCAGTAGGGAAAATGCTTATGGTGTATTAGCTAAAGCAACCAGAGAGCTGATGA<br>CGAGAACTGTGGAAATCAGGAATCCTTTGGTTAAAGGCTTTGAGATTTTCCAGTGGACAAA<br>CTATGCCAAGTTCTCAAGCGAAAAATTAGAATTAGTTTTTAGTGAAGAGATATTGCCTTATCT<br>TTTCCAGTTAAAAAAATTCATAAAATATAATCTGGAACATGTTAAGTCTTTTGAAAACAAATAC<br>TCTATGAGGATTTATGAGTGGTTATTAAAAGAACTAACACAAAAGAAAACTCACAAGGCAAA<br>TATAGAGATTAGCCTTGATGAATTTAAGTTCATGTTAATGCTTGAAAATAACTACCATGAGTTT<br>AAAAGGCTTAACCAATGGGTTTTGAAACCAATAAGTAAAGATTTAAACACTTACAGCAATATG<br>AAATTGGTGGTTGATAAGCGAGGCCGCCCGACTGATACGTTGATTTTCCAAGTTGAACTAG<br>ATAGACAAATGGATCTCGTAACCGAACTTGAGAACAACCAGATAAAAATGAATGGTGACAAA<br>ATACCAACAACCATTACATCAGATTCCTACCTACGTAACGGACTAAGAAAAACACTACACGA<br>TGCTTTAACTGCAAAAATTCAGCTCACCAGTTTTGAGGCAAAATTTTTGAGTGACATGCAAA<br>GTAAGTATGATCTCAATGGTTCGTTCTCATGGCTCACGCAAAAACAACGAACCACACTAGA<br>GAACATACTGGCTAAATACGGAAGGATCTGAGGTTCTTATGGCTCTTGTATCTATCAGTGAA<br>GCATCAAGACTAACAAACAAAAGTAGAACAACTGTTCACCGTTACATATCAAAGGGAAAACT<br>GTCCATATGCACAGATGAAAACGGTGTAAAAAAGATAGATACATCAGAGCTTTTACGAGTTT<br>TTGGTGCATTTAAAGCTGTTCACCATGAACAGATCGACAATGTAACAGATGAACAGCATGTA<br>ACACCTAATAGAACAGGTGAAACCAGTAAAACAAAGCAACTAGAACATGAAATTGAACACCT<br>GAGACAACTTGTTACAGCTCAACAGTCACACATAGACAGCCTGAAACAGGCGATGCTGCTT<br>ATCGAATCAAAGCTGCCGACAACACGGGAGCCAGTGACGCCTCCCGTGGGGAAAAAATC<br>ATGGCAATTCTGGAAGAAATAGCGCTTTCATCCGGCAAACCTGAAACCGGATCTGCGATTC<br>TGATAACAAACTAGCAACACCAGAACAGCCCGTTTGCGGGCAGTAAAACCCGTTTTCATCA<br>TATCTGGCGTTAATGGAGTTTTTACGGCTAGCTCAGTCCTAGGTACAATGCTAGCGTTTTCA<br>TTAAAGAGGAGAAAGGAAGCCATGGAGGACGTTATCAAAGAGTTCATGCGTTTCAAAGTTC<br>GTATGGAAGGTAGCGTTAACGGTCACGAGTTCGAAATCGAAGGTGAGGGTGAAGGTCGTC<br>CGTACGAAGGTACTCAAACCGCTAAACTGAAAGTTACCAAAGGTGGCCCTCTGCCGTTCG<br>CTTGGGACATCCTGTCCCCGCAGTTCCAGTACGGTAGCAAGGCTTACGTTAAACATCCGG<br>CTGACATTCCGGACTACTTAAAGCTGTCCTTTCCGGAAGGTTTCAAATGGGAGCGTGTTAT<br>GAACTTCGAGGACGGCGGTGTTGTTACCGTTACGCAGGACTCCTCCCTGCAAGACGGTG<br>AGTTCATCTACAAAGTTAAACTGCGTGGTACGAACTTCCCGTCCGACGGCCCGGTTATGCA<br>GAAAAAAACTATGGGTTGGGAGGCTTCCACCGAACGTATGTATCCGGAGGACGGTGCGCT<br>GAAAGGTGAAATCAAAATGCGTCTGAAACTGAAAGACGGCGGTCACTACGACGCGGAAGT<br>TAAAACTACCTACATGGCTAAAAAACCTGTTCAACTGCCGGGCGCTTACAAAACCGACATC<br>AAACTGGACATCACCTCCCACAACGAGGACTATACCATCGTTGAACAGTACGAACGCGCG<br>GAAGGTCGTCACTCCACTGGTGCTTAAGTTTAGCCGAACGCCCGCAAAAAACCCCGCTTC<br>GGCGGGGTTTTTTCGC |
| p840 | GTAACTGTCAGACCAAGTTTACTCATATATACTTTAGATTGATTTAAAACTTCATTTTTAATTTA<br>AAAGGATCTAGGTGAAGATCCTTTTTGATAATCTCATGACCAAAATCCCTTAACGTGAGTTTT<br>CGTTCCACTGAGCGTCAGACCCCGTTGACAGTAAGACGGGTAAGCCTGTTGATGATACCG<br>CTGCCTTACTGGGTGCATTAGCCAGTCTGAATGACCTGTCACGGGATAATCCGAAGTGGTC<br>AGACTGGAAAATCAGAGGGCAGGAACTGCTGAACAGCAAAAAGTCAGATAGCACCACATA<br>GCAGACCCGCCATAAAACGCCCTGAGAAGCCCGTGACGGGCTTTTCTTGTATTATGGGTA<br>GTTTCCTTGCATGAATCCATAAAAGGCGCCTGTAGTGCCATTTACCCCCATTCACTGCCAGA<br>GCCGTGAGCGCAGCGAACTGAATGTCACGAAAAAGACAGCGACTCAGGTGCCTGATGGT<br>CGGAGACAAAAGGAATATTCAGCGATTTGCCCGATTGCGGCCGCAACCGAGCTTGCGAGG<br>GTGCTACTTAAGCCTTTAGGGTTTTAAGGTCTGTTTTGTAGAGGAGCAAACAGCGTTTGCG |

| | |
|---|---|
| | ACATCCTTTTGTAATACTGCGGAACTGACTAAAGTAGTGAGTTATACACAGGGCTGGGATCT<br>ATTCTTTTTATCTTTTTTTATTCTTTCTTTATTCTATAAATTATAACCACTTGAATATAAACAAAAA<br>AAACACACAAAGGTCTAGCGGAATTTACAGAGGGTCTAGCAGAATTTACAAGTTTTCCAGC<br>AAAGGTCTAGCAGAATTTACAGATACCCACAACTCAAAGGAAAAGGACTAGTAATTATCATT<br>GACTAGCCCATCTCAATTGGTATAGTGATTAAAATCACCTAGACCAATTGAGATGTATGTCTG<br>AATTAGTTGTTTTCAAAGCAAATGAACTAGCGATTAGTCGCTATGACTTAACGGAGCATGAA<br>ACCAAGCTAATTTTATGCTGTGTGGCACTACTCAACCCCACGATTGAAAACCCTACAAGGA<br>AAGAACGGACGGTATCGTTCACTTATAACCAATACGCTCAGATGATGAACATCAGTAGGGAA<br>AATGCTTATGGTGTATTAGCTAAAGCAACCAGAGAGCTGATGACGAGAACTGTGGAAATCA<br>GGAATCCTTTGGTTAAAGGCTTTGAGATTTTCCAGTGGACAAACTATGCCAAGTTCTCAAG<br>CGAAAAATTAGAATTAGTTTTTAGTGAAGAGATATTGCCTTATCTTTTCCAGTTAAAAAAATTC<br>ATAAAATATAATCTGGAACATGTTAAGTCTTTTGAAAACAAATACTCTATGAGGATTTATGAGT<br>GGTTATTAAAAGAACTAACACAAAAGAAAACTCACAAGGCAAATATAGAGATTAGCCTTGAT<br>GAATTTAAGTTCATGTTAATGCTTGAAAATAACTACCATGAGTTTAAAAGGCTTAACCAATGG<br>GTTTTGAAACCAATAAGTAAAGATTTAAACACTTACAGCAATATGAAATTGGTGGTTGATAAG<br>CGAGGCCGCCCGACTGATACGTTGATTTTCCAAGTTGAACTAGATAGACAAATGGATCTCG<br>TAACCGAACTTGAGAACAACCAGATAAAAATGAATGGTGACAAAATACCAACAACCATTACA<br>TCAGATTCCTACCTACGTAACGGACTAAGAAAAACACTACACGATGCTTTAACTGCAAAAAT<br>TCAGCTCACCAGTTTTGAGGCAAAATTTTTGAGTGACATGCAAAGTAAGTATGATCTCAATG<br>GTTCGTTCTCATGGCTCACGCAAAAACAACGAACCACACTAGAGAACATACTGGCTAAATA<br>CGGAAGGATCTGAGGTTCTTATGGCTCTTGTATCTATCAGTGAAGCATCAAGACTAACAAAC<br>AAAAGTAGAACAACTGTTCACCGTTACATATCAAAGGGAAAACTGTCCATATGCACAGATGA<br>AAACGGTGTAAAAAAGATAGATACATCAGAGCTTTTACGAGTTTTTGGTGCATTTAAAGCTG<br>TTCACCATGAACAGATCGACAATGTAACAGATGAACAGCATGTAACACCTAATAGAACAGGT<br>GAAACCAGTAAAACAAAGCAACTAGAACATGAAATTGAACACCTGAGACAACTTGTTACAG<br>CTCAACAGTCACACATAGACAGCCTGAAACAGGCGATGCTGCTTATCGAATCAAAGCTGCC<br>GACAACACGGGAGCCAGTGACGCCTCCCGTGGGGAAAAAATCATGGCAATTCTGGAAGA<br>AATAGCGCTTTCATCCGGCAAACCTGAAACCGGATCTGCGATTCTGATAACAAACTAGCAA<br>CACCAGAACAGCCCGTTTGCGGGCAGTAAAACCCGTTTTGCCGAAAACGTAAAGCTTCAG<br>CTGTTTTACGGCTAGCTCAGTCCTAGGTACAATGCTAGCGTTTTCATTAAAGAGGAGAAAG<br>GAAGCCATGGAGGACGTTATCAAAGAGTTCATGCGTTTCAAAGTTCGTATGGAAGGTAGCG<br>TTAACGGTCACGAGTTCGAAATCGAAGGTGAGGGTGAAGGTCGTCCGTACGAAGGTACTC<br>AAACCGCTAAACTGAAAGTTACCAAAGGTGGCCCTCTGCCGTTCGCTTGGGACATCCTGT<br>CCCCGCAGTTCCAGTACGGTAGCAAGGCTTACGTTAAACATCCGGCTGACATTCCGGACTA<br>CTTAAAGCTGTCCTTTCCGGAAGGTTTCAAATGGGAGCGTGTTATGAACTTCGAGGACGG<br>CGGTGTTGTTACCGTTACGCAGGACTCCTCCCTGCAAGACGGTGAGTTCATCTACAAAGTT<br>AAACTGCGTGGTACGAACTTCCCGTCCGACGGCCCGGTTATGCAGAAAAAAACTATGGGT<br>TGGGAGGCTTCCACCGAACGTATGTATCCGGAGGACGGTGCGCTGAAAGGTGAAATCAAA<br>ATGCGTCTGAAACTGAAAGACGGCGGTCACTACGACGCGGAAGTTAAAACTACCTACATG<br>GCTAAAAAACCTGTTCAACTGCCGGGCGCTTACAAAACCGACATCAAACTGGACATCACCT<br>CCCACAACGAGGACTATACCATCGTTGAACAGTACGAACGCGCGGAAGGTCGTCACTCCA<br>CTGGTGCTTAAGTTTAGCCGAACGCCCGCAAAAAACCCCGCTTCGGCGGGGTTTTTTCGC<br>TACGGTTATCCACAGAATCAGGGGATAACGCAGGCCTCCGAAATCCTGCAGGGAAAGCCA<br>CGTTGTGTCTCAAAATCTCTGATGTTACATTGCACAAGATAAAAATATATCATCATGAACAATA<br>AAACTGTCTGCTTACATAAACAGTAATACAAGGGGTGTTATGAGCCATATTCAACGGGAAAC<br>GTCTTGCTCGAGGCCGCGATTAAATTCCAACATGGATGCTGATTTATATGGGTATAAATGGG<br>CTCGCGATAATGTCGGGCAATCAGGTGCGACAATCTATCGATTGTATGGGAAGCCCGATGC<br>GCCAGAGTTGTTTCTGAAACATGGCAAAGGTAGCGTTGCCAATGATGTTACAGATGAGATG |

| | |
|---|---|
| | GTCAGACTAAACTGGCTGACGGAATTTATGCCTCTTCCGACCATCAAGCATTTTATCCGTAC<br>TCCTGATGATGCATGGTTACTCACCACTGCGATCCCCGGGAAAACAGCATTCCAGGTATTA<br>GAAGAATATCCTGATTCAGGTGAAAATATTGTTGATGCGCTGGCAGTGTTCCTGCGCCGGT<br>TGCATTCGATTCCTGTTTGTAATTGTCCTTTTAACAGCGATCGCGTATTTCGTCTCGCTCAG<br>GCGCAATCACGAATGAATAACGGTTTGGTTGATGCGAGTGATTTTGATGACGAGCGTAATG<br>GCTGGCCTGTTGAACAAGTCTGGAAAGAAATGCATAAGCTTTTGCCATTCTCACCGGATTC<br>AGTCGTCACTCATGGTGATTTCTCACTTGATAACCTTATTTTTGACGAGGGGAAATTAATAG<br>GTTGTATTGATGTTGGACGAGTCGGAATCGCAGACCGATACCAGGATCTTGCCATCCTATG<br>GAACTGCCTCGGTGAGTTTTCTCCTTCATTACAGAAACGGCTTTTTCAAAAATATGGTATTG<br>ATAATCCTGATATGAATAAATTGCAGTTTCATTTGATGCTCGATGAGTTTTTCTAAATCAGAAT<br>TGGTTAATTGGTTGTGGAAAC |
| p764 | TACGGTTATCCACAGAATCAGGGGATAACGCAGGCCTCCGAAATCCTGCAGGGAAAGCCA<br>CGTTGTGTCTCAAAATCTCTGATGTTACATTGCACAAGATAAAAATATATCATCATGAACAATA<br>AAACTGTCTGCTTACATAAACAGTAATACAAGGGGTGTTATGAGCCATATTCAACGGGAAAC<br>GTCTTGCTCGAGGCCGCGATTAAATTCCAACATGGATGCTGATTTATATGGGTATAAATGGG<br>CTCGCGATAATGTCGGGCAATCAGGTGCGACAATCTATCGATTGTATGGGAAGCCCGATGC<br>GCCAGAGTTGTTTCTGAAACATGGCAAAGGTAGCGTTGCCAATGATGTTACAGATGAGATG<br>GTCAGACTAAACTGGCTGACGGAATTTATGCCTCTTCCGACCATCAAGCATTTTATCCGTAC<br>TCCTGATGATGCATGGTTACTCACCACTGCGATCCCCGGGAAAACAGCATTCCAGGTATTA<br>GAAGAATATCCTGATTCAGGTGAAAATATTGTTGATGCGCTGGCAGTGTTCCTGCGCCGGT<br>TGCATTCGATTCCTGTTTGTAATTGTCCTTTTAACAGCGATCGCGTATTTCGTCTCGCTCAG<br>GCGCAATCACGAATGAATAACGGTTTGGTTGATGCGAGTGATTTTGATGACGAGCGTAATG<br>GCTGGCCTGTTGAACAAGTCTGGAAAGAAATGCATAAGCTTTTGCCATTCTCACCGGATTC<br>AGTCGTCACTCATGGTGATTTCTCACTTGATAACCTTATTTTTGACGAGGGGAAATTAATAG<br>GTTGTATTGATGTTGGACGAGTCGGAATCGCAGACCGATACCAGGATCTTGCCATCCTATG<br>GAACTGCCTCGGTGAGTTTTCTCCTTCATTACAGAAACGGCTTTTTCAAAAATATGGTATTG<br>ATAATCCTGATATGAATAAATTGCAGTTTCATTTGATGCTCGATGAGTTTTTCTAAATCAGAAT<br>TGGTTAATTGGTTGTGGAAACGTAACTGTCAGACCAAGTTTACTCATATATACTTTAGATTGA<br>TTTAAAACTTCATTTTTAATTTAAAAGGATCTAGGTGAAGATCCTTTTTGATAATCTCATGACC<br>AAAATCCCTTAACGTGAGTTTTCGTTCCACTGAGCGTCAGACCCCGTTGACAGTAAGACGG<br>GTAAGCCTGTTGATGATACCGCTGCCTTACTGGGTGCATTAGCCAGTCTGAATGACCTGTC<br>ACGGGATAATCCGAAGTGGTCAGACTGGAAAATCAGAGGGCAGGAACTGCTGAACAGCAA<br>AAAGTCAGATAGCACCACATAGCAGACCCGCCATAAAACGCCCTGAGAAGCCCGTGACGG<br>GCTTTTCTTGTATTATGGGTAGTTTCCTTGCATGAATCCATAAAAGGCGCCTGTAGTGCCATT<br>TACCCCCATTCACTGCCAGAGCCGTGAGCGCAGCGAACTGAATGTCACGAAAAAGACAGC<br>GACTCAGGTGCCTGATGGTCGGAGACAAAAGGAATATTCAGCGATTTGCCCGATTGCGGC<br>CGCAACCGAGCTTGCGAGGGTGCTACTTAAGCCTTTAGGGTTTTAAGGTCTGTTTTGTAGA<br>GGAGCAAACAGCGTTTGCGACATCCTTTTGTAATACTGCGGAACTGACTAAAGTAGTGAGT<br>TATACACAGGGCTGGGATCTATTCTTTTTATCTTTTTTTATTCTTTCTTTATTCTATAAATTATAA<br>CCACTTGAATATAAACAAAAAAAACACACAAAGGTCTAGCGGAATTTACAGAGGGTCTAGCA<br>GAATTTACAAGTTTTCCAGCAAAGGTCTAGCAGAATTTACAGATACCCACAACTCAAAGGAA<br>AAGGACTAGTAATTATCATTGACTAGCCCATCTCAATTGGTATAGTGATTAAAATCACCTAGA<br>CCAATTGAGATGTATGTCTGAATTAGTTGTTTTCAAAGCAAATGAACTAGCGATTAGTCGCTA<br>TGACTTAACGGAGCATGAAACCAAGCTAATTTTATGCTGTGTGGCACTACTCAACCCCACG<br>ATTGAAAACCCTACAAGGAAAGAACGGACGGTATCGTTCACTTATAACCAATACGCTCAGAT<br>GATGAACATCAGTAGGGAAAATGCTTATGGTGTATTAGCTAAAGCAACCAGAGAGCTGATGA<br>CGAGAACTGTGGAAATCAGGAATCCTTTGGTTAAAGGCTTTGAGATTTTCCAGTGGACAAA<br>CTATGCCAAGTTCTCAAGCGAAAAATTAGAATTAGTTTTTAGTGAAGAGATATTGCCTTATCT |

| | |
|---|---|
| | TTTCCAGTTAAAAAAATTCATAAAATATAATCTGGAACATGTTAAGTCTTTTGAAAACAAATAC<br>TCTATGAGGATTTATGAGTGGTTATTAAAAGAACTAACACAAAAGAAAACTCACAAGGCAAA<br>TATAGAGATTAGCCTTGATGAATTTAAGTTCATGTTAATGCTTGAAAATAACTACCATGAGTTT<br>AAAAGGCTTAACCAATGGGTTTTGAAACCAATAAGTAAAGATTTAAACACTTACAGCAATATG<br>AAATTGGTGGTTGATAAGCGAGGCCGCCCGACTGATACGTTGATTTTCCAAGTTGAACTAG<br>ATAGACAAATGGATCTCGTAACCGAACTTGAGAACAACCAGATAAAAATGAATGGTGACAAA<br>ATACCAACAACCATTACATCAGATTCCTACCTACGTAACGGACTAAGAAAAACACTACACGA<br>TGCTTTAACTGCAAAAATTCAGCTCACCAGTTTTGAGGCAAAATTTTTGAGTGACATGCAAA<br>GTAAGTATGATCTCAATGGTTCGTTCTCATGGCTCACGCAAAAACAACGAACCACACTAGA<br>GAACATACTGGCTAAATACGGAAGGATCTGAGGTTCTTATGGCTCTTGTATCTATCAGTGAA<br>GCATCAAGACTAACAAACAAAAGTAGAACAACTGTTCACCGTTACATATCAAAGGGAAAACT<br>GTCCATATGCACAGATGAAAACGGTGTAAAAAAGATAGATACATCAGAGCTTTTACGAGTTT<br>TTGGTGCATTTAAAGCTGTTCACCATGAACAGATCGACAATGTAACAGATGAACAGCATGTA<br>ACACCTAATAGAACAGGTGAAACCAGTAAAACAAAGCAACTAGAACATGAAATTGAACACCT<br>GAGACAACTTGTTACAGCTCAACAGTCACACATAGACAGCCTGAAACAGGCGATGCTGCTT<br>ATCGAATCAAAGCTGCCGACAACACGGGAGCCAGTGACGCCTCCCGTGGGGAAAAAATC<br>ATGGCAATTCTGGAAGAAATAGCGCTTTCATCCGGCAAACCTGAAACCGGATCTGCGATTC<br>TGATAACAAACTAGCAACACCAGAACAGCCCGTTTGCGGGCAGTAAAACCCGTTTTCCATG<br>ACAACGGACAGCAGTTATACGCAAATCAGTCGTCACTCACTGGTTTACGGCTAGCTCAGTC<br>CTAGGTACAATGCTAGCGTTTTCATTAAAGAGGAGAAAGGAAGCCATGGAGGACGTTATCA<br>AAGAGTTCATGCGTTTCAAAGTTCGTATGGAAGGTAGCGTTAACGGTCACGAGTTCGAAAT<br>CGAAGGTGAGGGTGAAGGTCGTCCGTACGAAGGTACTCAAACCGCTAAACTGAAAGTTAC<br>CAAAGGTGGCCCTCTGCCGTTCGCTTGGGACATCCTGTCCCCGCAGTTCCAGTACGGTAG<br>CAAGGCTTACGTTAAACATCCGGCTGACATTCCGGACTACTTAAAGCTGTCCTTTCCGGAA<br>GGTTTCAAATGGGAGCGTGTTATGAACTTCGAGGACGGCGGTGTTGTTACCGTTACGCAG<br>GACTCCTCCCTGCAAGACGGTGAGTTCATCTACAAAGTTAAACTGCGTGGTACGAACTTCC<br>CGTCCGACGGCCCGGTTATGCAGAAAAAAACTATGGGTTGGGAGGCTTCCACCGAACGTA<br>TGTATCCGGAGGACGGTGCGCTGAAAGGTGAAATCAAAATGCGTCTGAAACTGAAAGACG<br>GCGGTCACTACGACGCGGAAGTTAAAACTACCTACATGGCTAAAAAACCTGTTCAACTGCC<br>GGGCGCTTACAAAACCGACATCAAACTGGACATCACCTCCCACAACGAGGACTATACCATC<br>GTTGAACAGTACGAACGCGCGGAAGGTCGTCACTCCACTGGTGCTTAAGTTTAGCCGAAC<br>GCCCGCAAAAAACCCCGCTTCGGCGGGGTTTTTTCGC |
| p1400 | GGAGAACTGGCGCGAATATCTGGCCCGTGAGCGCCAGAATCTTTCTGATGGTCTGGTCAT<br>TGAGCTTCCGGTAAAGCAAAAGGCGCAACTTTCGCAGATGGCGGACAGTGAGCGCGCGC<br>AGCTGCTTGCCGATCGCTTTGATGGCGTTTGCGTACATCCTGAAAGTGAAATCGTTCACGT<br>ATGGTGCGGCGGGGTATGGTGTCCGGTCAGCACAATGGAGCTGAGCCGCGAAATGGTGG<br>CGATCTATTCAGAGCACAGGGCCACTTTCAGCAAGCGCGTAATCAATAACGCCGTGGAAG<br>CGTTAAAAGTTATTGCCGAACCAATGGGCGAGCCGTCCGGCGATTTGCTGCCGTTCGCCA<br>ATGGTGCGCTTGACCTGAAAACGGGGGAATTTTCCCCGCACACGCCGGAGAACTGGATCA<br>CCACGCACAACGGCATTGAGTACACGCCACCAGCACCCGGGGAGAACATCCGCGATAAC<br>GCGCCAAACTTTCATAAATGGCTTGAGCACGCAGCCGGAAAAGACCCGCGCAAGATGATG<br>CGTATATGTGCCGCGCTGTACATGATTATGGCGAACCGGTACGACTGGCAGATGTTTATTGA<br>GGCCACCGGAGACGGCGGGAGCGGTAAAAGTACATTCACACACATAGCCAGCCTTCTGG<br>CAGGGAAACAAAACACGGTAAGCGCTGAAATGACATCGCTTGATGATGCTGGTGGGCGTG<br>CGCAGGTTGTCGGGAGTCGTCTTATCGTCCTGGCAGACCAGCCGAAATATACAGGCGAAG<br>GAACGGGCATCAAGAAAATCACGGGCGGCGACCCCGTGGAAATTAACCCGAAATATGAAA<br>AGCGTTTTACGGCGGTAATCAGGGCGGTGGTGCTGGCAACCAATAACAATCCGATGATATT<br>CACCGAACGGGCCGGAGGTGTGGCACGTCGTCGGGTGATATTCCGGTTCGATAACATCGT |

| | |
|---|---|
| | AAGCGAGGCAGAAAAAGACAGGGAGCTACCGGAAAAGATCGCGGCTGAAATCCCTGTCAT<br>TATCCGCCGCTTGCTGGCGAACTTTGCCGACCCTGAAAAGGCACGGGCTTTACTCATTGA<br>ACAGCGTGACGGTGATGAAGCACTGGCAATAAAGCAACAGACGGATCCGGTTATTGAGTT<br>TTGCCAGTTCCTGAATTTTCTGGAGGAAGCACGCGGCCTGATGATGGGCGGCGGTGGCG<br>ATTCAGTGAAGTACACGACCAGAAACAGCCTTTACCGCGTCTATCTGGCGTTTATGGCGTA<br>CGCAGGCAGGAGCAAACCGCTAAACGTAAATGACTTTGGCAAGGCTATGAAGCCAGCCGC<br>GAAAGTTTACGGACATGAATATATTACGCGGAAAGTTAAAGGAGTAACGCAGACTAACGCAA<br>TAACAACAGACGATTGCGACGCGTTTTTATAATGACGCATCCTCACGATAATATCCGGGTAG<br>GACGAACAATAAGGCCGCAAATCGCGGCCTTTTTTATTGATAACAAAAGGACAGTTTTCCCT<br>TTGATATGTAACGGTGAACAGTTGTTCTACTTTTGTTTGTTAGTCTTGATGCTTCACTGATAG<br>ATACAAGAGCCATAAGAACCTCAGATCCTTCCGTATTTAGCCAGTATGTTCTCTAGTGTGGT<br>TCGTTGTTTTTGCGTGAGCCATGAGAACGAACCATTGAGATCATACTTACTTTGCATGTCAC<br>TCAAAAATTTTGCCTCAAAACTGGTGAGCTGAATTTTTGCAGTTAAAGCATCGTGTAGTGTT<br>TTTCTTAGTCCGTTATGTAGGTAGGAATCTGATGTAATGGTTGTTGGTATTTTGTCACCATTC<br>ATTTTTATCTGGTTGTTCTCAAGTTCGGTTACGAGATCCATTTGTCTATCTAGTTCAACTTGG<br>AAAATCAACGTATCAGTCGGGCGGCCTCGCTTATCAACCACCAATTTCATATTGCTGTAAGT<br>GTTTAAATCTTTACTTATTGGTTTCAAAACCCATTGGTTAAGCCTTTTAAACTCATGGTAGTTA<br>TTTTCAAGCATTAACATGAACTTAAATTCATCAAGGCTAATCTCTATATTTGCCTTGTGAGTTT<br>TCTTTTGTGTTAGTTCTTTTAATAACCACTCATAAATCCTCATAGAGTATTTGTTTTCAAAAGA<br>CTTAACATGTTCCAGATTATATTTTATGAATTTTTTTAACTGGAAAAGATAAGGCAATATCTCTT<br>CACTAAAAACTAATTCTAATTTTTCGCTTGAGAACTTGGCATAGTTTGTCCACTGGAAAATCT<br>CAAAGCCTTTAACCAAAGGATTCCTGATTTCCACAGTTCTCGTCATCAGCTCTCTGGTTGCT<br>TTAGCTAATACACCATAAGCATTTTCCCTACTGATGTTCATCATCTGAGCGTATTGGTTATAAG<br>TGAACGATACTGTCCGTTCTTTCCTTGTAGGGTTTTCAATCGTGGGGTTGAGTAGTGCCAC<br>ACAGCATAAAATTAGCTTGGTTTCATGCTCCGTTAAGTCATAGCGACTAATCGCTAGTTCATT<br>TGCTTTGAAAACAACTAATTCAGACATACATCTCAATTGGTCTAGGTGATTTTAATCACTATAC<br>CAATTGAGATGGGCTAGTCAATGATAATTACTAGTCCTTTTCCTTTGAGTTGTGGGTATCTGT<br>AAATTCTGCTAGACCTTTGCTGGAAAACTTGTAAATTCTGCTAGACCCTCTGTAAATTCCGC<br>TAGACCTTTGTGTGTTTTTTTTGTTTATATTCAAGTGGTTATAATTTATAGAATAAAGAAAGAAT<br>AAAAAAAGATAAAAAGAATAGATCCCAGCCCTGTGTATAACTCACTACTTTAGTCAGTTCCG<br>CAGTATTACAAAAGGATGTCGCAAACGCTGTTTGCTCCTCTACAAAACAGACCTTAAAACCC<br>TAAAGGCTTAAGTAGCACCCTCGCAAGCTCGGGCAAATCGCTGAATATTCCTTTTGTCTCC<br>GACCATCAGGCACCTGAGTCGCTGTCTTTTTCGTGACATTCAGTTCGCTGCGCCCGACGT<br>ACGGTGAATCTGATTCGTTACCAATTGACATGATACGAAACGTACCGTATCGTTAAGGTCGT<br>GATTAAACGATCCCGTTTTCAAAAATGAAACTGGCACCGAACGTAAAACAGCAGTCACGCG<br>GCATAAAACACAAAGAAACAGAAGTCATTATTTTTGCGGGTAGTGATGCCTGGTCACACGC<br>AAAACAATGGCAGGAACATGACGCGCGTATGGCCGGAGATAATGAGCCTCCTGTGTGGCT<br>TGGGGAGCAGCAGTTATCCGAACTGGATAAGCTGCAAATTGTGCCGGAAGGCAGAAAATC<br>CGTGCGCATATTCAGGGCCGGATATCTTGCGCCAGTAATGATAAAGGCGATTGGTCAGAAG<br>CTGGCGGCGGCAGGCGTACAGGATGCAAATTTTTACCCTGATGGTATGCACGGTCAGAAG<br>GT |
| p872 | TCAGATCCTTCCGTATTTAGCCAGTATGTTCTCTAGTGTGGTTCGTTGTTTTTGCGTGAGCC<br>ATGAGAACGAACCATTGAGATCATACTTACTTTGCATGTCACTCAAAAATTTTGCCTCAAAAC<br>TGGTGAGCTGAATTTTTGCAGTTAAAGCATCGTGTAGTGTTTTTCTTAGTCCGTTACGTAGG<br>TAGGAATCTGATGTAATGGTTGTTGGTATTTTGTCACCATTCATTTTTATCTGGTTGTTCTCAA<br>GTTCGGTTACGAGATCCATTTGTCTATCTAGTTCAACTTGGAAAATCAACGTATCAGTCGGG<br>CGGCCTCGCTTATCAACCACCAATTTCATATTGCTGTAAGTGTTTAAATCTTTACTTATTGGT<br>TTCAAAACCCATTGGTTAAGCCTTTTAAACTCATGGTAGTTATTTTCAAGCATTAACATGAAC |

| | |
|---|---|
| | TTAAATTCATCAAGGCTAATCTCTATATTTGCCTTGTGAGTTTTCTTTTGTGTTAGTTCTTTTA<br>ATAACCACTCATAAATCCTCATAGAGTATTTGTTTTCAAAAGACTTAACATGTTCCAGATTATA<br>TTTTATGAATTTTTTTAACTGGAAAAGATAAGGCAATATCTCTTCACTAAAAACTAATTCTAATT<br>TTTCGCTTGAGAACTTGGCATAGTTTGTCCACTGGAAAATCTCAAAGCCTTTAACCAAAGG<br>ATTCCTGATTTCCACAGTTCTCGTCATCAGCTCTCTGGTTGCTTTAGCTAATACACCATAAG<br>CATTTTCCCTACTGATGTTCATCATCTGAGCGTATTGGTTATAAGTGAACGATACCGTCCGTT<br>CTTTCCTTGTAGGGTTTTCAATCGTGGGGTTGAGTAGTGCCACACAGCATAAAATTAGCTTG<br>GTTTCATGCTCCGTTAAGTCATAGCGACTAATCGCTAGTTCATTTGCTTTGAAAACAACTAAT<br>TCAGACATACATCTCAATTGGTCTAGGTGATTTTAATCACTATACCAATTGAGATGGGCTAGT<br>CAATGATAATTACTAGTCCTTTTCCTTTGAGTTGTGGGTATCTGTAAATTCTGCTAGACCTTT<br>GCTGGAAAACTTGTAAATTCTGCTAGACCCTCTGTAAATTCCGCTAGACCTTTGTGTGTTTT<br>TTTTGTTTATATTCAAGTGGTTATAATTTATAGAATAAAGAAAGAATAAAAAAAGATAAAAAGAA<br>TAGATCCCAGCCCTGTGTATAACTCACTACTTTAGTCAGTTCCGCAGTATTACAAAAGGATG<br>TCGCAAACGCTGTTTGCTCCTCTACAAAACAGACCTTAAAACCCTAAAGGCTTAAGTAGCA<br>CCCTCGCAAGCTCGGTTGCGGCCGCAATCGGGCAAATCGCTGAATATTCCTTTTGTCTCC<br>GACCATCAGGCACCTGAGTCGCTGTCTTTTTCGTGACATTCAGTTCGCTGCGCTCACGGC<br>TCTGGCAGTGAATGGGGGTAAATGGCACTACAGGCGCCTTTTATGGATTCATGCAAGGAAA<br>CTACCCATAATACAAGAAAAGCCCGTCACGGGCTTCTCAGGGCGTTTTATGGCGGGTCTG<br>CTATGTGGTGCTATCTGACTTTTTGCTGTTCAGCAGTTCCTGCCCTCTGATTTTCCAGTCTG<br>ACCACTTCGGATTATCCCGTGACAGGTCATTCAGACTGGCTAATGCACCCAGTAAGGCAGC<br>GGTATCATCAACGGGGTCTGACGCTCAGTGGAACGAAAACTCACGTTAAGGGATTTTGGT<br>CATGAGATTATCAAAAAGGATCTTCACCTAGATCCTTTTAAATTAAAAATGAAGTTTTAAATCA<br>ATCTAAAGTATATATGAGTAAACTTGGTCTGACAGTTACGTTTCCACAACCAATTAACCAATT<br>CTGATTTAGAAAAACTCATCGAGCATCAAATGAAACTGCAATTTATTCATATCAGGATTATCAA<br>TACCATATTTTTGAAAAAGCCGTTTCTGTAATGAAGGAGAAAACTCACCGAGGCAGTTCCAT<br>AGGATGGCAAGATCCTGGTATCGGTCTGCGATTCCGACTCGTCCAACATCAATACAACCTA<br>TTAATTTCCCCTCGTCAAAAATAAGGTTATCAAGTGAGAAATCACCATGAGTGACGACTGAA<br>TCCGGTGAGAATGGCAAAAGCTTATGCATTTCTTTCCAGACTTGTTCAACAGGCCAGCCAT<br>TACGCTCGTCATCAAAATCACTCGCATCAACCAAACCGTTATTCATTCGTGATTGCGCCTGA<br>GCGAGACGAAATACGCGATCGCTGTTAAAAGGACAATTACAAACAGGAATCGAATGCAACC<br>GGCGCAGGAACACTGCCAGCGCATCAACAATATTTTCACCTGAATCAGGATATTCTTCTAAT<br>ACCTGGAATGCTGTTTTCCCGGGGATCGCAGTGGTGAGTAACCATGCATCATCAGGAGTA<br>CGGATAAAATGCTTGATGGTCGGAAGAGGCATAAATTCCGTCAGCCAGTTTAGTCTGACCA<br>TCTCATCTGTAACATCATTGGCAACGCTACCTTTGCCATGTTTCAGAAACAACTCTGGCGCA<br>TCGGGCTTCCCATACAATCGATAGATTGTCGCACCTGATTGCCCGACATTATCGCGAGCCC<br>ATTTATACCCATATAAATCAGCATCCATGTTGGAATTTAATCGCGGCCTCGAGCAAGACGTTT<br>CCCGTTGAATATGGCTCATAACACCCCTTGTATTACTGTTTATGTAAGCAGACAGTTTTATTG<br>TTCATGATGATATATTTTTATCTTGTGCAATGTAACATCAGAGATTTTGAGACACAACGTGGC<br>TTTCCCTGCAGGATTTCGGAGGCCTGCGTTATCCCCTGATTCTGTGGATAACCGTATTACC<br>GCCTTTGAGTGAGCTGATACCGCTCGCCGCAGCCGAACGCCGACTAGTGGATTTTACGGC<br>TAGCTCAGTCCTAGGTACAATGCTAGCGAATTCATTAAAGAGGAGAAAGGTACCCATGGCA<br>CGTACCCCGAGCCGTAGCAGCATTGGTAGCCTGCGTAGTCCGCATACCCATAAAGCAATTC<br>TGACCAGCACCATTGAAATCCTGAAAGAATGTGGTTATAGCGGTCTGAGCATTGAAAGCGT<br>TGCACGTCGTGCCGGTGCAAGCAAACCGACCATTTATCGTTGGTGGACCAATAAAGCAGC<br>ACTGATTGCCGAAGTGTATGAAAATGAAAGCGAACAGGTGCGTAAATTTCCGGATCTGGGT<br>AGCTTTAAAGCCGATCTGGATTTTCTGCTGCGTAATCTGTGGAAAGTTTGGCGTGAAACCA<br>TTTGTGGTGAAGCATTTCGTTGTGTTATTGCAGAAGCACAGCTGGACCCTGCAACCCTGAC<br>CCAGCTGAAAGATCAGTTTATGGAACGTCGTCGTGAGATGCCGAAAAAACTGGTTGAAAAT |

| | |
|---|---|
| | GCCATTAGCAATGGTGAACTGCCGAAAGATACCAATCGTGAACTGCTGCTGGATATGATTTT<br>TGGTTTTTGTTGGTATCGCCTGCTGACCGAACAGCTGACCGTTGAACAGGATATTGAAGAA<br>TTTACCTTCCTGCTAATTAATGGTGTTTGTCCGGGTACACAGCGTTAACTAGGGCCCATACC<br>CCCAATTATTGAAGGCCGCTAACGCGGCCTTTTTTTGTTTCTGGTCTGCCCGACGTACGGT<br>GAATCTGATTCGTTACCAATTGACATGATACGAAACGTACCGTATCGTTAAGGTTACTAGATT<br>AAAGAGGAGAAATACTAGATGGCAGTAAAGATTTCAGGAGTCCTGAAAGACGGCACAGGA<br>AAACCGGTACAGAACTGCACCATTCAGCTGAAAGCCAGACGTAACAGCACCACGGTGGTG<br>GTGAACACGGTGGGCTCAGAGAATCCGGATGAAGCCGGGCGTTACAGCATGGATGTGGA<br>GTACGGTCAGTACAGTGTCATCCTGCAGGTTGACGGTTTTCCACCATCGCACGCCGGGAC<br>CATCACCGTGTATGAAGATTCACAACCGGGGACGCTGAATGATTTTCTCTGTGCCATGACG<br>GAGGATGATGCCCGGCCGGAGGTGCTGCGTCGTCTTGAACTGATGGTGGAAGAGGTGGC<br>GCGTAACGCGTCCGTGGTGGCACAGAGTACGGCAGACGCGAAGAAATCAGCCGGCGATG<br>CCAGTGCATCAGCTGCTCAGGTCGCGGCCCTTGTGACTGATGCAACTGACTCAGCACGC<br>GCCGCCAGCACGTCCGCCGGACAGGCTGCATCGTCAGCTCAGGAAGCGTCCTCCGGCG<br>CAGAAGCGGCATCAGCAAAGGCCACTGAAGCGGAAAAAAGTGCCGCAGCCGCAGAGTCC<br>TCAAAAAACGCGGCGGCCACCAGTGCCGGTGCGGCGAAAACGTCAGAAACGAATGCTGC<br>AGCGTCACAACAATCAGCCGCCACGTCTGCCTCCACCGCGGCCACGAAAGCGTCAGAGG<br>CCGCCACTTCAGCACGAGATGCGGTGGCCTCAAAAGAGGCAGCAAAATCATCAGAAACGA<br>ACGCATCATCAAGTGCCGGTCGTGCAGCTTCCTCGGCAACGGCGGCAGAAAATTCTGCCA<br>GGGCGGCAAAAACGTCCGAGACGAATGCCAGGTCATCTGAAACAGCAGCGGAACGGAGC<br>GCCTCTGCCGCGGCAGACGCAAAAACAGCGGCGGCGGGGAGTGCGTCAACGGCATCCA<br>CGAAGGCGACAGAGGCTGCGGGAAGTGCGGTATCAGCATCGCAGAGCAAAAGTGCGGCA<br>GAAGCGGCGGCAATACGTGCAAAAAATTCGGCAAAACGTGCAGAAGATATAGCTTCAGCT<br>GTCGCGCTTGAGGATGCGGACACAACGAGAAAGGGGATAGTGCAGCTCAGCAGTGCAAC<br>CAACAGCACGTCTGAAACGCTTGCTGCAACGCCAAAGGCGGTTAAGGTGGTAATGGATGA<br>GACTAATCGTAAGGCACCTCTGGACAGTCCGGCACTGACCGGAACGCCAACAGCACCAA<br>CCGCGCTCAGGGGAACAAACAATACCCAGATTGCGAACACCGCTTTTGTACTGGCCGCGA<br>TTGCAGATGTTATCGACGCGTCACCTGACGCACTGAATACGCTGAATGAACTGGCCGCAG<br>CGCTCGGGAATGATCCAGATTTTGCTACCACCATGACTAACGCGCTTGCGGGTAAACAACC<br>GAAGAATGCGACACTGACGGCGCTGGCAGGGCTTTCCACGGCGAAAAATAAATTACCGTA<br>TTTTGCGGAAAATGATGCCGCCAGCCTGACTGAACTGACTCAGGTTGGCAGGGATATTCT<br>GGCAAAAAATTCCGTTGCAGATGTTCTTGAATACCTTGGGGCCGGTGAGAATTCGGGGAG<br>CGCTACAGACGTTATGATTCAGCTGGCGGCAAATGATGGCTTCAAATTCATCGGTCAGTGC<br>CCAGACATCTTGACCCTGCGTACTATCGAGCCGGAAAAAAACGGTCAGCGTATCACCTTAC<br>GTCAACATACGATTGGCACTGGCTTAGGCGGTGGCGTTTTCCGTGCAGTTCTGGACGGCA<br>CTGGCTATACCGATGACGACGGTGTGGTGATCAAAACCGCTGGGGGCAGCGTTTGGCTG<br>CGTGTCAACGCTGACAAAGTTAACCCGTTCATGTTCGGTGCAACCGGAGTAGCGGACGAC<br>ACCGCCGCCCTGCAAAAAATGCTGGAATGCGGTCGTGCGGCGGAACTGGGGACTAACGT<br>ATGGAAAGCAAGCAATCTGGAACTGAACAACAAATCTTGCTCTCTGTCCGGCAGTGGCCT<br>GCACGTTTCTCGTATTGAACAGATTTCCGGTGCAACCGGAGCATTGTTAACCATCACCCAA<br>GACTGTTCGCTGATTTACCTGTCCGATTGTGGCCTGTACGGCGATGGCATCACCGCAGGC<br>ACGAGCGGTGTTACTATGGAAACGGGTAATCCGGGTGGCGCTCCGTCTTACCCTTTCAATA<br>CCGCTCCGGACGTTCGTCGTGACCTGTACATCTCTAACGTGCACATCACGGGCTTCGACG<br>AGCTGGGTTTTGATTATCCGGAAACCAATTTCTCTGTTTCGACGCATGGCCTCTTCATCCGT<br>AACATCAAAAAAACGGGTGCAAAGATTGGTACTACGGACTTCACTTGGACTAACCTGCAAA<br>TTGATACTTGCGGTCAGGAATGTCTGGTGCTGGACGGTGCGGGTAACTGCCGTATTATTGG<br>TGCAAAACTGATTTGGGCAGGTAGCGAAAACGAAACGCCATACTCTGGCCTGCGTATTAGC<br>AACTCTCAAAATGTAAATATGACTGGCGTAGAGTTACAAGACTGCGCGTATGATGGTTTATA |

| | |
|---|---|
| | CATCAAGAACTCTACGGTTGCAATTTCAGGCTTAAACACCAATCGCAATAGCGCATCCTCTA<br>ATCTGTCCTACCATAACATGGTATTCGAAAATTCTATTGTAACTGTTGATGGTTATGTGTGTC<br>GTAACTACGCGGCGACTTCGCTGTACGACCTGAACAGCCAAGCAGGCAACGTCCGTTGCA<br>TCGGTAGCGACAGCACCGTTTTAATCAACGGCATCTACGAAAGCGAAGTCAATAGCGAGC<br>GCCTGATGGGTGATAACAACCTGATCCAGCCGTATAGTGGTGATCTGATCATTAACGGCCT<br>GAAAAATTACTACACCTATACTGGTAGCGTAAAAAACAACATTCCGACCTTCGACGGCGTTG<br>TTACTACGGCAACCTATGTGAGCGCACCGTCTATTCTGGGTCAGGGCAATATGCTCAAACT<br>GACCCAGTCTAATAAAGACAAACTGTTATTTAGCGATAAAGTTAGCCGTCATGGCTGTACCA<br>TCGGCTTAGTTCTGATTCCGTCCTTTACGGGCGCGACCACTATGACGGCGTTCACGCTGG<br>GTAGCGGTTACTCTCCATCCGGTAACTCCGCCGTGATGCAGTTCATTGTTAACAGTTCCGG<br>TGTACAAACCATTGCGATTTTATTATCCGGCGACGGTATTACCCAAACCCTGACCAGCGATC<br>TGACCACGGAACAAGCACTGGCGAGCGGTGGCGTGTATCATTTTGCAATGGGTTTTGCGC<br>CGGGTCGTTTATGGTGGAGCATTATCGATATTAACACGGGCAGGCGTATTCGTCGCGCCTA<br>CCGTCAGCCGGATCTGCACGCGGCGTTCAACTCTATCTTCAACTCCGGCACGTCGTCTATT<br>ACCGCATTTAGCGGGCCACTGGCGGGCGACATTGCTTGCGAAGGTGCAGGTAGCCATGTA<br>TACGTTGGCGGTTTTTCGTCGGAATCTGATTACGCGGCTAGCCGTATGTATGGCCTGTTCA<br>CTCCGGTCGATCTGGACAAGCAGTATAGCTTCCGTACCCTGAACGGTAACATTAACTATATT<br>GTGAGGCTTGCATAATGGCATTCAGAATGAGTGAACAACCACGGACCATAAAAATTTATAAT<br>CTGCTGGCCGGAACTAATGAATTTATTGGTGAAGGTGACGCATATATTCCGCCTCATACCGG<br>TCTGCCTGCAAACAGTACCGATATTGCACCGCCAGATATTCCGGCTGGCTTTGTGGCTGTT<br>TTCAACAGTGATGAGGCATCGTGGCATCTCGTTGAAGACCATCGGGGTAAAACCGTCTATG<br>ACGTGGCTTCCGGCGACGCGTTATTTATTTCTGAACTCGGTCCGTTACCGGAAAATTTTAC<br>CTGGTTATCGCCGGGAGGGGAATATCAGAAGTGGAACGGCACAGCCTGGGTGAAGGATA<br>CGGAAGCAGAAAAACTGTTCCGGATCCGGGAGGCGGAAGAAACAAAAAAAAGCCTGATG<br>CAGGTAGCCAGTGAGCATATTGCGCCGCTTCAGGATGCTGCAGATCTGGAAATTGCAACG<br>AAGGAAGAAACCTCGTTGCTGGAAGCCTGGAAGAAGTATCGGGTGTTGCTGAACCGTGTT<br>GATACATCAACTGCACCTGATATTGAGTGGCCTGCTGTCCCTGTTATGGAGTAATGACGCAT<br>CCTCACGATAATATCCGGGTAGGACGAACAATAAGGCCGCAAATCGCGGCCTTTTTTATTGA<br>TAACAAAAGGACAGTTTTCCCTTTGATATGTAACGGTGAACAGTTGTTCTACTTTTGTTTGTT<br>AGTCTTGATGCTTCACTGATAGATACAAGAGCCATAAGAACC |

**Supplementary Table 7. Plasmids used in this study**

| stx1 gene variants | | stx2 gene variants | | All stx gene variants | All isolates |
|---|---|---|---|---|---|
| stx1 guide 1 | stx1 guide 2 | stx2 guide 1 | stx2 guide 2 | 4 guides | |
| 100% | 99.34% | 100% | 100% | 100% | 100% |

**Supplementary Table 8. Percentage of *stx* gene variants and clinical isolates targeted by EB003 *in silico.***
